# A ppGpp-mediated brake on photosynthesis is required for acclimation to nitrogen limitation in Arabidopsis

**DOI:** 10.1101/2021.09.18.460674

**Authors:** Shanna Romand, Hela Abdelkefi, Cecile Lecampion, Mohamed Belaroussi, Melanie Dussenne, Brigitte Ksas, Sylvie Citerne, José Caius, Stefano D’Alessandro, Hatem Fakhfakh, Stefano Caffarri, Michel Havaux, Ben Field

## Abstract

Guanosine pentaphosphate and tetraphosphate (together referred to as ppGpp) are hyperphosphorylated nucleotides found in bacteria and the chloroplasts of plants and algae. In plants and algae artificial ppGpp accumulation can inhibit chloroplast gene expression, and influence photosynthesis, nutrient remobilisation, growth, and immunity. However, it is so far unknown whether ppGpp is required for abiotic stress acclimation in plants. Here, we demonstrate that ppGpp biosynthesis is necessary for acclimation to nitrogen starvation in Arabidopsis. We show that ppGpp is required for remodeling the photosynthetic electron transport chain to downregulate photosynthetic activity and for protection against oxidative stress. Furthermore, we demonstrate that ppGpp is required for coupling chloroplastic and nuclear gene expression during nitrogen starvation. Altogether, our work indicates that ppGpp is a pivotal regulator of chloroplast activity for stress acclimation in plants.

## Introduction

Plants cannot easily escape harsh environmental fluctuations, and so their survival hinges on facing each threat. To this end plants have developed intricate stress perception and response mechanisms (Devireddy et al., 2021), where the chloroplast is recognized as both a major signalling hub and a target for acclimation (Kleine et al., 2021). Likely candidates for regulating chloroplast stress signalling are the hyperphosphorylated nucleotides guanosine pentaphosphate and tetraphosphate (together referred to as ppGpp) that are synthesized from ATP and GDP/GTP by chloroplast localized enzymes of the RelA SpoT Homologue (RSH) family (Boniecka et al., 2017; Field, 2018). In bacteria, where ppGpp was originally discovered, a considerable body of work indicates that ppGpp and related nucleotides regulate growth rate and peak during stress to promote acclimation (Ronneau and Hallez, 2019; Bange et al., 2021; Anderson et al., 2021). In plants, ppGpp signalling is less well understood both at the physiological and mechanistic levels. ppGpp levels increase transiently in diverse plants in response to treatment with a range of abiotic stresses and stress related hormones (abscisic acid, jasmonate and ethylene)(Takahashi et al., 2004; Ihara et al., 2015). The accumulation of ppGpp is controlled by the ppGpp synthesis and hydrolysis activity of three families of chloroplast targeted RSH enzyme, RSH1, RSH2/3 and RSH4/CRSH (Atkinson et al., 2011; Ito et al., 2017; Avilan et al., 2019). In the angiosperm Arabidopsis, where ppGpp signalling is best characterized, RSH1 lacks ppGpp synthase activity and acts as the main ppGpp hydrolase (Sugliani et al., 2016), the closely related RSH2 and RSH3 act as the major ppGpp synthases during the day (Mizusawa et al., 2008; Maekawa et al., 2015; Sugliani et al., 2016), and the calcium activated RSH (CRSH) is responsible for ppGpp synthesis at night and in response to darkness (Ihara et al., 2015; Ono et al., 2020). RSH2 and RSH3 are bifunctional ppGpp synthase / hydrolase enzymes, and their hydrolase activity was recently shown to be necessary for constraining CRSH-mediated ppGpp production at night (Ono et al., 2020). ppGpp itself has been shown to act as an inhibitor of the expression of certain chloroplast genes *in vivo* (Yamburenko et al., 2015; Maekawa et al., 2015; Sugliani et al., 2016; Ono et al., 2020). Artificial accumulation of ppGpp in the chloroplast by the over-expression of RSH3 or bacterial ppGpp synthase domains has shown that ppGpp can inhibit photosynthesis in both land plants (Maekawa et al., 2015; Sugliani et al., 2016; Honoki et al., 2018; Harchouni et al., 2021) and algae (Avilan et al., 2021). Interestingly, the artificial accumulation of ppGpp also appears to protect plants against nitrogen deprivation (Maekawa et al., 2015; Honoki et al., 2018). In addition, a few studies have now demonstrated that ppGpp is required for different physiological processes; these include the regulation of plant growth and development (Sugliani et al., 2016; Ono et al., 2020), plant immunity (Abdelkefi et al., 2018), and photosynthesis under standard growth conditions (Sugliani et al., 2016). However, there are so far no demonstrations that ppGpp is required for abiotic stress acclimation in plants.

Here, we found that ppGpp biosynthesis by RSH2 and RSH3 is necessary for acclimation to nitrogen starvation in Arabidopsis. During nitrogen starvation ppGpp acts by remodeling the photosynthetic electron transport chain to downregulate photosynthetic activity, and by protecting against oxidative stress and tissue damage. Furthermore, we show that ppGpp couples chloroplastic and nuclear gene expression during nitrogen starvation. Overall, our work indicates that ppGpp is a pivotal regulator of chloroplast activity with a photoprotective role and that ppGpp signalling is required for the acclimation of plants to harsh environmental conditions.

## Results

### ppGpp is required for acclimation to nitrogen deprivation

To determine whether ppGpp plays a significant physiological role during nitrogen starvation we grew a series of *RSH* lines on a nitrogen limiting medium that imposes a progressive nitrogen starvation (Fig. 1A). The *RSH* lines were previously shown to accumulate lower (OX:RSH1, *RSH* quadruple mutant [*rsh_QM_*]) or higher (*rsh1-1*) amounts of ppGpp under standard growth conditions (Sugliani et al., 2016). Growth arrest occurred for all lines 9-10 days after sowing on nitrogen limiting media and was followed by the production of anthocyanins and loss of chlorophyll (Fig. 1A, S1). Strikingly, we observed that the low ppGpp lines OX:RSH1 and *rsh_QM_* showed significantly higher rates of cotyledon death than in the corresponding wild-type plants (Fig. 1A, B). Interestingly, between 16 and 22 days we also saw a greater rate of new leaf initiation in the ppGpp deficient lines (Fig. 1A).

**Figure 1.**
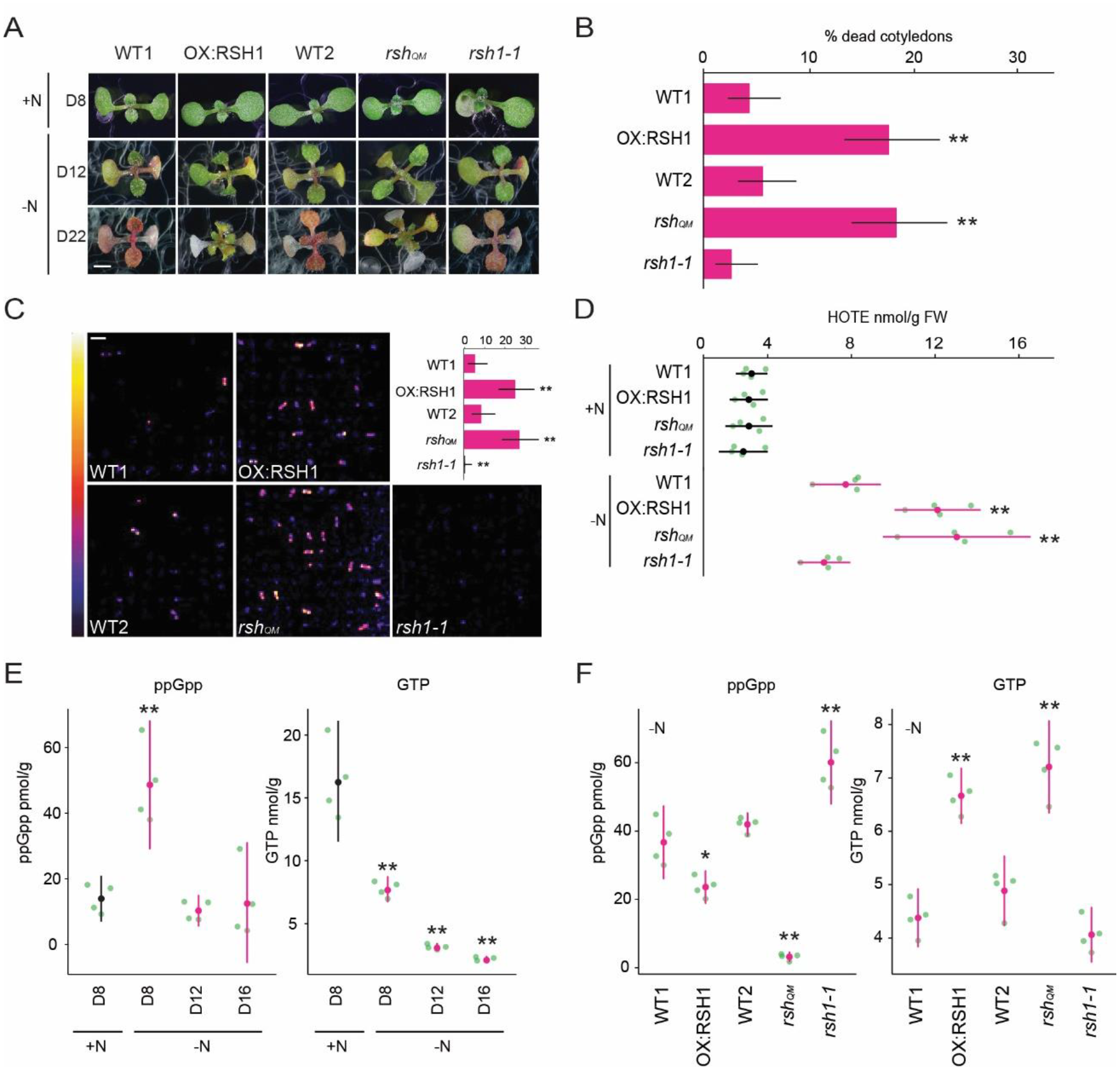
ppGpp is required for acclimation to nitrogen deprivation. **(A)** Images of seedlings grown on nitrogen replete (+N) or nitrogen limiting (-N) medium for 8 (D8), 12 (D12) and 22 (D22) days. Scale, 3 mm.WT1 (Col-0) and WT2 (*qrt1-2*) are the wildtypes for OXRSH1 and *rsh_QM_* / *rsh1-1* respectively. **(B)** Percentage of plants with dead cotyledons (completely white with collapsed tissue) for different genotypes grown on -N medium for 22 days. Three pooled biological replicates, mean +/-95% CI, n = 285 - 298 seedlings per genotype. **(C)** Bioluminescence emission from lipid peroxides in seedlings grown on -N medium for 16 days. Inset graph shows means +/- 95% CI, n = 100 seedlings. Scale, 1.1 cm. **(D)** Quantification of HOTEs from seedlings grown in +N for 8 days or -N for 12 days. Mean +/- 95% CI, n = 4 biological replicates. Concentrations of ppGpp and GTP in wildtype plants were determined **(E)** at the indicated time points during growth on +N and -N medium and **(F)** in different genotypes after 12 days of growth in -N medium (equivalent to 10 days in the experiment in panel E). Mean +/- 95% CI, n = 4 biological replicates. Statistical tests shown against respective wild type controls, * *P*<0.05, ** *P*<0.01.

We reasoned that the increased cotyledon death in the low ppGpp lines could be due to overproduction of reactive-oxygen species (ROS). We therefore imaged the autoluminescence of lipid peroxides, a signature of ROS accumulation (Fig. 1C). Under nitrogen-deprivation conditions strong autoluminescence was observed only in the OX:RSH1 and *rsh_QM_* low ppGpp lines, while *rsh1-1* plants showed almost no autoluminescence (Fig. 1C). Quantification of lipid peroxidation products (hydroxy-octadecatrienoic acids [HOTEs], the oxidation products of linolenic acid, the major fatty acid in Arabidopsis leaves) by HPLC further supported these findings: HOTEs increased in the wild-type in response to nitrogen deprivation, and this increase was significantly larger in the ppGpp deficient lines (Fig. 1D). These results indicate that nitrogen deprivation promotes ROS accumulation, and that ppGpp is required to prevent overaccumulation of ROS, oxidative stress and death of cotyledons under these conditions.

### ppGpp levels increase at an early stage of nitrogen deprivation

In bacteria and in plants ppGpp is known to peak in response to stress before stabilizing at lower levels (Varik et al., 2017; Takahashi et al., 2004a; Ihara et al., 2015). To determine the kinetics of changes in ppGpp concentration during nitrogen deprivation we quantified ppGpp levels in wild-type plants at different timepoints during growth on nitrogen limiting media. We observed a peak in ppGpp levels after 8 days on nitrogen limiting medium that was not observed in plants of the same age grown on nitrogen replete medium (Fig. 1E). The ppGpp concentration decreased and stabilised after 12 days on nitrogen limiting medium. In contrast, the level of GTP decreased throughout the timecourse. We next determined how (p)ppGpp levels were affected in each of the *RSH* lines during growth on nitrogen limiting medium (Fig. 1F). As anticipated we found that ppGpp levels in OX:RSH1 and *rsh_QM_* lines were lower than in wild-type plants, while they were higher in *rsh1-1* plants. Strikingly, we found that the lines with low ppGpp levels (OX:RSH1 and *rsh_QM_*) maintained higher levels of GTP than in the wild-type controls or *rsh1-1* (Fig. 1F), resulting in dramatic differences in the ppGpp/GTP ratio (Fig. S2). Therefore, nitrogen limitation leads to an early increase in ppGpp levels and is followed by a progressive diminution of the GTP pool that is dependent on ppGpp accumulation. Notably, this situation results in a sustained increase in the ppGpp/GTP ratio during nitrogen limitation.

### Nitrogen deprivation promotes a ppGpp-dependent drop in photosynthetic activity

Under standard growth conditions artificial production of ppGpp has been shown to downregulate photosynthetic activity in Arabidopsis (Sugliani et al., 2016; Maekawa et al., 2015). Defective downregulation of photosynthesis during nitrogen starvation could cause the over-accumulation of ROS and lipid peroxides observed in the low ppGpp lines due to oversaturation of photosynthesis in the presence of diminished metabolic sinks. We therefore measured different photosynthetic parameters in the wild type and *RSH* lines during growth on nitrogen limiting medium. A decrease in the maximum quantum yield (Fv/Fm) of photosystem II (PSII) was observed from 10 days in the cotyledons of wild-type seedlings (Fig. 2A-B, S3A). Remarkably, this decrease was almost completely suppressed in the low ppGpp *rsh_QM_* and OX:RSH1 lines, and was enhanced in the *rsh1-1* mutant (Fig. 2A, 2B, S3A). Analysis of *rsh2-1* and *rsh3-1* single mutants, as well as the complementation of an *rsh2-1 rsh3-1* double mutant with the genomic version of *RSH3*, indicates that the RSH3 enzyme is the major ppGpp synthase responsible for driving the Fv/Fm decrease under nitrogen starvation conditions (Fig. S3B). Likewise, the enhanced Fv/Fm decrease in the *rsh1-1* mutant and complementation of *rsh1-1* by over-expression of the genomic *RSH1* indicate that the ppGpp hydrolase RSH1 acts antagonistically to RSH3, probably by constraining ppGpp accumulation during nitrogen deprivation (Fig. 1F. 2A, 2B, S3C).

**Figure 2.**
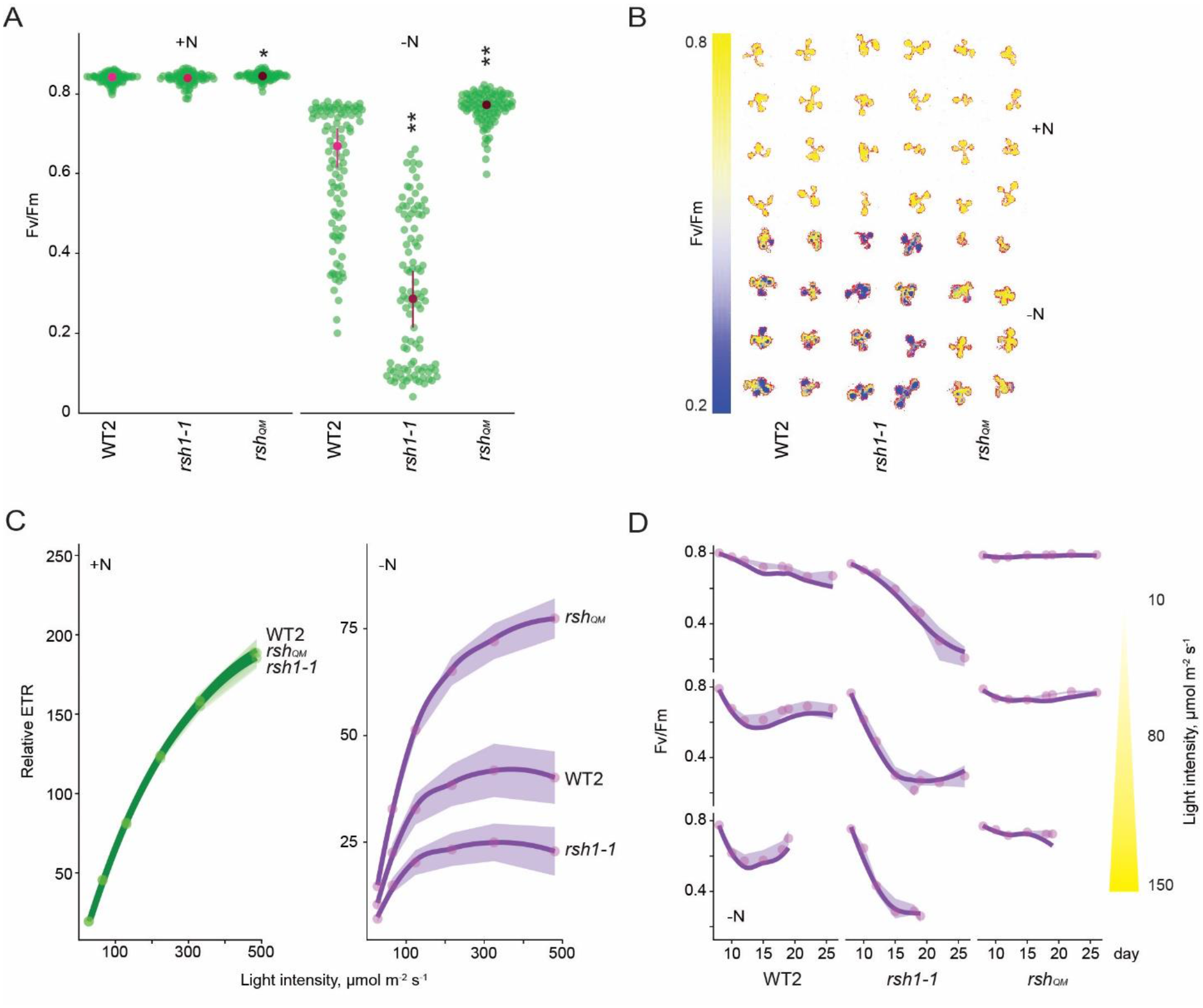
Nitrogen deprivation promotes a ppGpp-dependent drop in photosynthetic capacity. Seedlings were grown 8 days on nitrogen replete media (+N) or 12 days on nitrogen limiting (-N) media and **(A)** the maximal yield of PSII (Fv/Fm) measured by fluorescence imaging individual seedlings. **(B)** Fv/Fm images of whole seedlings grown on +N and -N media for 12 days. **(C)** Relative ETR measurements in different lines grown 8 days on +N media or 12 days on -N media. **(D)** Fv/Fm time-courses from seedlings grown on -N media and transferred to three different light intensities (photosynthetic photon flux density, 10, 80 and 150 µmol m^-2^s^-1^) after 6 days. Means +/- 95% CI, n = 95-100 seedlings. Statistical tests shown against respective wild type controls, * *P*<0.05, ** *P*<0.01.

While unlikely, it is possible that RSH enzymes could influence Fv/Fm via mechanisms that do not require ppGpp synthesis or hydrolysis. To exclude this possibility we conditionally overexpressed a heterologous chloroplast targeted ppGpp hydrolase from Drosophila (MESH)(Sugliani et al., 2016) (Fig. S3D). We observed that ppGpp depletion by MESH suppresses the Fv/Fm decrease in response to nitrogen limitation, confirming a specific role for ppGpp in this process.

Our *in vitro* experiments were carried out on seedlings and in the presence of exogenous sugar in the media. To test the robustness of our observations to different experimental set- ups we exposed mature plants grown in quartz sand to nitrogen starvation conditions (Fig S3E). We observed responses from the wild type, *rsh_QM_* and *rsh1-1* mutant that were consistent with those observed *in vitro*, confirming the robustness of the ppGpp-dependent decrease in Fv/Fm in response to nitrogen limitation.

Fv/Fm measurements provide information principally on PSII. To understand the state of the entire photosynthetic electron transport chain we next measured the relative rate of electron transport (ETR). We found that nitrogen deprivation led to a large decrease in ETR in wild type plants (Fig. 2C, S4A). The *RSH* lines showed similar ETRs to the wild type under nitrogen replete conditions. However, nitrogen deprivation led to large differences in the *RSH* lines compared to the wild type: a low ETR was observed in the *rsh1-1* mutant, and a substantially higher ETR in the low ppGpp *rsh_QM_* and OX:RSH1 lines. Induction of the MESH ppGpp hydrolase also prevented much of the decrease in ETR observed under nitrogen deprivation, again indicating that this phenomenon is dependent on the activity of RSH enzymes and ppGpp accumulation (Fig. S4B).

### The ppGpp-dependent decline in photosynthesis occurs even at low light fluences

The ppGpp-dependent drop in Fv/Fm during nitrogen deprivation is reminiscent of photoinhibition, a process whose rate is proportional to light intensity (Tyystjarvi and Aro, 1996). We therefore asked whether the ppGpp-dependent decrease in Fv/Fm is similarly dependent on the energetic pressure on the photosystems. Strikingly, under nitrogen deprivation the Fv/Fm of the wild type decreased to a similar minimum regardless of the light intensity (photosynthetic photon flux density; low light, 10 µmol m^-2^s^-1^; growth light, 80 µmol m^-2^s^-1^; and higher light, 150 µmol m^-2^s^-1^)(Fig. 2D). However, the rate of the Fv/Fm decrease was proportional to light intensity, showing a slower decrease to the minimum under low light than under normal light or high light. A similar phenomenon was observed in the *rsh1-1* line, except that the Fv/Fm dropped at a faster rate and to a considerably lower level. In contrast, in the *rsh_QM_* mutant we observed almost no decrease in Fv/Fm, regardless of the light intensity tested. The insensitivity of the *rsh_QM_* mutant indicates that the process is completely dependent on ppGpp biosynthesis. Furthermore, the response of the wild-type and even stronger response of the *rsh1-1* mutant at all light fluences indicate that ppGpp-initiated regulation of Fv/Fm is triggered in response to nitrogen deprivation rather than to changes in the excitation status of the photosynthetic electron transport chain (Fig. 2D).

### ppGpp is required for the timely degradation of photosynthetic proteins

In order to better understand the changes in the photosynthetic machinery that underlie the reduction of photosynthetic capacity during nitrogen starvation we analyzed the abundance of representative photosynthetic proteins in the wild type and OX:RSH1 (Fig. 3A). In the wild type, we observed a decrease in proteins representative of nearly all the major photosynthetic complexes analysed (PsbA, LHCB1, PsbO, PetA, PsaD, LHCA1, RBCL) and PTOX analyzed between 8 and 16 days of growth under nitrogen deprivation. The decrease in photosynthetic proteins was defective in OX:RSH1 plants, which showed a marked delay for the decrease in abundance of PsbA, PsbO, PetA, PTOX, PsaD and RBCL. This delay was especially visible after 12 days on nitrogen limiting medium. On the contrary, changes in LHCB1 and LHCA1, the nuclear encoded light harvesting proteins of PSII and PSI, were indistinguishable between wild type and OX:RSH1 plants.

**Figure 3.**
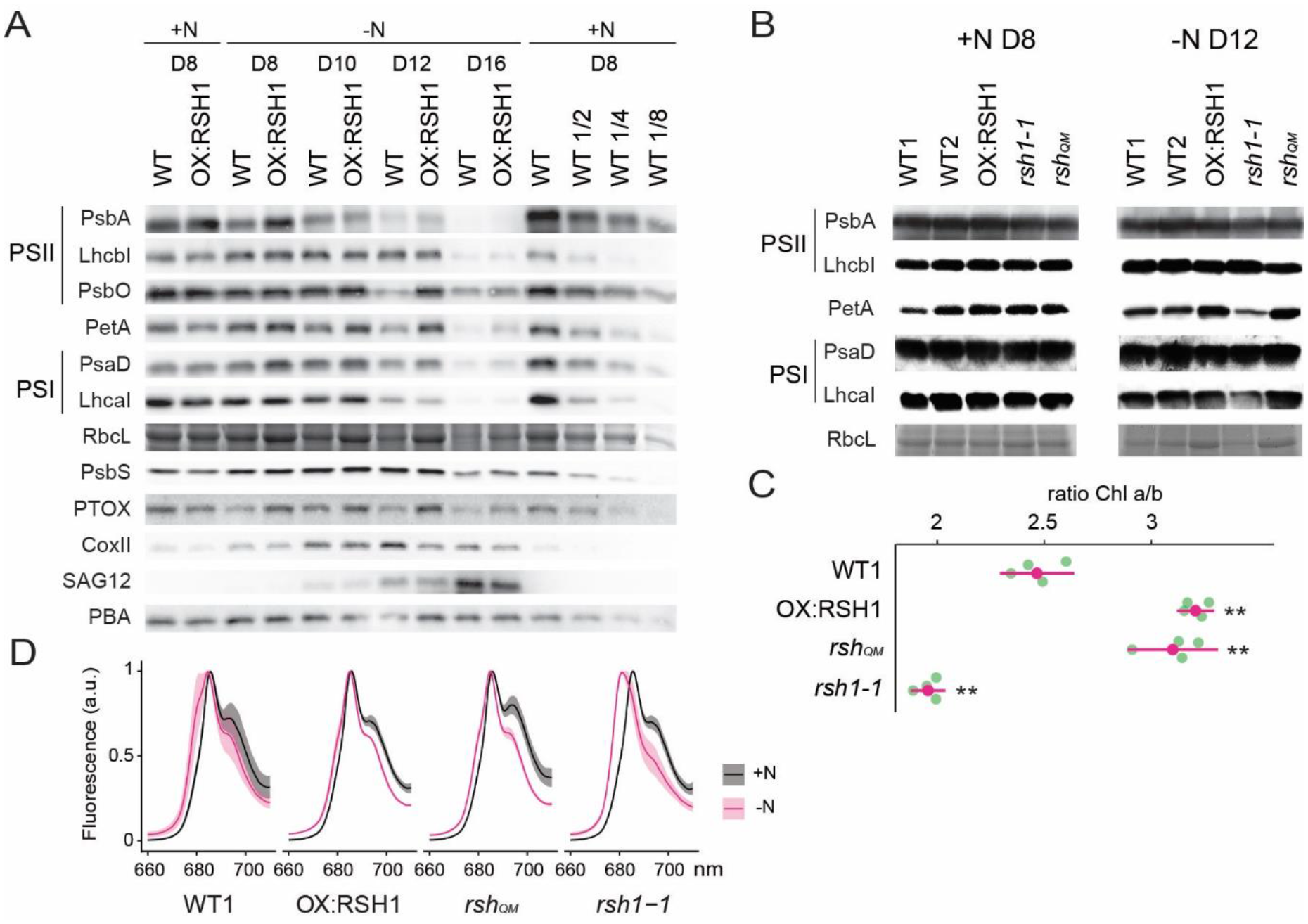
ppGpp-dependent alterations in the photosynthetic machinery during nitrogen deficiency. **(A)** Immunoblots showing evolution in abundance of the indicated proteins in seedlings grown in nitrogen replete (+N) or nitrogen limiting (-N) media for the indicated number of days. RbcL was revealed by Coomassie Brilliant Blue. Equal quantities of total proteins were loaded and PBA1, a subunit of the proteasome, was used as a protein normalisation control. **(B)** Immunoblots showing the abundance of the indicated proteins in purified thylakoid membranes from seedlings grown in +N for 8 days or -N for 12 days. RbcL was revealed by Coomassie Brilliant Blue staining. Equal quantities of total chlorophyll were loaded. **(C)** Chlorophyll a/b ratios in extracts from seedlings subjected to -N for 12 days. Means +/- 95% CI, data from four biological replicates. **(D)** Emission spectrum of chlorophyll fluorescence at 77°K between 660 and 720 nm. Measurements made seedlings grown in +N for 8 days or -N for 12 days. Means ± 95% CI; data from four biological replicates. Statistical tests, ** *P<*0.01.

In contrast to the downregulation of many components of the photosynthetic chain during N deprivation, mitochondrial activity appeared to increase. The mitochondrial marker protein COXII, a subunit of the mitochondrial complex IV, increased to reach a maximum on day 12 of nitrogen deprivation. Interestingly, the COXII maximum was higher in the wild type than in OX:RSH1 plants suggesting increased ppGpp levels could also positively influence mitochondrial activity. PsbS, a marker of non-photochemical quenching, and SAG12, a marker of senescence, showed similar patterns of induction in WT and OX:RSH1 plants.

We next extended our analysis from OX:RSH1 to the other RSH lines to confirm that ppGpp is responsible for the observed changes in photosynthetic proteins during nitrogen deprivation (Fig. 3B). In this experiment we normalised to total chlorophyll to exclude possible artefacts linked to normalization on total protein, yet we obtained broadly similar results. PetA and RBCL, the proteins with the greatest differences in OX:RSH1 during the timecourse, were more abundant in the low ppGpp lines OX:RSH1 and *rsh_QM_* at day 12 under nitrogen deprivation. The same proteins were also less abundant than the wild type in the high ppGpp line *rsh1-1*. Interestingly, PsbA accumulation did not appear different between the lines, contrasting with results from the OX:RSH1 time course. However, this difference is likely linked to the chlorophyll normalization because LHCB1 levels were higher in the wild type and *rsh1-1* mutant, indicating a similar direction of change for the ratio of PSII core to antenna. Finally, to unequivocally link these changes to ppGpp we analyzed representative protein levels in induced MESH ppGpp hydrolase lines, which again showed the ppGpp is required for diminution of PetA and RBCL levels during nitrogen deprivation (Fig. S5). Altogether our immunoblotting experiments indicate the ppGpp is required for the timely reduction in abundance of key photosynthetic proteins during nitrogen starvation, and that proteins such as PsbO, PetA and RBCL appear to be preferentially affected.

### Nitrogen starvation promotes ppGpp-dependent energetic uncoupling of PSII antenna

The ppGpp dependent drop in Fv/Fm and drop in the PsbA / LHCB1 ratio suggest that ppGpp specifically remodels the structure of the PSII complex during nitrogen deprivation. Supporting this idea, we also found a ppGpp dependent drop in the Chl a/b ratio (Fig. 3C), indicating a substantial decrease in PSII RC, which lacks Chl b, to Chl b rich PSII antenna. An increase in relative PSII antenna abundance might be accompanied by the energetic uncoupling from the PSII RC. Low temperature chlorophyll fluorescence spectra confirmed this hypothesis. During nitrogen deprivation there was a shift of the PSII antenna emission peak towards lower wavelengths in the wild type, indicating an energetic decoupling from PSII RC (Crepin et al., 2016)(Fig. 3D, S6). This shift is mainly ppGpp dependent because it was smaller in OX:RSH1 and *rsh_QM_*, and much stronger in *rsh1-1*. Therefore, multiple approaches show that ppGpp is necessary for PSII remodeling and decreasing the energetic pressure on the photosystems during nitrogen starvation.

### ppGpp plays a major role during acclimation to nitrogen deprivation

Our results show that defects in the capacity of plants to accumulate ppGpp lead to aberrant photosynthesis and stress phenotypes under nitrogen deprivation. To determine the extent and specificity of the impact of ppGpp on cellular processes we analyzed global nuclear and organellar transcript abundance in wild type and OX:RSH1 plants.

Nitrogen deprivation caused a massive alteration in transcript levels in both the wild type and OXRSH1, significantly affecting the accumulation of about 15,000 transcripts (Fig. 4A, Table S1). For the wild type there was substantial overlap (80%) with differentially accumulating transcripts recently identified in plants grown under chronic nitrogen limitation for 60 days (Luo et al., 2020). Strikingly, we observed large increases in *RSH2* and *RSH3* transcript levels in response to nitrogen deprivation (Fig 4B). These were accompanied by a decrease in *RSH1* transcript levels, suggesting a coordinated upregulation of ppGpp biosynthetic capacity. The expression profile of OX:RSH1 plants showed clear differences to the wild type, especially under nitrogen deprivation. Under control conditions OX:RSH1 plants had a very similar gene expression profile to the wild type, with the differential accumulation of only 150 transcripts (Fig. 4A, Table S1). However, under nitrogen deprivation we observed the differential accumulation of 2700 transcripts in OX:RSH1. The greater deregulation of the OX:RSH1 transcript profile under nitrogen deprivation indicates that ppGpp plays a major and specific role in the acclimation response.

**Figure 4.**
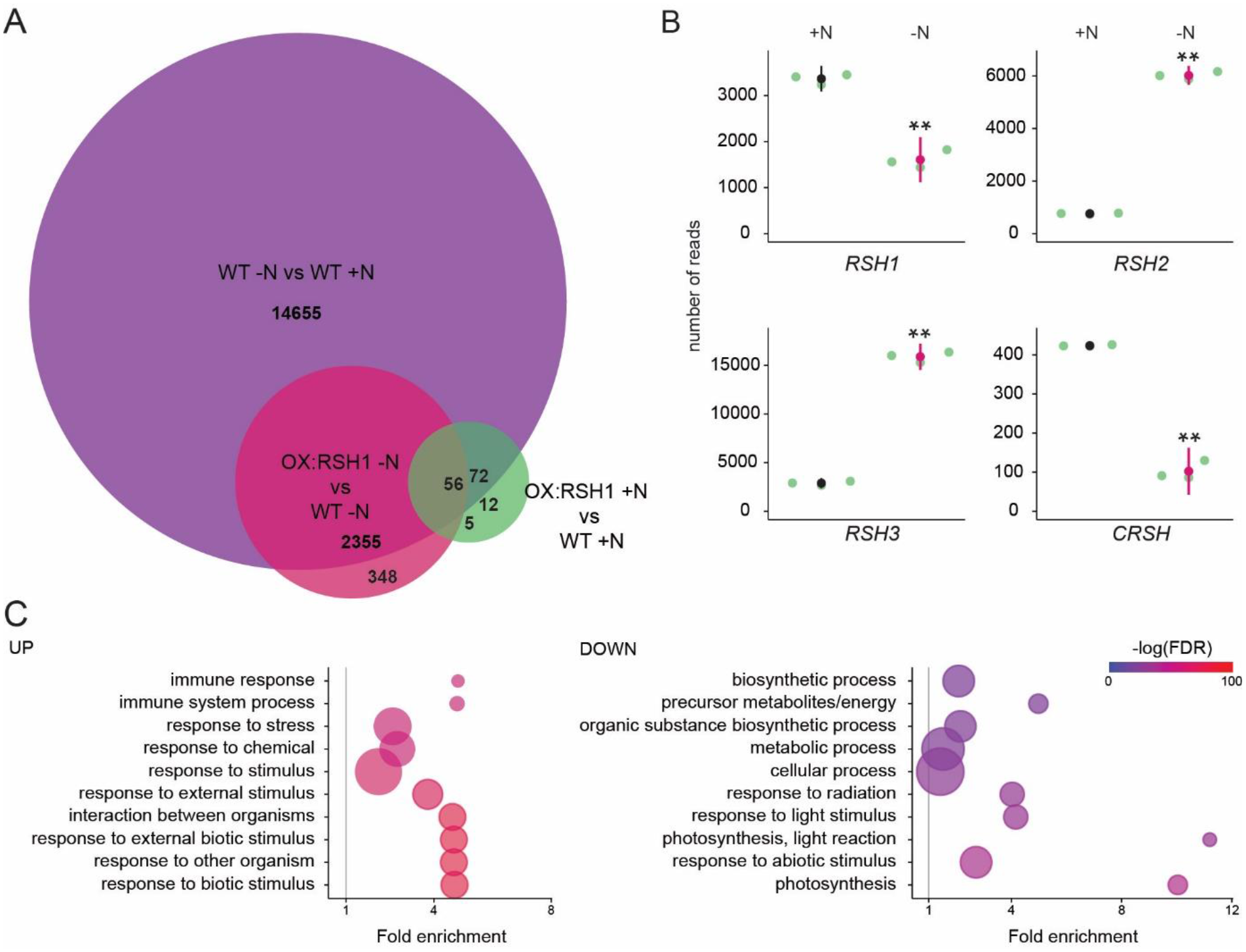
ppGpp plays a major role during acclimation to nitrogen deprivation. RNA-seq experiments were performed on WT and OX:RSH1 seedlings grown 8 days on nitrogen replete media (+N) or 12 days on nitrogen limiting (-N) media. **(A)** Venn diagram for transcripts showing differential accumulation for each of three comparisons. **(B)** RNA-seq transcript levels for the four *RSH* genes in the WT, +N vs -N. Mean +/ 95% CI, n=3 biological repeats. ** *P<*0.01. **(C)** Enriched gene ontology terms among significantly up- and down-regulated transcripts in OX:RSH1 vs WT under -N. The ten most significant terms are shown, point size is proportional to gene number. FDR, false discovery rate.

Analysis of enrichment for gene ontology terms corroborated our findings that low ppGpp plants are unable to properly acclimate to nitrogen deprivation. OX:RSH1 plants accumulated more transcripts associated with stress, including oxidative stress than the wild type (Fig. 4C, Table S2). We also observed a significant decrease in the abundance of transcripts for genes involved in photosynthesis and chloroplast activity. Notably, these downregulated genes were all nucleus encoded (Fig. 4C).

### ppGpp is required for global downregulation of chloroplast gene expression

We next turned to the effects of nitrogen deprivation on organellar gene expression. Nitrogen deprivation leads to a strong decrease in the levels of all chloroplast transcripts except one in the wild type (Fig. 5A, Table S3). The OX:RSH1 mutant showed minor differences to the wild type under normal growth conditions. However, under nitrogen starvation the vast majority of chloroplast transcripts (120 of 133 transcripts analyzed) were present at significantly higher levels in OX:RSH1 than in the wild type, indicating that ppGpp is required for globally reducing chloroplast gene expression. In contrast to the monodirectional response of the chloroplast to nitrogen deprivation, we found that mitochondrial transcripts showed diverse responses in the wild type (Fig. 5B). Significantly, mitochondrial transcript levels in the OX:RSH1 line were highly similar to the wild type under control and nitrogen deprivation conditions. Taken together, these results indicate that ppGpp is required for the downregulation of chloroplast gene expression during nitrogen starvation and demonstrate that ppGpp acts specifically within the chloroplast.

**Figure 5.**
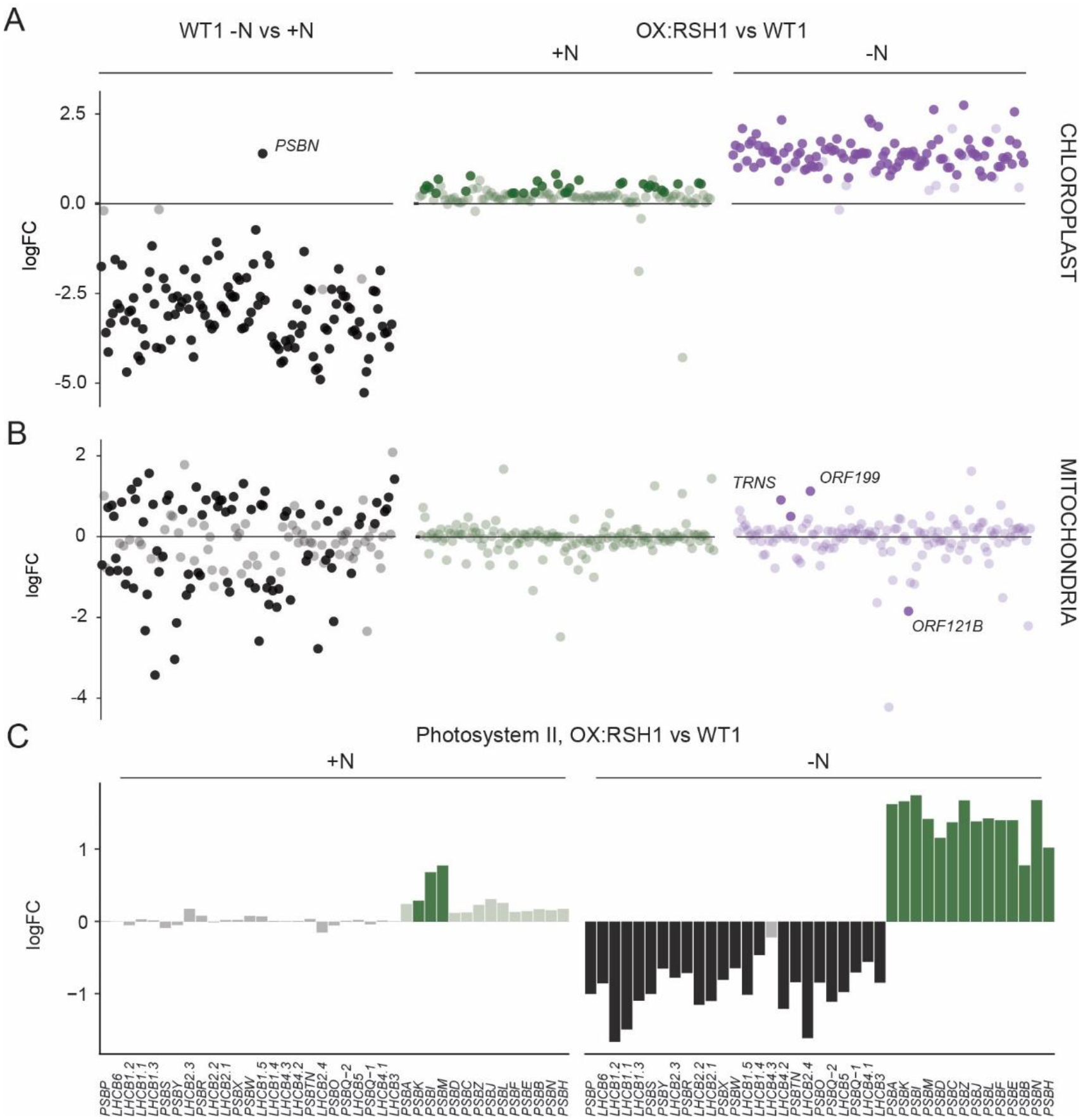
ppGpp is required for the downregulation of chloroplast gene expression during nitrogen deficiency. **(A)** The differential expression of chloroplast transcripts ordered along the chloroplast genome. **(B)** The differential expression of mitochondrial transcripts ordered along the mitochondrial genome. **(C)** Relative transcript levels in OX:RSH1 versus the wild type control for nuclear (black) and chloroplast (green) genes encoding subunits of the photosystem II complex. Solid colors indicate significantly different changes in expression (*P*<0.05), transparent colors indicate non-significant changes.

### ppGpp is required for coordinating chloroplast and nuclear gene expression

Our gene expression analysis showed the ppGpp depletion during nitrogen starvation appeared to have opposite effects on transcript abundance for nuclear encoded chloroplast genes and chloroplast encoded genes. Many chloroplast protein complexes contain subunits encoded by both the nuclear and chloroplast genomes. Notably, in the wild-type chloroplast protein complexes showed coordinate downregulation of both nuclear and chloroplast transcripts in response to nitrogen deprivation (Fig. S7). However, ppGpp depletion caused widespread mis-regulation of transcript abundance for complex subunits (Fig 5C, S8). For the PSII complex there was little difference between OX:RSH1 and the wild type under nitrogen replete conditions. However, in OX:RSH1 nitrogen deprivation caused the large scale uncoupling of the coordination of gene expression between the nuclear and chloroplast genomes: all but one of the transcripts for nuclear-encoded subunits of PSII were downregulated, while all the transcripts for chloroplast-encoded subunits of PSII were upregulated (Fig. 5C). Similar genome uncoupling was observed in OX:RSH1 for transcripts encoding subunits of the chloroplastic CytB6f, PSI, ATP synthase, NDH, TIC, transcription and translation complexes (Fig. S8). These results indicate first that nitrogen deprivation promotes the coordinated downregulation of genes encoding chloroplastic proteins in the nuclear and chloroplastic genomes, and second that ppGpp plays a pivotal role in coupling gene expression between nuclear and chloroplast genomes under nitrogen limitation.

## Discussion

Nitrogen deprivation has long been known to cause a drop in photosynthetic capacity in plants (Terashima and Evans, 1988; Nunes et al., 1993; Verhoeven et al., 1997; Lu and Zhang, 2000; Garai and Tripathy, 2018). Here we show that ppGpp signalling plays a major role in this process. We demonstrate that nitrogen starvation leads to an early and transient increase in ppGpp levels (Fig. 1E), and that the capacity to synthesize and accumulate ppGpp is then required for protecting plants against excess ROS accumulation, tissue damage and stress (Fig. 1A-D, 4C). We show that ppGpp is likely to mediate the acclimation response by affecting the ppGpp/GTP ratio (Fig. S2), promoting the downregulation of photosynthetic capacity (Fig. 2A-D, 4C, 5A) and remodelling the photosynthetic machinery (Fig. 3A-D, 5C). Finally, we show that ppGpp may function by specifically downregulating chloroplast transcript abundance during nitrogen deprivation to maintain an equilibrium between chloroplast and nuclear gene expression (Fig. 5A-C, S7). ppGpp was initially discovered thanks to the identification of “relaxed” bacterial mutants that fail to acclimate to amino acid starvation (Stent and Brenner, 1961; Cashel and Gallant, 1969). Our results here therefore indicate that the fundamental involvement of ppGpp signalling in acclimation to nitrogen deprivation is likely to have been maintained despite the large evolutionary distance separating bacteria from plants that includes major shifts including the domestication of the cyanobacterial ancestor of the chloroplast, and the different regulatory logic and signalling networks of photosynthetic eukaryotes.

### The physiological relevance of ppGpp signalling in plants

Depletion or removal of ppGpp is usually necessary for directly establishing the implication of ppGpp in a physiological process. Up to now a handful of studies in plants have directly implicated ppGpp physiological processes by showing a requirement for ppGpp synthesis (Sugliani et al., 2016; Abdelkefi et al., 2018; Honoki et al., 2018). We now add to these studies by using multiple approaches, including three different ppGpp depletion methods, to demonstrate that ppGpp is required for acclimation to nitrogen deprivation. Previous work hinted that ppGpp may play such a role by showing that plants ectopically over-accumulating ppGpp appear able to better withstand transfer to media lacking nitrogen (Maekawa et al., 2015; Honoki et al., 2018). Notably, Honoki et al. (2018) also used a similar *rsh2 rsh3* double mutant in their experiments though, apart from a delay in Rubisco degradation, did not observe clear-cut differences from the wild type in terms of photosynthetic activity after transfer to media lacking nitrogen. The reasons for this are not clear, but it could be related to differences in experimental set up such as the timepoints for analysis, or in the imposition of nitrogen starvation.

Our findings further reinforce the evidence that RSH enzymes function in the chloroplast. GFP fusion experiments have shown that RSH enzymes are localised to the chloroplasts (Maekawa et al., 2015), and the pleiotropic phenotypes of RSH3 overexpressing plants can be suppressed by chloroplast expression of a ppGpp hydrolase (Sugliani et al., 2016). However, some proteins show low levels of dual targeting that are not detectable using full length fluorescent protein fusions (Sharma et al., 2019). Coverage of the full mitochondrial transcriptome here shows that overexpression of RSH1 has almost no effect on the large-scale changes in mitochondrial transcript abundance that occur in response to nitrogen deprivation (Fig. 5B). These results therefore confirm that RSH enzymes operate within the chloroplast.

### New insights into ppGpp signalling in planta

The kinetics of changes in ppGpp concentration during nitrogen deprivation provide new information on how ppGpp acts *in planta*. We observed a transient peak in ppGpp levels after eight days of growth on nitrogen limiting medium (Fig. 1E). Similar transient peaks of ppGpp accumulation were also observed in plants subjected to different stresses (Takahashi et al., 2004; Ihara et al., 2015; Ono et al., 2020), and are observed in bacterial cells (Varik et al., 2017). Notably, the peak in ppGpp that we observed in response to nitrogen deprivation occurs prior to major changes in Fv/Fm (Fig. 2D), protein levels (Fig. 3A), or anthocyanin accumulation (Fig. S1). These factors, along with the progressive nature of the nitrogen starvation suggest that a signalling mechanism activates ppGpp biosynthesis relatively soon after perception of nitrogen supply limitation. The upregulation of *RSH2* and *RSH3* expression (Fig. 4B) indicates that there is at least a transcriptional component to this response. However, additional layers of regulation are likely, including for example the allosteric activation of RSH enzymes by protein-protein or protein small molecule interactions as found for bacterial RSH (Ronneau and Hallez, 2019; Irving and Corrigan, 2018). After the initial peak the ppGpp concentration then drops to levels not very different to those observed in plants grown in nitrogen replete conditions (Fig. 1E). Furthermore, OX:RSH1 plants only show slightly lower levels of ppGpp than the wild type despite showing very similar phenotypes to the *rsh_QM_* mutant (Fig. 1F). Together these observations indicate that ppGpp continues to exert an effect after peaking, suggesting the activation of irreversible processes or increased sensitivity to the presence of ppGpp. The second of these possibilities is supported by the phenotype of the *rsh1-1* mutant which displays much stronger photosynthesis phenotypes than wild type plants despite a relatively modest increase in ppGpp levels (Fig. 1F, 2A-D, 3A-D).

GTP is a key nucleotide in ppGpp signalling because it is both the substrate for ppGpp biosynthesis and its biosynthetic pathway a potential target of ppGpp (Field, 2018). We observed a striking decrease in GTP levels during nitrogen deprivation (Fig. 1E). Remarkably GTP levels were closely and inversely correlated to ppGpp levels in the wild type and the *RSH* lines (Fig. 1E, F). GTP is present at levels almost three orders of magnitude higher than ppGpp, so the relationship between ppGpp and GTP cannot be simply explained by the consumption of GTP for ppGpp biosynthesis. Therefore, ppGpp accumulation drives the depletion of the cellular GTP pools by other means, which could for example include the inhibition of enzymes involved in chloroplastic GTP biosynthesis such as guanylate kinase (Nomura et al., 2014). Even more remarkable is that the ppGpp-driven GTP depletion occurs specifically under nitrogen deprivation because accumulation of ppGpp to high levels in OX:RSH3 plants does not affect the GTP pool under normal growth conditions (Bartoli et al., 2020). ppGpp can act as a competitive inhibitor of GTP dependent enzymes due to its structural resemblance to GTP (Pausch et al., 2018). An increase in the ppGpp / GTP ratio will therefore significantly augment the competitivity of ppGpp and may explain the different response of the RSH lines to nitrogen deprivation (Fig. S2). We therefore propose that an increase in the ppGpp / GTP ratio during nitrogen limitation is likely to explain how ppGpp maintains its influence despite decreasing in absolute amounts.

### The impact of ppGpp on photosynthesis under nitrogen deprivation

Our work also reveals the wide-ranging physiological effects of ppGpp on photosynthetic activity during nitrogen deprivation. These effects are reminiscent of the photosynthetic changes that occur during senescence (Krieger-Liszkay et al., 2019), and are remarkably similar to those found using the ectopic over-accumulation of ppGpp in Arabidopsis, moss and algae (Maekawa et al., 2015; Sugliani et al., 2016; Imamura et al., 2018; Honoki et al., 2018; Avilan et al., 2021; Harchouni et al., 2021). However, compared to these studies our more detailed analysis here highlights first the global effect of ppGpp on chloroplast transcript abundance (Fig. 5A), and second the strong effect of ppGpp on certain photosynthetic proteins such as PsbO from PSII, PetA from Cyt b_6_f, and RbcL (Fig. 3A, B). ppGpp may control the abundance of these proteins via the downregulation of chloroplast transcript abundance (Fig. 5A). However, due to the selective nature of protein disappearance, it is likely that more active processes are involved. This idea is supported by previous work showing the high stability of Cyt b_6_f in tobacco (Hojka et al., 2014), the selective loss of Cyt b_6_f during natural senescence (Krieger-Liszkay et al., 2019; Roberts et al., 1987), and the protease-mediated degradation mechanism for Cyt b_6_f and Rubisco that operates under nitrogen limitation in Chlamydomonas (Majeran et al., 2000; Wei et al., 2014). PSII architecture was also modified in a ppGpp dependent manner with an increase in the ratio of PSII antenna to RC at both protein (Fig. 3A-D) and transcript levels (Fig. 5C). This was also accompanied by a controlled decrease in Fv/Fm, which is typically characterized as photoinhibition and occurs when damage to the PSII RC exceeds repair capacity. While commonly used as a sign of plant stress, controlled photoinhibition is increasingly recognized as a circuit breaker that can protect PSI against overexcitation damage (Tikkanen et al., 2014). The PSII RC subunit PsbA suffers photodamage at all light intensities (Tyystjarvi and Aro, 1996). However, under normal light fluences PSII function is unaffected thanks to the PSII repair pathway that rapidly replaces damaged PsbA with newly synthesised protein. Before this study, the drop in Fv/Fm observed during nitrogen starvation might have been explained by overexcitation of photosynthetic electron transport chain or reduced PSII repair efficiency due to the absence of building blocks for protein synthesis. However, our data indicate that ppGpp signalling actively controls this process by blocking PSII RC repair so that photoinactivation gains the upper hand and Fv/Fm drops. How does ppGpp inhibit PSII repair? As for the selective loss of certain photosynthetic proteins we cannot say at this stage whether this is only a consequence of transcriptional downregulation or may also involve other processes such as an inhibition of translation due to the altered ppGpp / GTP ratio.

### ppGpp mediates a novel stress inducible photoprotection pathway

We show that ROS increases in the wild type in response to nitrogen deprivation, and that defects in ppGpp biosynthesis lead to greater increases in ROS and cell death (Fig. 1B, D). The controlled ROS increase in the wild type is in agreement with a very recent study also showing ROS accumulation in the leaves of nitrogen starved plants (Safi et al., 2021). Chloroplasts are the major source of ROS in photosynthetic plant leaves (Rogers and Munné-Bosch, 2016; Domínguez and Cejudo, 2021). The role for ppGpp signalling in the prevention of ROS over-accumulation and tissue death during nitrogen limitation (Fig. 1A-D) is therefore very likely related to its role in downregulating photosynthetic activity (Fig. 2, S3 & S4). In addition to downregulating photosynthetic activity, ppGpp signalling triggers the controlled photoinhibition of PSII. Altogether, our data on the role of ppGpp in the downregulation of photosynthetic capacity, the activation of controlled photoinhibition, and the prevention of ROS accumulation strongly suggest that ppGpp signalling functions as a photoprotective mechanism in plants. This mechanism is triggered in response to nutrient limitation rather than light as for the known inducible photoprotective pathways (Pinnola and Bassi, 2018; Malnoë, 2018). The wide range of stresses known to induce ppGpp accumulation (Takahashi et al., 2004) and the association between stress and overexcitation of the photosynthetic electron transport chain (Bechtold and Field, 2018), together suggest that ppGpp signalling may allow plants to anticipate and acclimate to harsh environmental conditions that would otherwise lead to overexcitation of the photosynthetic machinery and oxidative stress.

## Materials and Methods

### Plant materials and growth conditions

*Arabidopsis thaliana* mutant lines, overexpressors and complementation lines and respective wild-type controls are listed in Table S5. In each experiment the seeds for each line were derived from the same batch of plants grown together. For in vitro growth, seeds were surface sterilized with 75% ethanol, rinsed with 100% ethanol, dried and placed in a ten by ten grid pattern on square culture dishes containing 45 ml of nitrogen replete (+N) (0.5x Murashige and Skoog salts [Caisson Labs], 1% sucrose, 0.5 g/L MES and 0.4% Phytagel [Merck Sigma-Aldrich], adjusted to pH 5.7 with KOH) or nitrogen limiting (-N) (+N medium diluted 1/25 in 0.5X Murashige and Skoog medium without nitrogen [Caisson Labs], 0.5 g/L MES and 0.4% Phytagel [Merck Sigma-Aldrich], adjusted to pH 5.7 with KOH) growth medium (for detailed medium composition see Table S6). Plates were placed at 4°C for 2 days in the darkness, and then then transferred to a 16 h light (at 22°C) / 8 h darkness (at 19.5°) photoperiod with 80 μmol m^-2^ s^-1^ lighting. For growth in quartz sand, seeds were germinated in soil, and then transferred into pots containing quartz sand at 7 days after germination. The plants were then grown under a 8 h light (at 22°C)/16 h dark (at 19.5°) photoperiod at 18 / 22°C with 110 µmol m^-2^ s^-1^ lighting. Plants were treated weekly with a complete nutrient solution. After 50 days the pots were rinsed, and then treated weekly with 0.5X Murashige and Skoog medium with or without nitrogen for three weeks.

### Creation of *rsh1-1* complementation lines

The 9.1 kb genomic *RSH1* sequence including the 3′ untranslated region, 5′ untranslated region, and 3 kb of upstream sequence containing the promoter was amplified from Arabidopsis genomic DNA using Phusion polymerase (New England Biolabs) using flanking primers and then the primers RSH1-F (TCCGTCTTGTCTGAATCAGCT) and RSH1-R (TTTCTAGATTTACTTTGGTTTTGTCCA) with attB1 / attB2 adapters. The resulting PCR product was then introduced by Invitrogen BP Gateway recombination (Thermofisher) into pDONR207. The entry clone was confirmed by sequencing and recombined by Invitrogen LR Gateway recombination into the binary vector pGWB410 (Nakagawa et al., 2007) which carries a kanamycin resistance cassette for selection in plants. The resulting construct was transferred into Agrobacterium (strain GV3101) and used to transform *rsh1-1* plants by floral dipping.

### Autoluminescence imaging and quantification of HOTEs

Peroxidated lipids were visualized by autoluminescence imaging as described previously (Birtic et al., 2011) and similar results were observed in four biological replicates. The intensity of the leaf autoluminescence signal is proportional to the amount of lipid peroxides present in the sample, the slow spontaneous decomposition of which produces luminescent species. For quantification of HOTEs, lipids were extracted from about 300 mg of seedlings in methanol / chloroform and analyzed by HPLC-UV as detailed elsewhere (Montillet et al., 2004; Shumbe et al., 2017).

### Nucleotide triphosphate quantification

Nucleotides were extracted from about 150 mg of seedlings, and quantified by HPLC-MS/MS using stable isotope labelled ppGpp and GTP standards as described previously (Bartoli et al., 2020).

### Chlorophyll fluorescence measurements

Plants were dark adapted for 20 minutes and chlorophyll fluorescence was measured in a Fluorcam FC 800-O imaging fluorometer (Photon System Instruments). PSII maximum quantum yield (Fv/Fm) was calculated as (Fm − F0)/Fm. For relative electron transfer rate (ETR) measurements plants were exposed to 2 minutes periods of increasing actinic light intensity, with Fv’/Fm’ measurements at the end of each period. Relative ETR was calculated as Fv’/Fm’ X light intensity (photosynthetic photon flux density). Photosynthetic photon flux density was measured using a Universal Light Meter 500 (ULM-500, Walz). All experiments on photosynthetic parameters were repeated two to five times with similar results. Steady state 77K chlorophyll fluorescence measurements were obtained from frozen seedling powder suspended in 85% (w/v) glycerol, 10 mM HEPES, pH 7.5 as described previously (Galka et al., 2012).

### Protein extraction and immunoblotting

Protein extraction and immunoblotting were performed as described previously (Sugliani et al., 2016), with the exception of a protein precipitation step with 20% TCA after protein extraction. The precipitation step was necessary to remove compounds in nitrogen-limited seedlings that interfered with the bicinchoninic acid assay for determining protein concentration. The following primary antibodies were used against CoxII (Agrisera; polyclonal; ref. AS04 053A), LHCA1 (Agrisera; polyclonal, ref. AS01 005), LHCB1 (Agrisera; polyclonal, ref. AS01 004), Pba1 (Abcam; polyclonal; ref. ab98861), PetA (Agrisera; polyclonal, ref. AS08 306), PsaD (Agrisera; polyclonal; ref. AS04 046), PsbA (Agrisera; polyclonal, ref. AS05 084), PsbO (Agrisera; polyclonal; ref. AS05 092), PTOX (Uniplastomic, Biviers, France; kindly provided by Xenie Johnson), SAG12 (Agrisera; polyclonal; ref. AS14 2771).

For the analysis of thylakoidal proteins, seedlings were harvested and then ground in liquid nitrogen. Proteins were extracted in an extraction buffer adapted from Pesaresi et al. (2011) (0.4 M sucrose, 10 mM NaCl, 5 mM MgCl2, 10 mM tricine KOH pH 7.5, 100 mM ascorbate, 0.2 mM PMSF, 5 mM aminocaproic acid). Samples were centrifuged at 1000g for 5 min at 4°C, then the pellets were resuspended in extraction buffer. The samples were centrifuged again at 1000g for 5 min at 4°C, and the resulting chloroplast enriched pellets suspended in lysis buffer as previously described (Chen and Schröder, 2016) (10 mM Tricine -NaOH pH 7.8) and incubated for 30 min. The samples were centrifuged at 5000g for 5 min at 4°C and the pellets containing the membrane fraction were re-suspended in lysis buffer.

Samples were centrifuged at 1000g for 5 min at 4°C, then the pellets were resuspended in extraction buffer (100 mM Tris pH 6.8, 20% glycerol, 10% SDS). The samples were heated to 40°C for 5 min and then centrifuged at 1300 g for 10 min at room temperature. The supernatant was recovered, and the samples were normalised on the amount of chlorophyll.

### Chlorophyll quantification

Pigments were extracted in 80% acetone then separated and quantified by HPLC-UV as described previously (Campoli et al., 2009).

### RNA sequencing

RNA sequencing was performed with three biological replicates on 100 bulked seedlings per line grown on +N medium for eight days or -N medium for 12 days. RNA was extracted from frozen seedling powder with Nucleozol (Macherey-Nagel) with 4-bromoanisole to reduce DNA and anthocyanin contamination. Total RNA was cleaned and concentrated using RNA Clean & Concentrator^TM^-25 (Zymo Research) according to the manufacturer’s instructions Genomic DNA was removed by treatment with DNase. RNA-seq libraries were constructed by the POPS platform (IPS2) using the TruSeq Stranded mRNA library prep kit (Illumina) with RiboZero plant (Illumina). Libraries were sequenced in single-end (SE) with a read length of 75 bases for each read on a NextSeq500 (Illumina). Approximately 30 million reads by sample were generated. Adapter sequences and bases with a Q-Score below 20 were trimmed out from reads using Trimmomatic (v0.36) (Bolger et al., 2014) and reads shorter than 30 bases after trimming were discarded. Reads corresponding to rRNA sequences were removed using sortMeRNA (v2.1)(Kopylova et al., 2012) against the silva-bac-16s-id90, silva-bac-23s-id98, silva-euk-18s-id95 and silva-euk-28s-id98 databases. Read quality checks were performed using FastQC (Version 0.11.5)(Andrews, 2010). The raw data (fastq) was then aligned against the Arabidopsis transcriptome (Araport11_cdna_20160703_representative_gene_model.fa) concatenated with non-coding RNA (TAIR10.ncrna.fa) using Bowtie2 (version 2.2.9)(Langmead and Salzberg, 2012). Default parameters were used. Reads were counted using a modified version of a command line previously described (Van Verk et al., 2013). Differential expression analysis was performed with SARTools (version 1.7.3)(Varet et al., 2016) using edgeR. Study details and Fastq files were deposited at the European Nucleotide Archive (https://www.ebi.ac.uk/ena) under accession number PRJEB46181. Workflow and analysis reports for RNAseq data analysis are provided in Supporting Information.

### Data analysis

The majority of analysis was conducted in R (R Core Team, 2020) and annotated R markdown scripts are provided in Supporting Information. Graphs were produced using the package ggplot2 (Wickham, 2009) with confidence intervals determined for normally distributed data using the Rmisc package (Hope, 2013), and non-parametric data with boostrap confidence interval calculation using the boot package (Canty and Ripley, 2021; Davison and Hinkley, 1997). Statistical analyses were performed on normally distributed data using ANOVA with posthoc Tukey test and non-parametric data using the Kruskal-Wallis test with posthoc Dunn test in the rstatix package (Kassambara, 2021). For categorical data (Fig. 1B,C) significance was calculated using a proportion test and posthoc Fisher test using the Rstatix and Rcompanion packages (Kassambara, 2021; Salvatore, 2021). All statistical tests included adjustments for multiple comparisons.

GO enrichment analysis for the RNAseq data was performed using a custom prepare_gene_ontology.pl script (https://github.com/cecile-lecampion/gene-ontology-analysis-and-graph) which automatically uses PANTHER and REVIGO for the identification and simplification of enriched GO terms according to the procedure proposed by Bonnot et al. (2019). Results were plotted using ggplot2.

## Supporting information

Table S1 DEGs

Table S2 GO analysis

Table S3 chloroplast DEGs

Table S4 mitochondria DEGs

Table S5 plant materials

Table S6 media composition

R scripts

RNAseq analysis reports

## Acknowledgements

We thank colleagues at the LGBP, Xenie Johnson and Wojciech Nawrocki for critical discussion of the manuscript. We thank Julia Bartoli and Emmanuelle Bouveret for supplying isotope labelled ppGpp. Nucleotide measurements were performed on the IJPB Plant Observatory technological platform, and transcriptomics on the POPS platform which are both supported by Saclay Plant Sciences-SPS (ANR-17-EUR-0007). The work was funded by Agence Nationale de la Recherche (ANR-17-CE13-0005).

## Competing interests

The authors declare no competing interests.

## Supporting information

Table S1 DEGs

Table S2 GO analysis

Table S3 Chloroplast gene expression

Table S4 Mitochondria gene expression

Table S5 List of plant lines used

Table S6 Media composition

Dataset1-RNA-seq analysis reports, R markdown scripts and example analyses.

**Fig. S1.**
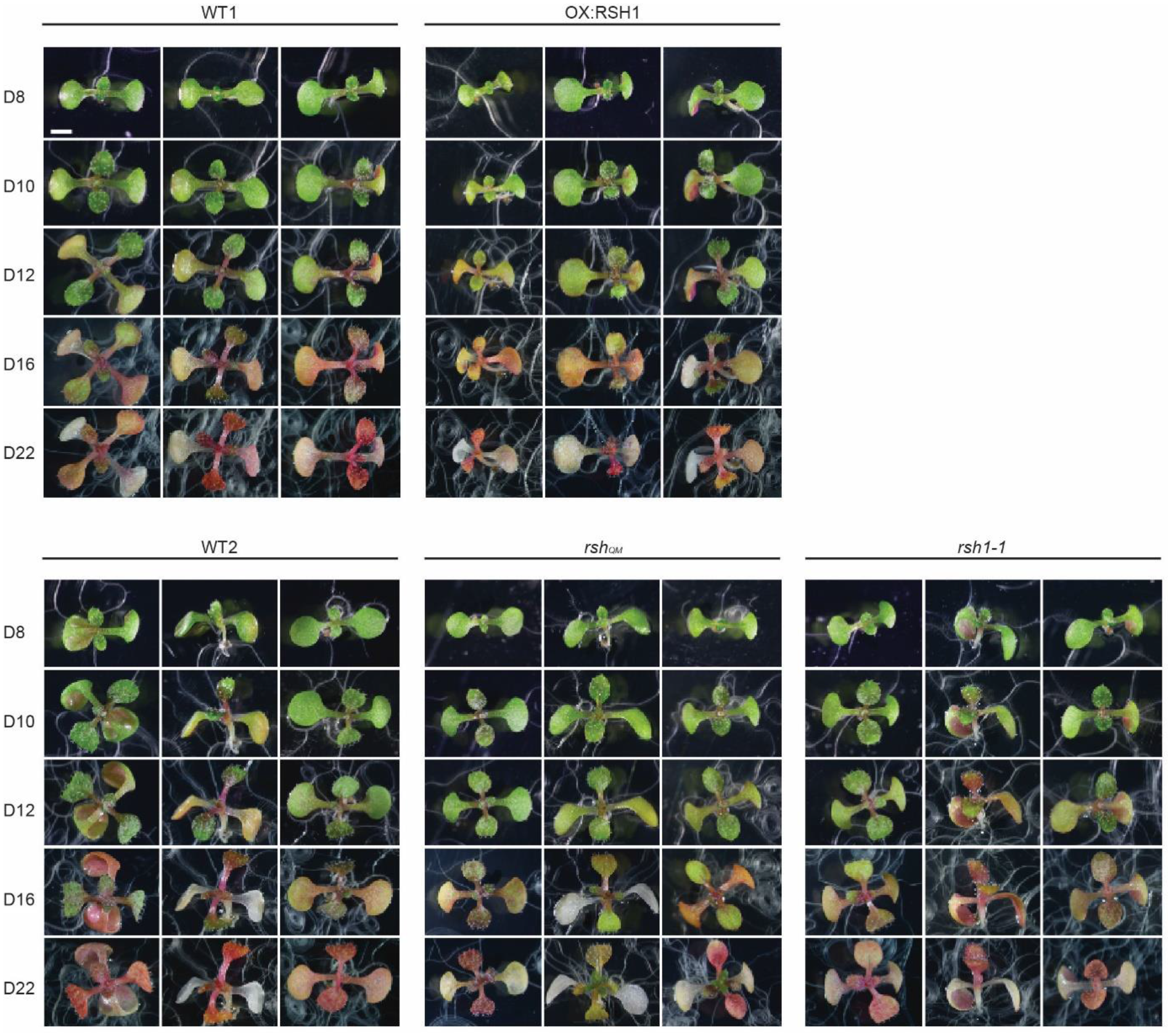
Extended timecourse of nitrogen deprivation. Images of seedlings grown on nitrogen limiting medium for the indicated number of days. Scale, 3 mm.

**Fig. S2.**
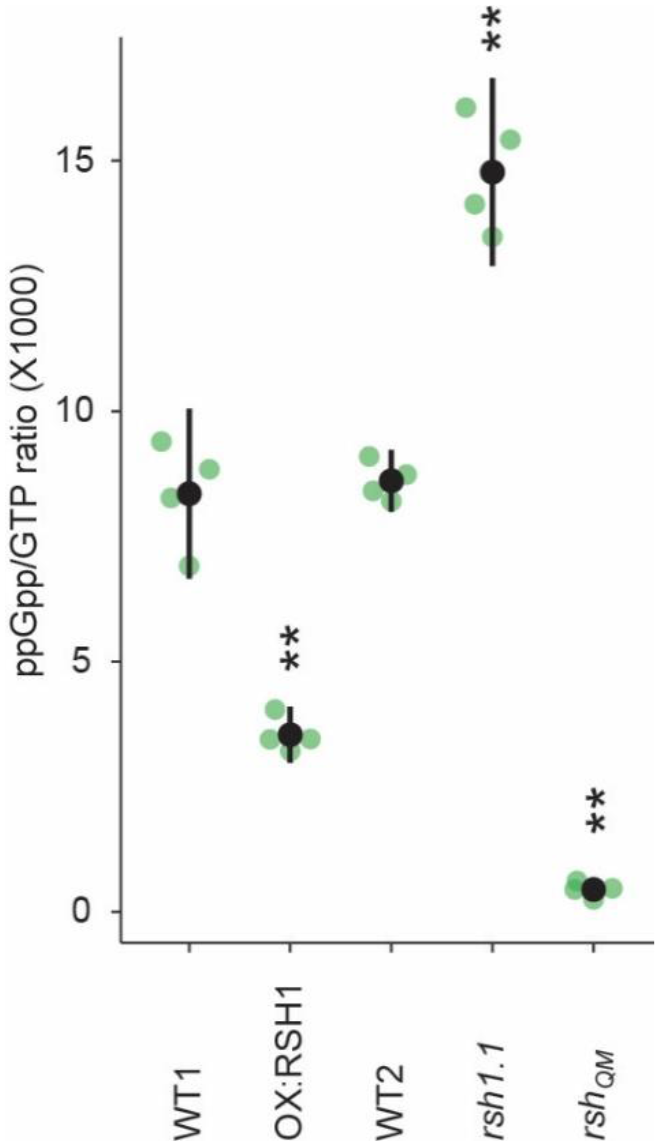
GTP/ppGpp ratios in RSH lines. The ratio of ppGpp to GTP concentration in seedlings grown for ten days on nitrogen limiting media. Mean +/-95% CI, n = 4 biological replicates. Statistical tests shown against respective wild type controls, * *P*<0.05, ** *P*<0.01.

**Fig. S3.**
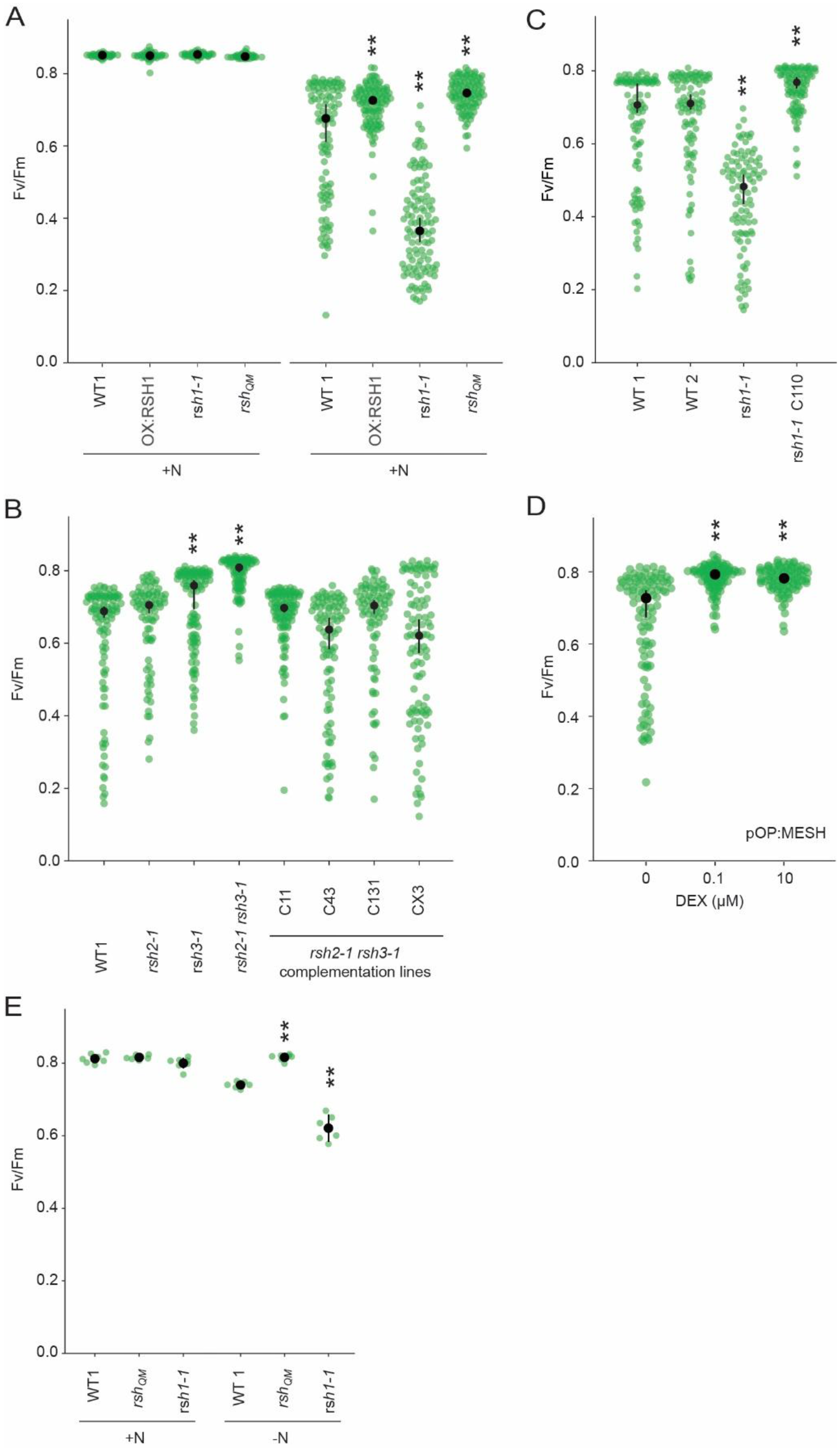
Role of ppGpp in the nitrogen deprivation induced decrease in PSII maximal yield. **(A)** Fv/Fm in seedlings of indicated lines grown 8 days on nitrogen replete media (+N) or 12 days on nitrogen limiting (-N) media. **(B)(C)(D)** Fv/Fm in seedlings of indicated lines grown 12 days on -N. *rsh1-1* C110 is complementation line complemented with the full length *RSH1* (*pRSH1:RSH1*). pOP:MESH is a dexamethasone (DEX) inducible line that expresses a chloroplast targeted ppGpp hydrolase MESH. **(E)** Fv/Fm in mature plants grown on quartz sand supplemented with nitrogen replete media (+N) or nitrogen free (-N) media. Means +/-95% CI, n = 95-100 seedlings (A-D), 6 plants (E). Statistical tests shown against respective wild type controls, * *P*<0.05, ** *P*<0.01.

**Fig. S4.**
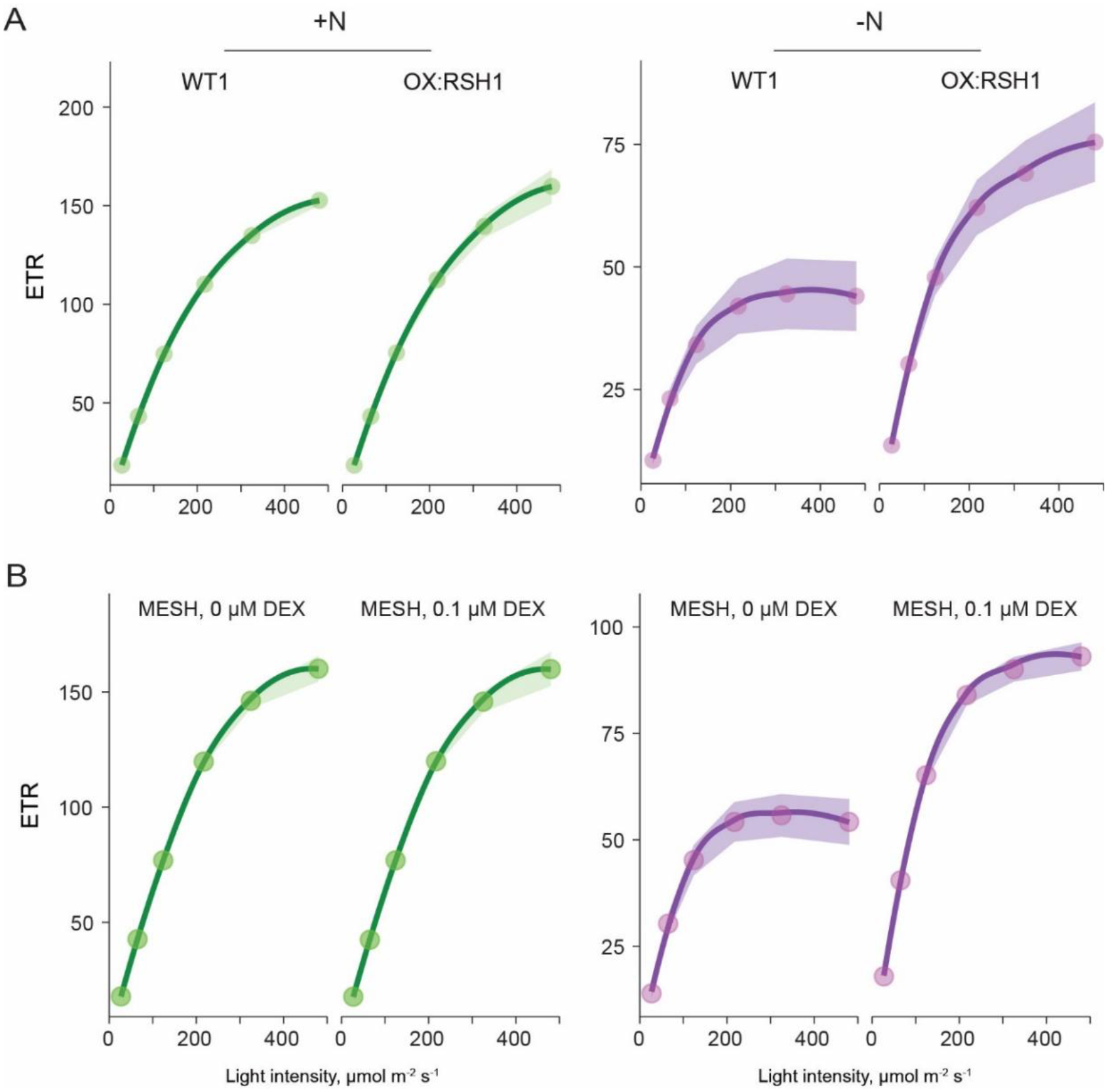
ppGpp is required for reducing ETR during nitrogen deprivation. Relative ETR measurements in different lines grown 8 days on +N media (left) or 12 days on -N media (right). (A) A comparison between OX:RSH1 and the wild type control. (B) ETR in the dexamethasone (DEX) inducible pOP:MESH line grown on non-inducing medium (0 µM DEX), or inducing medium (10 uM DEX). MESH is a chloroplast targeted ppGpp hydrolase. Means +/-95% CI, n = 95-100 seedlings.

**Fig. S5.**
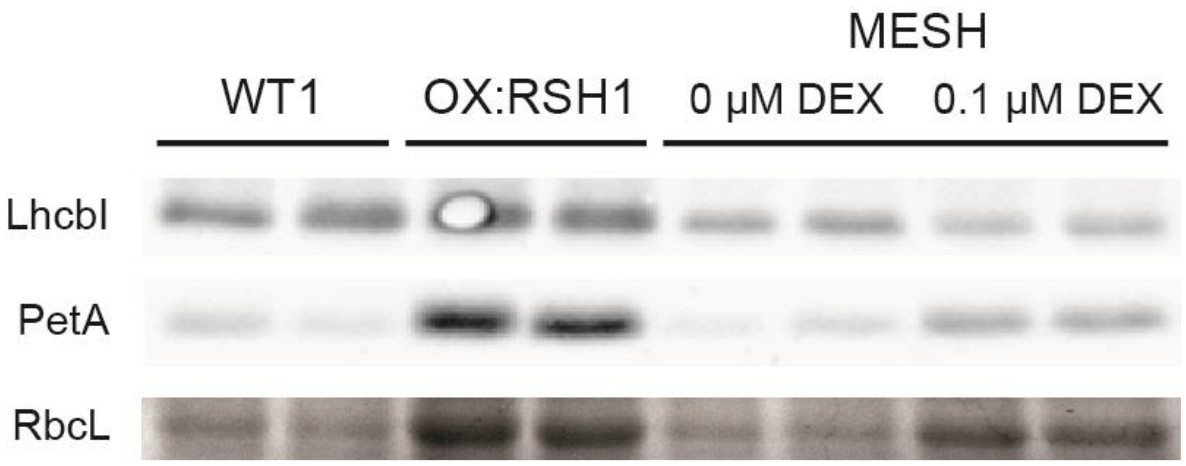
ppGpp depletion by MESH also affects abundance of chloroplast proteins. Immunoblots showing the abundance of the indicated proteins in extracts from seedlings grown in -N for 12 days from two independent biological replicates. MESH seedlings were induced by inclusion of dexamethasone (DEX) or not in the growth medium. RbcL was revealed by Coomassie Brilliant Blue. Equal quantities of total protein were loaded.

**Fig. S6.**
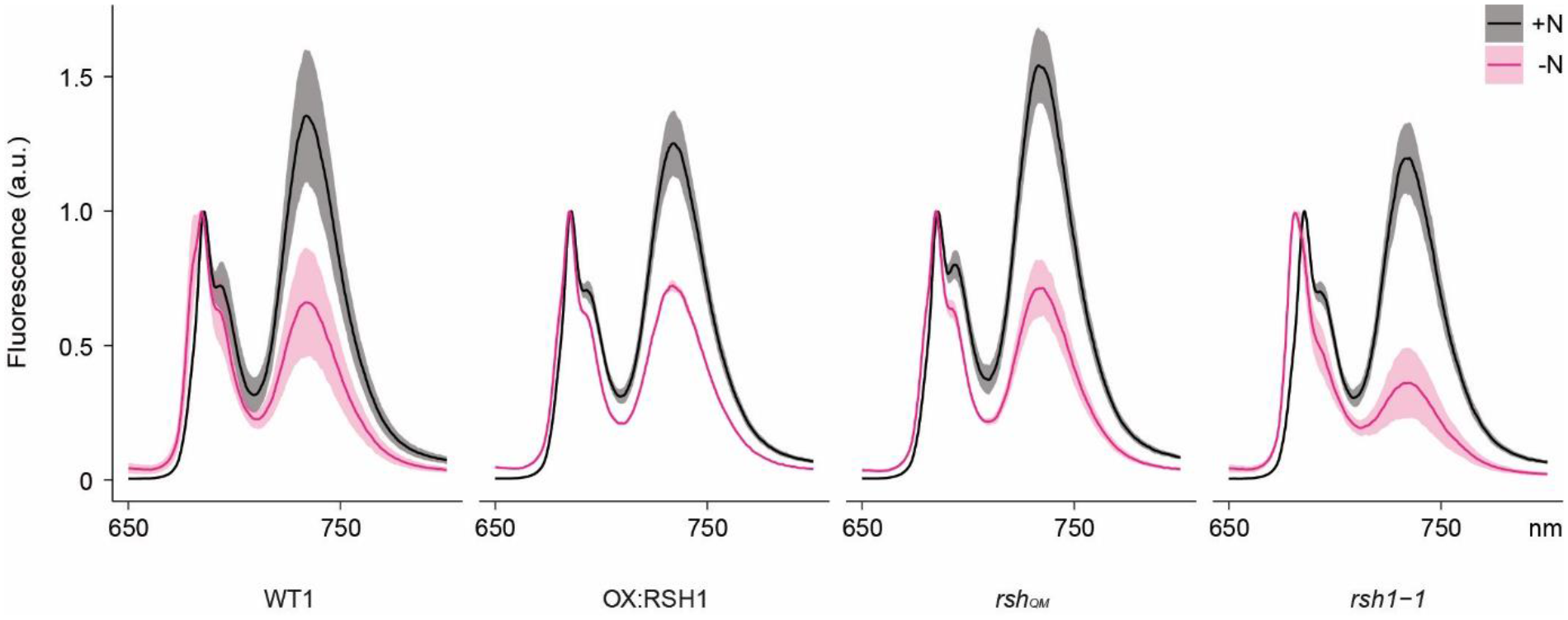
Full 77K chlorophyll fluorescence spectra under nitrogen deprivation. Emission spectrum of chlorophyll fluorescence at 77°K. Measurements made on 8 or 12 day-old seedlings grown under +N or -N. Means ± 95% CI; data from four biological replicates.

**Fig. S7 (1/2).**
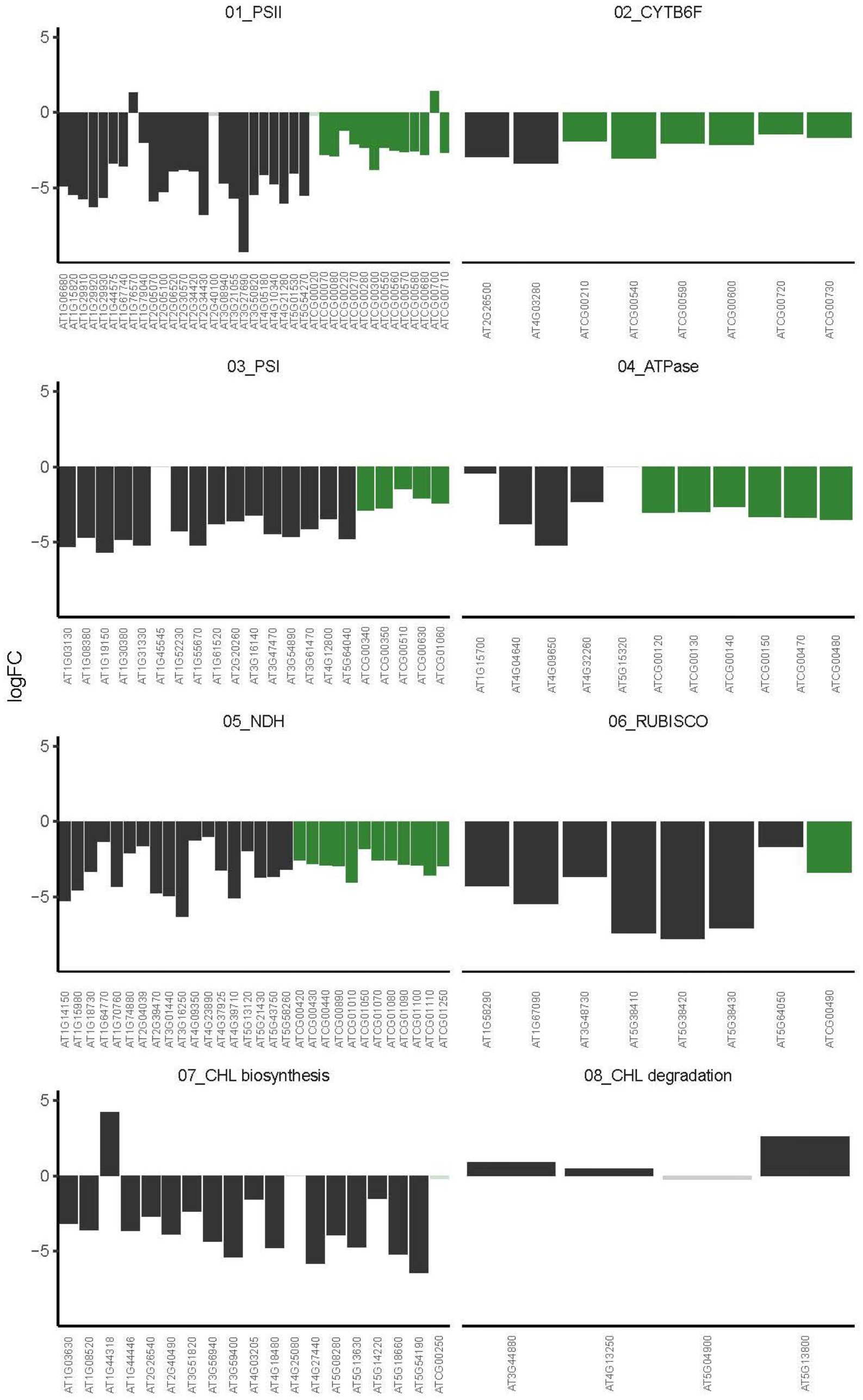
Coordinated regulation of photosynthetic complexes under nitrogen deprivation. Relative transcript levels in wild type -N versus +N for nuclear (black) and chloroplast (green) genes encoding subunits of the indicated photosynthetic complexes. Solid colors indicate significantly different changes in expression, transparent colors indicate non-significant changes.

**Fig. S7 (2/2).**
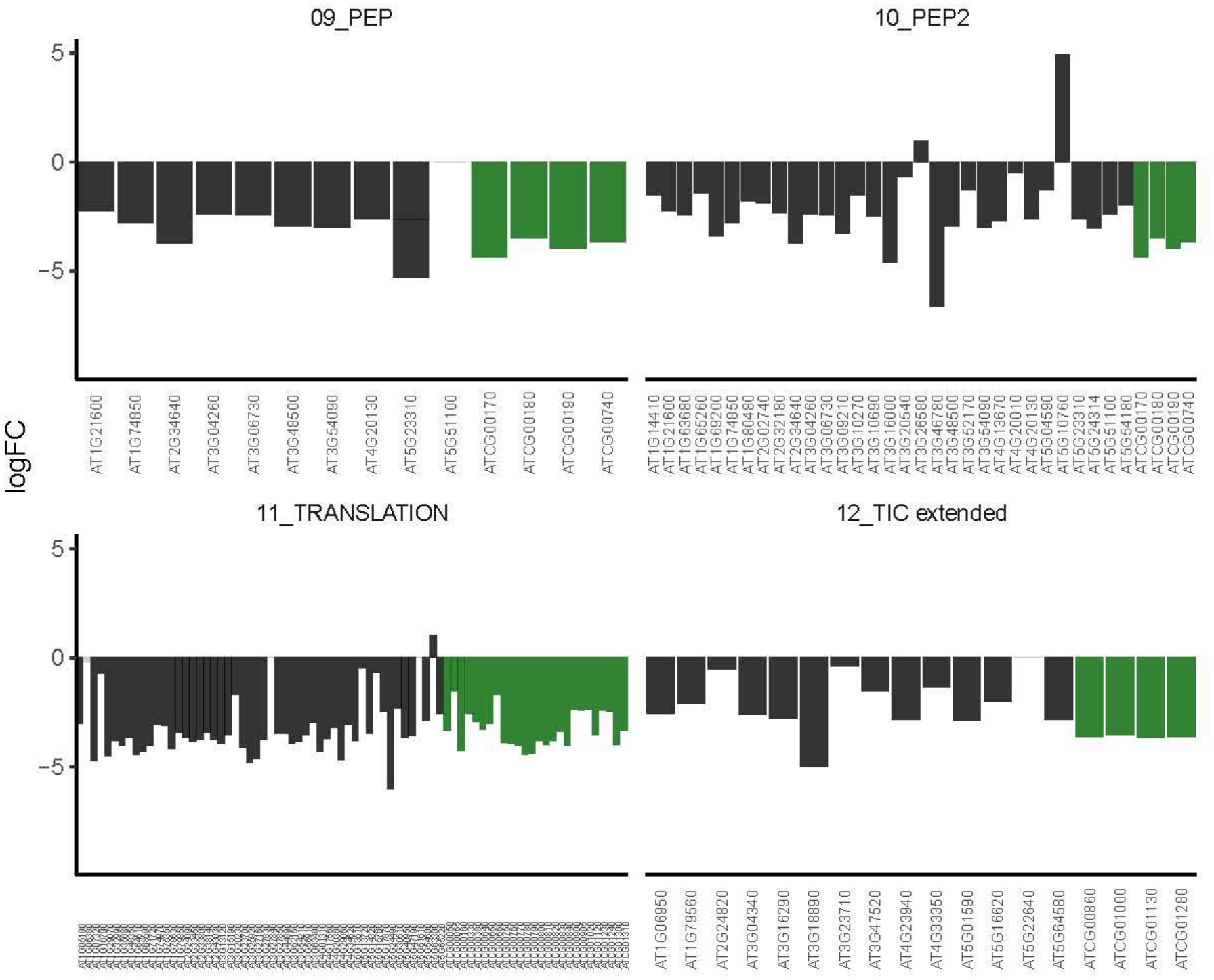
Coordinated regulation of photosynthetic complexes under nitrogen deprivation. Relative transcript levels in wild type -N versus +N for nuclear (black) and chloroplast (green) genes encoding subunits of the indicated photosynthetic complexes. Solid colors indicate significantly different changes in expression, transparent colors indicate non-significant changes.

**Fig. S8 (1/3).**
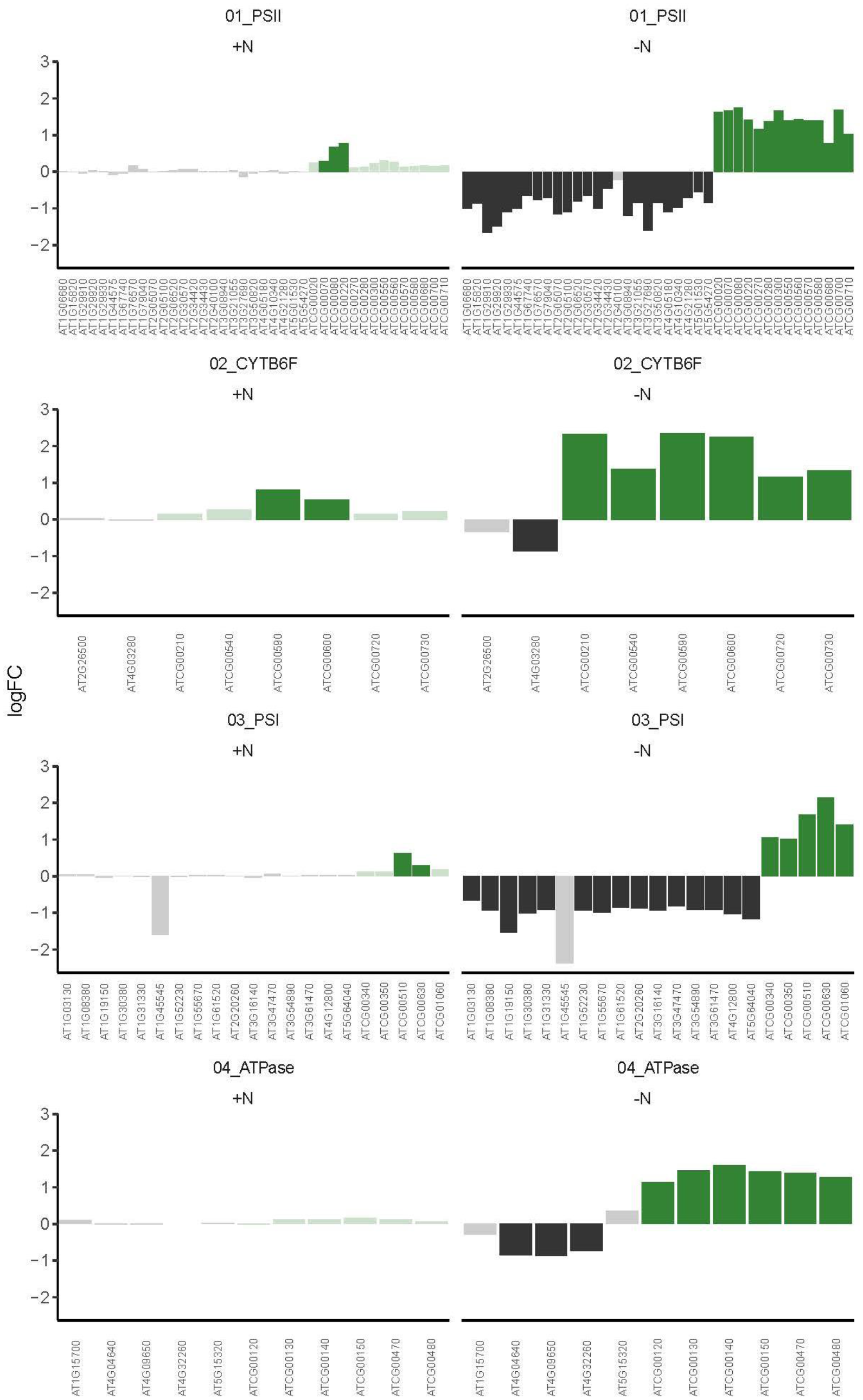
Disrupted regulation of photosynthetic complexes in OX:RSH1. Relative transcript levels under -N for OX:RSH1 versus wild type for nuclear (black) and chloroplast (green) genes encoding subunits of the indicated photosynthetic complexes. Solid colors indicate significantly different changes in expression, transparent colors indicate non-significant changes.

**Fig. S8 (2/3).**
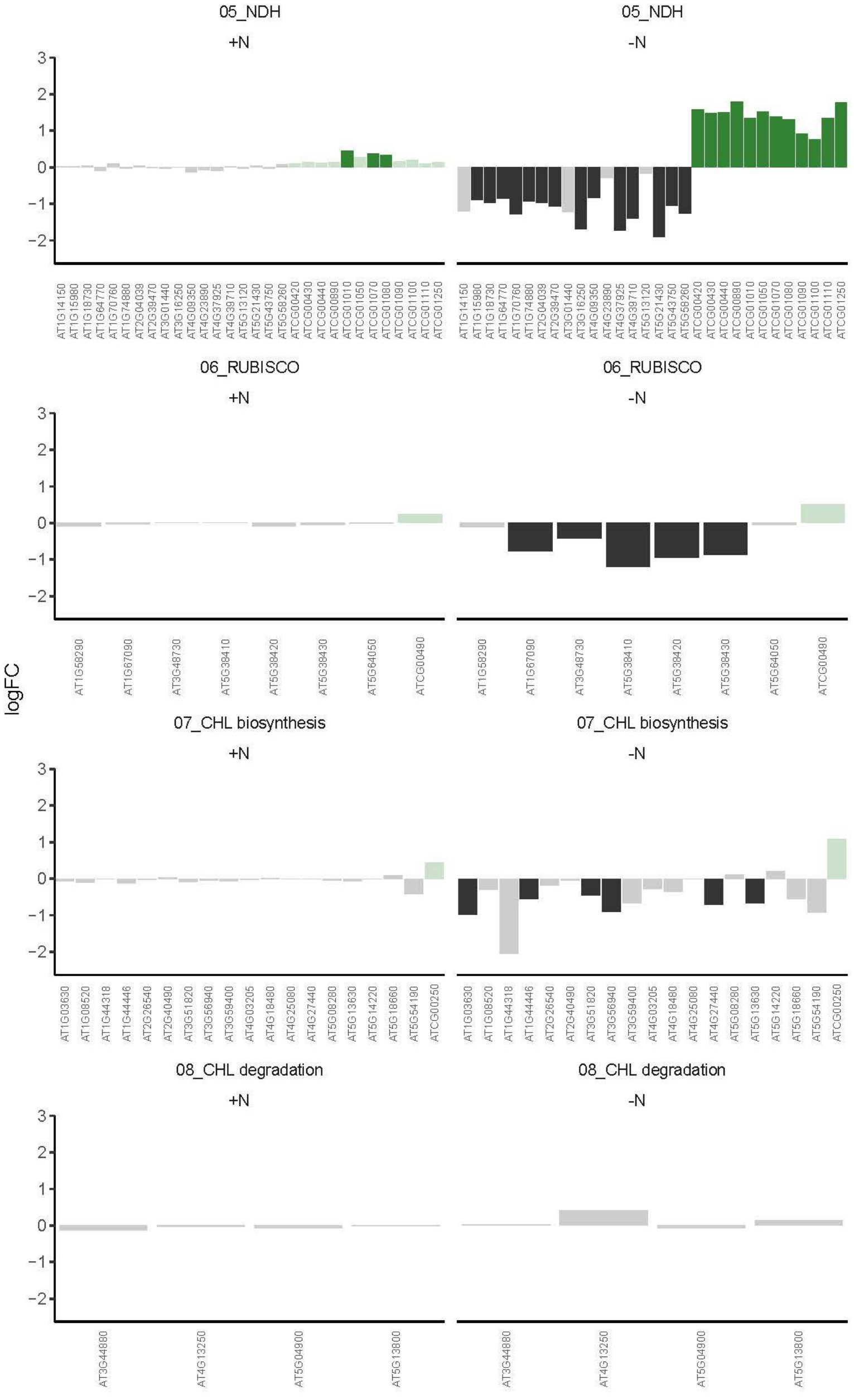
Disrupted regulation of photosynthetic complexes in OX:RSH1. Relative transcript levels under -N for OX:RSH1 versus wild type for nuclear (black) and chloroplast (green) genes encoding subunits of the indicated photosynthetic complexes. Solid colors indicate significantly different changes in expression, transparent colors indicate non-significant changes.

**Fig. S8 (3/3).**
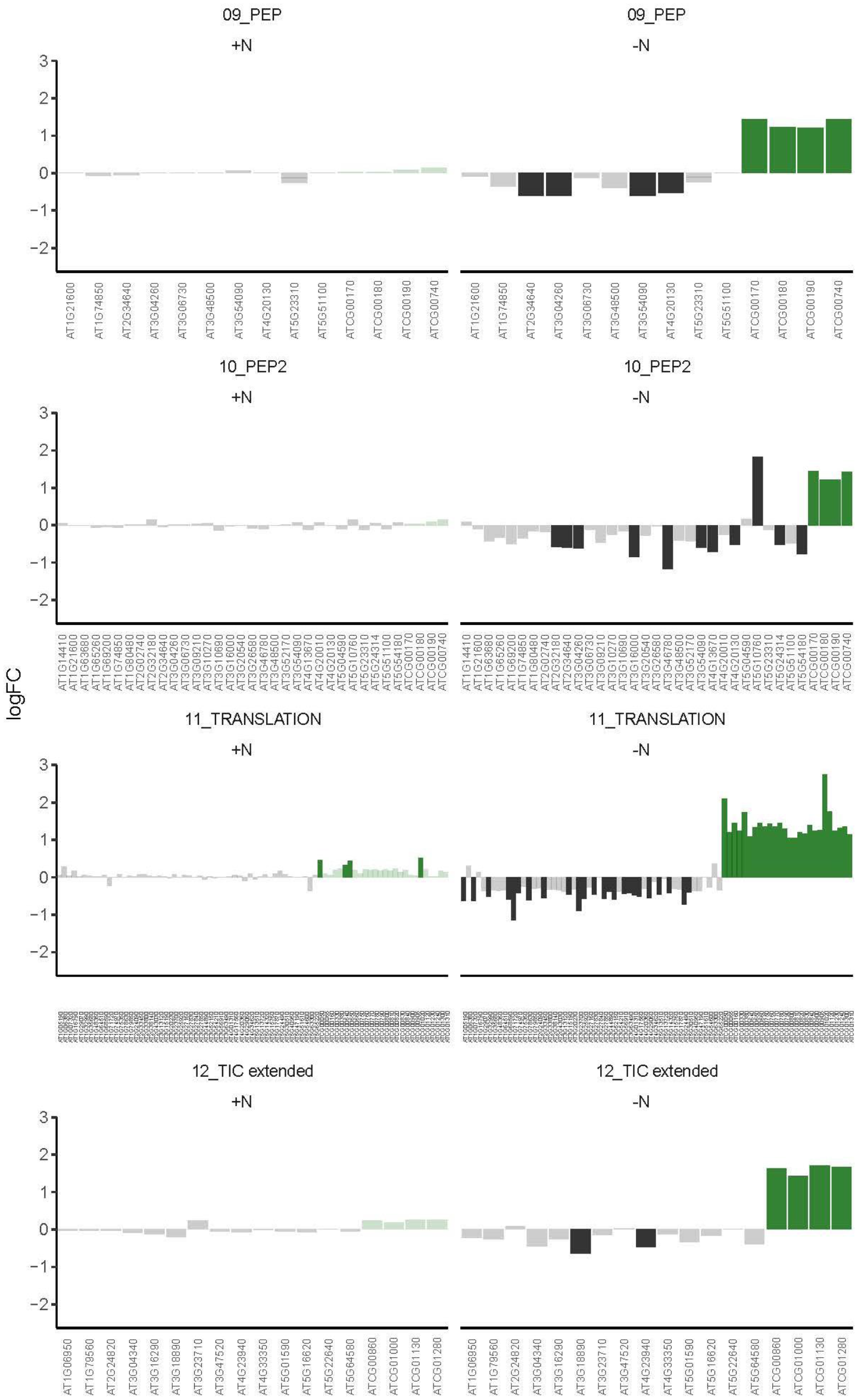
Disrupted regulation of photosynthetic complexes in OX:RSH1. Relative transcript levels under -N for OX:RSH1 versus wild type for nuclear (black) and chloroplast (green) genes encoding subunits of the indicated photosynthetic complexes. Solid colors indicate significantly different changes in expression, transparent colors indicate non-significant changes.

## Notes

### Competing Interest Statement

The authors have declared no competing interest.

## References

Abdelkefi, H., Sugliani, M., Ke, H., Harchouni, S., Soubigou-Taconnat, L., Citerne, S., Mouille, G., Fakhfakh, H., Robaglia, C., and Field, B. (2018). Guanosine tetraphosphate modulates salicylic acid signalling and the resistance of Arabidopsis thaliana to Turnip mosaic virus. Mol. Plant Pathol. 19: 634–646.

Anderson, B.W., Fung, D.K., and Wang, J.D. (2021). Regulatory Themes and Variations by the Stress-Signaling Nucleotide Alarmones (p)ppGpp in Bacteria. Annu. Rev. Genet. 55: doi: 10.1146/annurev-genet-021821-025827.

Andrews, S. (2010). FastQC. A quality control tool for high throughput sequence data. Version 0.11.5 https://www.bioinformatics.babraham.ac.uk/projects/fastqc/.

Atkinson, G.C., Tenson, T., and Hauryliuk, V. (2011). The RelA/SpoT homolog (RSH) superfamily: distribution and functional evolution of ppGpp synthetases and hydrolases across the tree of life. PLoS One 6: e23479.

Avilan, L., Lebrun, R., Puppo, C., Citerne, S., Cuiné, S., Li-Beisson, Y., Menand, B., Field, B., and Gontero, B. (2021). ppGpp influences protein protection, growth and photosynthesis in Phaeodactylum tricornutum. New Phytol. 230: 1517–1532.

Avilan, L., Puppo, C., Villain, A., Bouveret, E., Menand, B., Field, B., and Gontero, B. (2019). RSH enzyme diversity for (p)ppGpp metabolism in Phaeodactylum tricornutum and other diatoms. Sci. Rep. 9: 1–11.

Bange, G., Brodersen, D.E., Liuzzi, A., and Steinchen, W. (2021). Two P or Not Two P: Understanding Regulation by the Bacterial Second Messengers (p)ppGpp. Annu. Rev. Microbiol.

Bartoli, J., Citerne, S., Mouille, G., Bouveret, E., and Field, B. (2020). Quantification of guanosine triphosphate and tetraphosphate in plants and algae using stable isotope-labelled internal standards. Talanta 219: 121261.

Bechtold, U. and Field, B. (2018). Molecular mechanisms controlling plant growth during abiotic stress. J. Exp. Bot. 69: 2753–2758.

Birtic, S., Ksas, B., Genty, B., Mueller, M.J., Triantaphylidès, C., and Havaux, M. (2011). Using spontaneous photon emission to image lipid oxidation patterns in plant tissues. Plant J. 67: 1103–1115.

Bolger, A.M., Lohse, M., and Usadel, B. (2014). Trimmomatic: a flexible trimmer for Illumina sequence data. Bioinforma. Oxf. Engl. 30: 2114–2120.

Boniecka, J., Prusinska, J., Dabrowska, G.B., and Goc, A. (2017). Within and beyond the stringent response-RSH and (p)ppGpp in plants. Planta 246: 817–842.

Bonnot, T., Gillard, M.B., and Nagel, D.H. (2019). A Simple Protocol for Informative Visualization of Enriched Gene Ontology Terms. Bio-Protoc.: e3429–e3429.

Campoli, C., Caffarri, S., Svensson, J.T., Bassi, R., Stanca, A.M., Cattivelli, L., and Crosatti, C. (2009). Parallel pigment and transcriptomic analysis of four barley albina and xantha mutants reveals the complex network of the chloroplast-dependent metabolism. Plant Mol. Biol. 71: 173–191.

Canty, A. and Ripley, B. (2021). boot: Bootstrap R (S-Plus) Functions. Version 1.3–28 https://cran.r-project.org/web/packages/boot/index.html.

Cashel, M. and Gallant, J. (1969). Two compounds implicated in the function of the RC gene of Escherichia coli. Nature 221: 838.

Chen, Y. and Schröder, W. (2016). Comparison of methods for extracting thylakoid membranes of Arabidopsis plants. Physiol. Plant. 156: 3–12.

Crepin, A., Santabarbara, S., and Caffarri, S. (2016). Biochemical and Spectroscopic Characterization of Highly Stable Photosystem II Supercomplexes from Arabidopsis. J. Biol. Chem. 291: 19157–19171.

Davison, A.C. and Hinkley, D.V. (1997). Bootstrap Methods and their Application 1st edition. (Cambridge University Press).

Devireddy, A.R., Zandalinas, S.I., Fichman, Y., and Mittler, R. (2021). Integration of reactive oxygen species and hormone signaling during abiotic stress. Plant J. 105: 459–476.

Domínguez, F. and Cejudo, F.J. (2021). Chloroplast dismantling in leaf senescence. J. Exp. Bot. 72: 5905–5918.

Field, B. (2018). Green magic: regulation of the chloroplast stress response by (p)ppGpp in plants and algae. J. Exp. Bot. 69: 2797–2807.

Galka, P., Santabarbara, S., Khuong, T.T.H., Degand, H., Morsomme, P., Jennings, R.C., Boekema, E.J., and Caffarri, S. (2012). Functional analyses of the plant photosystem I-light-harvesting complex II supercomplex reveal that light-harvesting complex II loosely bound to photosystem II is a very efficient antenna for photosystem I in state II. Plant Cell 24: 2963–2978.

Garai, S. and Tripathy, B.C. (2018). Alleviation of Nitrogen and Sulfur Deficiency and Enhancement of Photosynthesis in Arabidopsis thaliana by Overexpression of Uroporphyrinogen III Methyltransferase (UPM1). Front. Plant Sci. 0.

Harchouni, S., England, S., Vieu, J., Aouane, A., Citerne, S., Legeret, B., Li-Beisson, Y., Menand, B., and Field, B. (2021). Guanosine tetraphosphate (ppGpp) accumulation inhibits chloroplast gene expression and promotes super grana formation in the moss Physcomitrium (Physcomitrella) patens. bioRxiv: 2021.01.06.425534.

Hojka, M., Thiele, W., Tóth, S.Z., Lein, W., Bock, R., and Schöttler, M.A. (2014). Inducible Repression of Nuclear-Encoded Subunits of the Cytochrome b6f Complex in Tobacco Reveals an Extraordinarily Long Lifetime of the Complex. Plant Physiol. 165: 1632– 1646.

Honoki, R., Ono, S., Oikawa, A., Saito, K., and Masuda, S. (2018). Significance of accumulation of the alarmone (p)ppGpp in chloroplasts for controlling photosynthesis and metabolite balance during nitrogen starvation in Arabidopsis. Photosynth Res 135: 299–308.

Hope, R. (2013). Rmisc: Ryan Miscellaneous. Version 1.5 https://cran.r-project.org/package=Rmisc.

Ihara, Y., Ohta, H., and Masuda, S. (2015). A highly sensitive quantification method for the accumulation of alarmone ppGpp in Arabidopsis thaliana using UPLC-ESI-qMS/MS. J Plant Res 128: 511–8.

Imamura, S., Nomura, Y., Takemura, T., Pancha, I., Taki, K., Toguchi, K., Tozawa, Y., and Tanaka, K. (2018). The checkpoint kinase TOR (target of rapamycin) regulates expression of a nuclear-encoded chloroplast RelA-SpoT homolog (RSH) and modulates chloroplast ribosomal RNA synthesis in a unicellular red alga. Plant J. 94: 327–339.

Irving, S.E. and Corrigan, R.M. (2018). Triggering the stringent response: signals responsible for activating (p)ppGpp synthesis in bacteria. Microbiol. Read. Engl. 164: 268–276.

Ito, D., Ihara, Y., Nishihara, H., and Masuda, S. (2017). Phylogenetic analysis of proteins involved in the stringent response in plant cells. J Plant Res 130: 625–634.

Kassambara, A. (2021). Rstatix: Pipe-Friendly Framework for Basic Statistical Tests. Version 0.7.0 https://CRAN.R-project.org/package=rstatix.

Kleine, T. et al. (2021). Acclimation in plants – the Green Hub consortium. Plant J. 106: 23– 40.

Kopylova, E., Noé, L., and Touzet, H. (2012). SortMeRNA: fast and accurate filtering of ribosomal RNAs in metatranscriptomic data. Bioinforma. Oxf. Engl. 28: 3211–3217.

Krieger-Liszkay, A., Krupinska, K., and Shimakawa, G. (2019). The impact of photosynthesis on initiation of leaf senescence. Physiol. Plant. 166: 148–164.

Langmead, B. and Salzberg, S.L. (2012). Fast gapped-read alignment with Bowtie 2. Nat. Methods 9: 357–359.

Lu, C. and Zhang, J. (2000). Photosynthetic CO2 assimilation, chlorophyll fluorescence and photoinhibition as affected by nitrogen deficiency in maize plants. Plant Sci. 151: 135–143.

Luo, J., Havé, M., Clément, G., Tellier, F., Balliau, T., Launay-Avon, A., Guérard, F., Zivy, M., and Masclaux-Daubresse, C. (2020). Integrating multiple omics to identify common and specific molecular changes occurring in Arabidopsis under chronic nitrate and sulfate limitations. J. Exp. Bot. 71: 6471–6490.

Maekawa, M., Honoki, R., Ihara, Y., Sato, R., Oikawa, A., Kanno, Y., Ohta, H., Seo, M., Saito, K., and Masuda, S. (2015). Impact of the plastidial stringent response in plant growth and stress responses. Nat. Plants 1: 15167.

Majeran, W., Wollman, F.-A., and Vallon, O. (2000). Evidence for a Role of ClpP in the Degradation of the Chloroplast Cytochrome b6f Complex. Plant Cell 12: 137–150.

Malnoë, A. (2018). Photoinhibition or photoprotection of photosynthesis? Update on the (newly termed) sustained quenching component qH. Environ. Exp. Bot. 154: 123–133.

Mizusawa, K., Masuda, S., and Ohta, H. (2008). Expression profiling of four RelA/SpoT-like proteins, homologues of bacterial stringent factors, in Arabidopsis thaliana. Planta 228: 553–62.

Montillet, J.-L., Cacas, J.-L., Garnier, L., Montané, M.-H., Douki, T., Bessoule, J.-J., Polkowska-Kowalczyk, L., Maciejewska, U., Agnel, J.-P., Vial, A., and Triantaphylidès, C. (2004). The upstream oxylipin profile of Arabidopsis thaliana: a tool to scan for oxidative stresses. Plant J. 40: 439–451.

Nakagawa, T. et al. (2007). Improved Gateway binary vectors: high-performance vectors for creation of fusion constructs in transgenic analysis of plants. Biosci. Biotechnol. Biochem. 71: 2095–2100.

Nomura, Y., Izumi, A., Fukunaga, Y., Kusumi, K., Iba, K., Watanabe, S., Nakahira, Y., Weber, A.P.M., Nozawa, A., and Tozawa, Y. (2014). Diversity in Guanosine 3′,5′-Bisdiphosphate (ppGpp) Sensitivity among Guanylate Kinases of Bacteria and Plants *. J. Biol. Chem. 289: 15631–15641.

Nunes, M.A., Ramalho, J., and Dias, M.A. (1993). Effect of Nitrogen Supply on the Photosynthetic Performance of Leaves from Coffee Plants Exposed to Bright Light. J. Exp. Bot. 44: 893–899.

Ono, S., Suzuki, S., Ito, D., Tagawa, S., Shiina, T., and Masuda, S. (2020). Plastidial (p)ppGpp Synthesis by the Ca2+-Dependent RelA–SpoT Homolog Regulates the Adaptation of Chloroplast Gene Expression to Darkness in Arabidopsis. Plant Cell Physiol. 61: 2077–2086.

Pausch, P., Steinchen, W., Wieland, M., Klaus, T., Freibert, S.-A., Altegoer, F., Wilson, D.N., and Bange, G. (2018). Structural basis for (p)ppGpp-mediated inhibition of the GTPase RbgA. J. Biol. Chem. 293: 19699–19709.

Pesaresi, P. (2011). Studying translation in Arabidopsis chloroplasts. Methods Mol Biol 774: 209–24.

Pinnola, A. and Bassi, R. (2018). Molecular mechanisms involved in plant photoprotection. Biochem. Soc. Trans. 46: 467–482.

R Core Team (2020). R: A language and environment for statistical computing. R Foundation for Statistical Computing, Vienna, Austria. R 4.1.0 http://www.r-project.org/index.html.

Roberts, D.R., Thompson, J.E., Dumbroff, E.B., Gepstein, S., and Mattoo, A.K. (1987). Differential changes in the synthesis and steady-state levels of thylakoid proteins during bean leaf senescence. Plant Mol. Biol. 9: 343–353.

Rogers, H. and Munné-Bosch, S. (2016). Production and Scavenging of Reactive Oxygen Species and Redox Signaling during Leaf and Flower Senescence: Similar But Different1[OPEN]. Plant Physiol. 171: 1560–1568.

Ronneau, S. and Hallez, R. (2019). Make and break the alarmone: regulation of (p)ppGpp synthetase/hydrolase enzymes in bacteria. FEMS Microbiol. Rev. 43: 389–400.

Safi, A. et al. (2021). GARP transcription factors repress Arabidopsis nitrogen starvation response via ROS-dependent and -independent pathways. J. Exp. Bot. 72: 3881–3901.

Salvatore, M. (2021). rcompanion: functions to support extension education program evaluation in R. Version 2.4.1 https://cran.r-project.org/web/packages/rcompanion/index.html.

Sharma, M., Kretschmer, C., Lampe, C., Stuttmann, J., and Klösgen, R.B. (2019). Targeting specificity of nuclear-encoded organelle proteins with a self-assembling split-fluorescent protein toolkit. J. Cell Sci. 132: jcs230839.

Shumbe, L., D’Alessandro, S., Shao, N., Chevalier, A., Ksas, B., Bock, R., and Havaux, M. (2017). METHYLENE BLUE SENSITIVITY 1 (MBS1) is required for acclimation of Arabidopsis to singlet oxygen and acts downstream of β-cyclocitral. Plant Cell Environ. 40: 216–226.

Stent, G.S. and Brenner, S. (1961). A genetic locus for the regulation of ribonucleic acid synthesis. Proc. Natl. Acad. Sci. U. S. A. 47: 2005–2014.

Sugliani, M., Abdelkefi, H., Ke, H., Bouveret, E., Robaglia, C., Caffarri, S., and Field, B. (2016). An Ancient Bacterial Signaling Pathway Regulates Chloroplast Function to Influence Growth and Development in Arabidopsis. Plant Cell 28: 661–679.

Takahashi, K., Kasai, K., and Ochi, K. (2004). Identification of the bacterial alarmone guanosine 5’-diphosphate 3’-diphosphate (ppGpp) in plants. Proc Natl Acad Sci USA 101: 4320–4324.

Terashima, I. and Evans, J.R. (1988). Effects of Light and Nitrogen Nutrition on the Organization of the Photosynthetic Apparatus in Spinach. Plant Cell Physiol. 29: 143– 155.

Tikkanen, M., Mekala, N.R., and Aro, E.-M. (2014). Photosystem II photoinhibition-repair cycle protects Photosystem I from irreversible damage. Biochim. Biophys. Acta BBA - Bioenerg. 1837: 210–215.

Tyystjarvi, E. and Aro, E.M. (1996). The rate constant of photoinhibition, measured in lincomycin-treated leaves, is directly proportional to light intensity. Proc. Natl. Acad. Sci. 93: 2213–2218.

Van Verk, M.C., Hickman, R., Pieterse, C.M.J., and Van Wees, S.C.M. (2013). RNA-Seq: revelation of the messengers. Trends Plant Sci. 18: 175–179.

Varet, H., Brillet-Guéguen, L., Coppée, J.-Y., and Dillies, M.-A. (2016). SARTools: A DESeq2- and EdgeR-Based R Pipeline for Comprehensive Differential Analysis of RNA-Seq Data. PLoS ONE 11: e0157022.

Varik, V., Oliveira, S.R.A., Hauryliuk, V., and Tenson, T. (2017). HPLC-based quantification of bacterial housekeeping nucleotides and alarmone messengers ppGpp and pppGpp. Sci. Rep. 7: 11022.

Verhoeven, A.S., Demmig-Adams, B., and Adams III, W.W. (1997). Enhanced Employment of the Xanthophyll Cycle and Thermal Energy Dissipation in Spinach Exposed to High Light and N Stress. Plant Physiol. 113: 817–824.

Wei, L., Derrien, B., Gautier, A., Houille-Vernes, L., Boulouis, A., Saint-Marcoux, D., Malnoë, A., Rappaport, F., de Vitry, C., Vallon, O., Choquet, Y., and Wollman, F.-A. (2014). Nitric Oxide–Triggered Remodeling of Chloroplast Bioenergetics and Thylakoid Proteins upon Nitrogen Starvation in Chlamydomonas reinhardtii[W]. Plant Cell 26: 353–372.

Wickham, H. (2009). ggplot2: Elegant Graphics for Data Analysis (Springer-Verlag: New York).

Yamburenko, M.V., Zubo, Y.O., and Borner, T. (2015). Abscisic acid affects transcription of chloroplast genes via protein phosphatase 2C-dependent activation of nuclear genes: repression by guanosine-3’-5’-bisdiphosphate and activation by sigma factor 5. Plant J 82: 1030–1041.

