## Supplementary material for "A ppGpp-mediated brake on photosynthesis is required for acclimation to nitrogen limitation in Arabidopsis": R scripts: script_77K.html

Plot 77K fluorescence data


### Plot 77K fluorescence data

#### ----

#### 1. Introduction

The analysis reported here is part of the manuscript Romand et al. 2021. The script plots 77K chlorophyll fluorescence data where averages and CI have already been calculated.

Working directory: C:/Users/Ben/Desktop/AMUBOX/Shared/ppGpp\_nitrogen\_starvation/Supp data sets/R scripts/R markdown 77K plot Fig\_3

##### Setup R packages

```
if (!require(knitr)) { install.packages("knitr", repos = "http://cran.us.r-project.org") }
if (!require(ggplot2)) { install.packages("ggplot2", repos = "http://cran.us.r-project.org") }

library(knitr)
library(ggplot2)
library(readxl)
```

#### 2. Data

Data is imported from an excel file containing a column for the wavelength `x` in nm, a column of fluorescence values `y` values, a column of 95% CI values `ci`, a column for the different lines `line` and a column for the different treatment conditions `cond`

```
 data1 <- read_excel("77K.xlsx")
```

#### 3. Plot Graph

Graphs are now plotted.

```
 # With 95% ci, all singing all dancing ZOOMED 660-710
p<-ggplot(data=data1, aes(x=x, y=y, fill=cond))+
  geom_ribbon(aes(ymin=y-ci, ymax=y+ci,x=x, fill=cond),alpha=0.3)+
  geom_line(data=data1,aes(x=x, y=y, color=cond), size=0.3)+
  scale_fill_manual(values=c("black", "red"))+
  scale_color_manual(values=c("black", "red"))+
  scale_x_continuous(breaks=seq(660,710,20), limits=c(660,710))+
   scale_y_continuous(breaks=seq(0,1,0.5), limits=c(-0.01,1.01))+
  theme_classic()+
  facet_grid(,vars(line))

print(p)
```

```
ggsave("zoom77K.pdf",width = 8, height = 3)

# With 95% ci, all singing all dancing  FULL 650-800
p<-ggplot(data=data1, aes(x=x, y=y, fill=cond))+
  geom_ribbon(aes(ymin=y-ci, ymax=y+ci,x=x, fill=cond),alpha=0.3)+
  geom_line(data=data1,aes(x=x, y=y, color=cond), size=0.5)+
  scale_fill_manual(values=c("black", "red"))+
  scale_color_manual(values=c("black", "red"))+
  scale_x_continuous(breaks=seq(650,800,100), limits=c(650,800))+
  theme_classic()+
  facet_grid(,vars(line))

print(p)
```

```
ggsave("full77K.pdf",width = 8, height = 3)
```

#### 4. R session information

```
InfoSession <- devtools::session_info()
print(InfoSession)
```

```
## - Session info ---------------------------------------------------------------
##  setting  value                       
##  version  R version 4.1.0 (2021-05-18)
##  os       Windows 10 x64              
##  system   x86_64, mingw32             
##  ui       RTerm                       
##  language (EN)                        
##  collate  English_United Kingdom.1252 
##  ctype    English_United Kingdom.1252 
##  tz       Europe/Paris                
##  date     2021-07-15                  
## 
## - Packages -------------------------------------------------------------------
##  package     * version date       lib source        
##  assertthat    0.2.1   2019-03-21 [1] CRAN (R 4.1.0)
##  cachem        1.0.5   2021-05-15 [1] CRAN (R 4.1.0)
##  callr         3.7.0   2021-04-20 [1] CRAN (R 4.1.0)
##  cellranger    1.1.0   2016-07-27 [1] CRAN (R 4.1.0)
##  cli           2.5.0   2021-04-26 [1] CRAN (R 4.1.0)
##  colorspace    2.0-1   2021-05-04 [1] CRAN (R 4.1.0)
##  crayon        1.4.1   2021-02-08 [1] CRAN (R 4.1.0)
##  DBI           1.1.1   2021-01-15 [1] CRAN (R 4.1.0)
##  desc          1.3.0   2021-03-05 [1] CRAN (R 4.1.0)
##  devtools      2.4.2   2021-06-07 [1] CRAN (R 4.1.0)
##  digest        0.6.27  2020-10-24 [1] CRAN (R 4.1.0)
##  dplyr         1.0.7   2021-06-18 [1] CRAN (R 4.1.0)
##  ellipsis      0.3.2   2021-04-29 [1] CRAN (R 4.1.0)
##  evaluate      0.14    2019-05-28 [1] CRAN (R 4.1.0)
##  fansi         0.5.0   2021-05-25 [1] CRAN (R 4.1.0)
##  farver        2.1.0   2021-02-28 [1] CRAN (R 4.1.0)
##  fastmap       1.1.0   2021-01-25 [1] CRAN (R 4.1.0)
##  fs            1.5.0   2020-07-31 [1] CRAN (R 4.1.0)
##  generics      0.1.0   2020-10-31 [1] CRAN (R 4.1.0)
##  ggplot2     * 3.3.4   2021-06-16 [1] CRAN (R 4.1.0)
##  glue          1.4.2   2020-08-27 [1] CRAN (R 4.1.0)
##  gtable        0.3.0   2019-03-25 [1] CRAN (R 4.1.0)
##  highr         0.9     2021-04-16 [1] CRAN (R 4.1.0)
##  htmltools     0.5.1.1 2021-01-22 [1] CRAN (R 4.1.0)
##  knitr       * 1.33    2021-04-24 [1] CRAN (R 4.1.0)
##  labeling      0.4.2   2020-10-20 [1] CRAN (R 4.1.0)
##  lifecycle     1.0.0   2021-02-15 [1] CRAN (R 4.1.0)
##  magrittr      2.0.1   2020-11-17 [1] CRAN (R 4.1.0)
##  memoise       2.0.0   2021-01-26 [1] CRAN (R 4.1.0)
##  munsell       0.5.0   2018-06-12 [1] CRAN (R 4.1.0)
##  pillar        1.6.1   2021-05-16 [1] CRAN (R 4.1.0)
##  pkgbuild      1.2.0   2020-12-15 [1] CRAN (R 4.1.0)
##  pkgconfig     2.0.3   2019-09-22 [1] CRAN (R 4.1.0)
##  pkgload       1.2.1   2021-04-06 [1] CRAN (R 4.1.0)
##  prettyunits   1.1.1   2020-01-24 [1] CRAN (R 4.1.0)
##  processx      3.5.2   2021-04-30 [1] CRAN (R 4.1.0)
##  ps            1.6.0   2021-02-28 [1] CRAN (R 4.1.0)
##  purrr         0.3.4   2020-04-17 [1] CRAN (R 4.1.0)
##  R6            2.5.0   2020-10-28 [1] CRAN (R 4.1.0)
##  Rcpp          1.0.6   2021-01-15 [1] CRAN (R 4.1.0)
##  readxl      * 1.3.1   2019-03-13 [1] CRAN (R 4.1.0)
##  remotes       2.4.0   2021-06-02 [1] CRAN (R 4.1.0)
##  rlang         0.4.11  2021-04-30 [1] CRAN (R 4.1.0)
##  rmarkdown     2.9     2021-06-15 [1] CRAN (R 4.1.0)
##  rprojroot     2.0.2   2020-11-15 [1] CRAN (R 4.1.0)
##  scales        1.1.1   2020-05-11 [1] CRAN (R 4.1.0)
##  sessioninfo   1.1.1   2018-11-05 [1] CRAN (R 4.1.0)
##  stringi       1.6.1   2021-05-10 [1] CRAN (R 4.1.0)
##  stringr       1.4.0   2019-02-10 [1] CRAN (R 4.1.0)
##  testthat      3.0.3   2021-06-16 [1] CRAN (R 4.1.0)
##  tibble        3.1.2   2021-05-16 [1] CRAN (R 4.1.0)
##  tidyselect    1.1.1   2021-04-30 [1] CRAN (R 4.1.0)
##  usethis       2.0.1   2021-02-10 [1] CRAN (R 4.1.0)
##  utf8          1.2.1   2021-03-12 [1] CRAN (R 4.1.0)
##  vctrs         0.3.8   2021-04-29 [1] CRAN (R 4.1.0)
##  withr         2.4.2   2021-04-18 [1] CRAN (R 4.1.0)
##  xfun          0.24    2021-06-15 [1] CRAN (R 4.1.0)
##  yaml          2.2.1   2020-02-01 [1] CRAN (R 4.1.0)
## 
## [1] C:/Users/Ben/Documents/R/win-library/4.1
## [2] C:/Program Files/R/R-4.1.0/library
```

#### 5. Citations

1. R Core Team. 2020. R: A Language and Environment for Statistical Computing. Vienna, Austria: R Foundation for Statistical Computing. https://www.R-project.org/.
2. Wickham, Hadley. 2016. Ggplot2: Elegant Graphics for Data Analysis. Springer-Verlag New York. https://ggplot2.tidyverse.org.
3. Wickham, Hadley, and Jennifer Bryan. 2019. Readxl: Read Excel Files. https://CRAN.R-project.org/package=readxl.
