## Supplementary material for "A ppGpp-mediated brake on photosynthesis is required for acclimation to nitrogen limitation in Arabidopsis": R scripts: script_77K.Rmd

````
---
title: "Plot 77K fluorescence data"
author: ""
date: "`r format(Sys.time(), '%H:%M, %d %B, %Y')`"
output:
   html_document:
      toc: true
      toc_float: true
      toc_depth: 4
      theme: cerulean
      highlight: kate
---

```{r setup, include=FALSE}
knitr::opts_chunk$set(echo = TRUE)
```

## 1. Introduction

The analysis reported here is part of the manuscript Romand et al. 2021. The script plots 77K chlorophyll fluorescence data where averages and CI have already been calculated.


<br/>

`r {"Working directory:"}`
`r  getwd()`

<br/>

### Setup R packages

```{r warning=FALSE, message= FALSE}

if (!require(knitr)) { install.packages("knitr", repos = "http://cran.us.r-project.org") }
if (!require(ggplot2)) { install.packages("ggplot2", repos = "http://cran.us.r-project.org") }

library(knitr)
library(ggplot2)
library(readxl)

```

## 2. Data

Data is imported from an excel file containing a column for the wavelength  `x` in nm, a column of fluorescence values `y` values,  a column of 95% CI values `ci`,  a column for the different lines `line` and a column for the different treatment conditions `cond`

```{r}

 data1 <- read_excel("77K.xlsx")


```

## 3. Plot Graph

Graphs are now plotted.

```{r warning=FALSE, message=FALSE}
 # With 95% ci, all singing all dancing ZOOMED 660-710
p<-ggplot(data=data1, aes(x=x, y=y, fill=cond))+
  geom_ribbon(aes(ymin=y-ci, ymax=y+ci,x=x, fill=cond),alpha=0.3)+
  geom_line(data=data1,aes(x=x, y=y, color=cond), size=0.3)+
  scale_fill_manual(values=c("black", "red"))+
  scale_color_manual(values=c("black", "red"))+
  scale_x_continuous(breaks=seq(660,710,20), limits=c(660,710))+
   scale_y_continuous(breaks=seq(0,1,0.5), limits=c(-0.01,1.01))+
  theme_classic()+
  facet_grid(,vars(line))

print(p)

ggsave("zoom77K.pdf",width = 8, height = 3)

# With 95% ci, all singing all dancing  FULL 650-800
p<-ggplot(data=data1, aes(x=x, y=y, fill=cond))+
  geom_ribbon(aes(ymin=y-ci, ymax=y+ci,x=x, fill=cond),alpha=0.3)+
  geom_line(data=data1,aes(x=x, y=y, color=cond), size=0.5)+
  scale_fill_manual(values=c("black", "red"))+
  scale_color_manual(values=c("black", "red"))+
  scale_x_continuous(breaks=seq(650,800,100), limits=c(650,800))+
  theme_classic()+
  facet_grid(,vars(line))

print(p)

ggsave("full77K.pdf",width = 8, height = 3)
 
```


## 4. R session information

```{r}
InfoSession <- devtools::session_info()
print(InfoSession)

```

## 5. Citations


1. R Core Team. 2020. R: A Language and Environment for Statistical Computing. Vienna, Austria: R Foundation for Statistical Computing. https://www.R-project.org/.

2. Wickham, Hadley. 2016. Ggplot2: Elegant Graphics for Data Analysis. Springer-Verlag New York. https://ggplot2.tidyverse.org.

3. Wickham, Hadley, and Jennifer Bryan. 2019. Readxl: Read Excel Files. https://CRAN.R-project.org/package=readxl.
````
