## Supplementary material for "A ppGpp-mediated brake on photosynthesis is required for acclimation to nitrogen limitation in Arabidopsis": R scripts: script_ci_G4P.html

Plot and statistical analysis


### Plot and statistical analysis

#### ----

#### 1. Introduction

The analysis reported here is part of the manuscript Romand et al. 2021. The script performs statistical analysis on data grouped by `Treatment` and `Plant Line` and plots a graph. Different approaches are used depending on the normality of the data as determined by the Shapiro test. If data follow a normal distribution, then the script performs Anova and post-hoc Tukey tests and draws a plot with confidence interval centered on mean. If data do not follow a normal distribution then a Kruskal-Wallis and a pot-hoc Dunn test are performed. The same plot is drawn using the medians and boostrap calculated 95% confidence intervals. The plots are also saved as pdf and vector (SVG) files.

#### 2. Data

A 3 column data set in `xlsx`. Column 1, `Line`; column 2, `Treatment` and column 3 the measured quantity, in this case `G4P`. The 2 first columns are factors that are used to group the data.

This script can be simply modified to analyse different parameters or different treatments. It can also be modified to analyse data grouped by additional factors such as Time.

#### 3. Data analysis

##### 3.1. Code

###### Set variables for personalisation

```
#1. Data
#------------------------------------------------------------------------------

# Put your input file in variable DATA 
DATA <- "G4P_mutants.xlsx"


#2. Parameters for plot
#------------------------------------------------------------------------------
# Points color
COLOUR <- "#00B050"

# Select a palette for CI color
PALETTE <- "PuRd"
```

Working directory: C:/Users/Ben/Desktop/AMUBOX/Shared/ppGpp\_nitrogen\_starvation/model markdown/markdown CI G4P

###### Setup R packages

```
if (!require(ggbeeswarm)) { install.packages("ggbeeswarm", repos = "http://cran.us.r-project.org") }
if (!require(Rmisc)) { install.packages("Rmisc", repos = "http://cran.us.r-project.org") }
if (!require(ggplot2)) { install.packages("ggplot2", repos = "http://cran.us.r-project.org") }
if (!require(RColorBrewer)) { install.packages("RColorBrewer", repos = "http://cran.us.r-project.org") }
if (!require(dplyr)) { install.packages("dplyr", repos = "http://cran.us.r-project.org") }
if (!require(rstatix)) { install.packages("rstatix", repos = "https://cloud.r-project.org") }
if (!require(purrr)) { install.packages("purrr", repos = "http://cran.us.r-project.org") }
if (!require(tidyr)) { install.packages("tidyr", repos = "http://cran.us.r-project.org") }
if (!require(magrittr)) { install.packages("magrittr", repos = "http://cran.us.r-project.org") }
if (!require(boot)) { install.packages("boot", repos = "http://cran.us.r-project.org") }
if (!require(devtools)) { install.packages("devtools", repos = "http://cran.us.r-project.org") }
if (!require(knitr)) { install.packages("knitr", repos = "http://cran.us.r-project.org") }
if (!require(svglite)) { install.packages("svglite", repos = "http://cran.us.r-project.org") }


library(ggplot2)
library(ggbeeswarm)
library(Rmisc)
library(readxl)
library(RColorBrewer)
library(rstatix)
library(boot)
library(magrittr)
library(dplyr)
library(purrr)
library(tidyr)
library(rstatix)
library(knitr)
library(svglite)
```

###### Functions used in the script

```
#===============================================================================
# Draw the plot if data follow a normal distribution
#===============================================================================

plot_normal <- function(df, my_colours, my_summary) {
    p<-ggplot(data = df, aes(x=Line, y=G4P)) +
        geom_quasirandom(dodge.width=0.8,alpha = 0.6, colour = COLOUR)+
        geom_pointrange(data = my_summary, aes(ymin=G4P-ci, ymax=G4P+ci, color=Line), position=position_dodge(width=0.8))+
        scale_colour_manual(values=my_colours)+
        facet_wrap(~Treatment, strip.position = "bottom", scales = "free_x") +
        theme_classic() + 
        theme(panel.spacing = unit(0, "lines"), 
              strip.background = element_blank(),
              strip.placement = "outside")+
        theme(axis.text.x = element_text(angle = 90, vjust = 0.5, hjust=1))
    print(p)
}

#===============================================================================
# Draw the plot if data do not follow a normal distribution
#===============================================================================

plot_not_normal <- function(df, my_colours, conf_int) {
    p<-ggplot(data=df, aes(x=Line, y=G4P)) +
        geom_quasirandom(dodge.width=0.8,alpha = 0.6, colour = COLOUR)+
        geom_linerange(data = booted_summary, aes(ymin=lower_ci_perc, ymax=upper_ci_perc, color=Line), position = position_dodge(width=0.8))+
        geom_point(data=booted_summary, aes(y=G4P, color=Line), size = 2, 
                   position=position_dodge(width=0.8))+
        scale_colour_manual(values=my_colours)+
        facet_wrap(~Treatment, strip.position = "bottom", scales = "free_x") +
        theme_classic() + 
        theme(panel.spacing = unit(0, "lines"), 
              strip.background = element_blank(),
              strip.placement = "outside")+
        theme(axis.text.x = element_text(angle = 90, vjust = 0.5, hjust=1))
        print(p)
       }

#===============================================================================
# Check normality of the data. Exit the loop if one set of data doesn't follow a 
# normal distribution.
# Return TRUE if all data follow a normal distribution
#===============================================================================

check_normality <- function(shapiro_df) {
    # Data are supposed to follow normal distribution
    flag_normal <- TRUE
    
    for (i in 1 : nrow(shapiro_df)) {
        if(shapiro_df[i, 4] > 0.05) {
            
        } else {
            
            flag_normal <- FALSE
            break
        }
    }
    return(flag_normal)
}

#===============================================================================
#Function to calculate the median (for bootstrapping)
#===============================================================================

samplemedian <- function(d, i) {
    median(d[i])
}

#===============================================================================
#Function for Dunn test
#===============================================================================
test_dunn <- function() {
    pval <- as.data.frame(df %>% group_by(Treatment) %>% dunn_test(G4P ~ Line, p.adjust.method = "BY"))
    print(df %>% group_by(Treatment) %>% dunn_test(G4P ~ Line, p.adjust.method = "BY"))
    return(pval)
}
```

###### Main script

```
# Import Data
df <- read_excel(DATA, col_types = c("text", "text", "numeric"))
df$Line <- make.names(df$Line)

# Define color
my_colours = brewer.pal(n = 9, PALETTE)[9:3]

# Determining data normality status
shapiro_df <- df %>%group_by(Treatment, Line)%>%
    summarise(statistic = shapiro.test(G4P)$statistic, 
              p.value = shapiro.test(G4P)$p.value)

flag_normal <- check_normality(shapiro_df)


# Data treatement according to normality status
if(flag_normal == TRUE) {
    #print("Data follow a normal distribution")
    
    # Summary
    my_summary <- summarySE(df, measurevar="G4P", groupvars=c("Line", "Treatment"))
    
    # Plot
    plot_normal(df, my_colours, my_summary)
    
    # Stats
    anova_results <- df %>% group_by(Treatment) %>%  anova_test(G4P ~ Line)
    
    tukey_results <- df %>% group_by(Treatment) %>%  tukey_hsd(G4P ~ Line)
    
    
    # Save plot
    ggsave("my_ggplot1.pdf", width=4, height=5)
    ggsave("my_ggplot1.svg", width=4, height=5)
    
} else {
    #print("Data do not follow a normal distribution")
    
    # Calculate Median and 95%CI by bootstrap (Largely based on code by Peter Kamerman: 
    #https://www.painblogr.org/2017-10-18-purrring-through-bootstraps.html)
    
    #1. Set the seed
    set.seed(12345)
    
    #2. Create nested datafile to allow boostrapping on each subgroup.
    my_data_nested <- df %>% group_by(Line,Treatment) %>% nest()
    
    #3. Add boostrapped median values to my_nested_data
    my_data_nested %<>% dplyr::mutate(booted = purrr::map(.x = data, ~ boot::boot(data = .x$G4P,statistic = samplemedian, R = 10000, stype = "i")))
    
    #4. Plot each <S3: boot> object to check boostrapping worked!
    ## Note: Saved to an object (plots) to stop the summary being printed
    plots<-purrr::map(.x = my_data_nested$booted, ~ plot(.x))
    print(plots)
    
    #5. Add boostrapped 95%CI around median values to my_nested_data, 
    #95%CI determined based on "perc" & "bca"
    my_data_nested %<>% mutate(booted_ci = purrr::map(.x = booted, 
                                                      ~ boot::boot.ci(.x,conf = 0.95, 
                                                                      type = c("perc", "bca"))))
    
    #6. Table of each <S3: bootci> object to check
    ci_table <-purrr::map(.x = my_data_nested$booted_ci, ~ print(.x))
    
    #7. Get the stats (median and 95% CI) from the nested dataframe into a regular dataframe 
    #(my_data_booted_summary) with 95%CI determined by "bca" or "perc" method
    booted_summary <- my_data_nested %>% mutate(G4P= purrr::map(.x = booted_ci, ~ .x$t0),
                                                lower_ci_bca = purrr::map(.x = booted_ci,~ .x$bca[[4]]),    
                                                upper_ci_bca = purrr::map(.x = booted_ci,~ .x$bca[[5]]),
                                                lower_ci_perc = purrr::map(.x = booted_ci,~ .x$perc[[4]]),
                                                upper_ci_perc = purrr::map(.x = booted_ci,~ .x$perc[[5]])) %>%  
        dplyr::select(-data, -booted, -booted_ci) %>% tidyr::unnest(cols = c(G4P, lower_ci_bca, upper_ci_bca, lower_ci_perc, upper_ci_perc))
    
    # Plot
    plot_not_normal(df, my_colours, booted_summary)
    
    # Stats
     
    kruskal_pval <- (df %>% group_by(Treatment)%>%kruskal_test(G4P ~ Line)) %>% select(Treatment, p)
    pval_dunn <- test_dunn()
    
    # Save plot
    ggsave("my_ggplot1.pdf", width=4, height=5)
    ggsave("my_ggplot1.svg", width=4, height=5)
}
```

##### 3.2. Results

###### a. Normality status

Table 1 : Shapiro test results.

| Treatment | Line | statistic | p.value |
| --- | --- | --- | --- |
| N50 | X1\_WT1 | 0.9524492 | 0.7314425 |
| N50 | X2\_OXRSH1 | 0.9982118 | 0.9944135 |
| N50 | X3\_WT2 | 0.8776720 | 0.3287883 |
| N50 | X4\_rsh1.1 | 0.9325955 | 0.6097369 |
| N50 | X5\_QM | 0.8527738 | 0.2352705 |

**Data follow a normal distribution. Now calculating means and confidence intervals for each line-treatment combination.**

###### b. Final data plot

**Fig 1. : Confidence interval plot**

###### c. Statistical analysis

###### Summary statistics

Table 2 : Statistical summary.

| Line | Treatment | N | G4P | sd | se | ci |
| --- | --- | --- | --- | --- | --- | --- |
| X1\_WT1 | N50 | 4 | 36.697 | 6.688 | 3.344 | 10.642 |
| X2\_OXRSH1 | N50 | 4 | 23.616 | 2.997 | 1.498 | 4.768 |
| X3\_WT2 | N50 | 4 | 41.950 | 2.129 | 1.065 | 3.388 |
| X4\_rsh1.1 | N50 | 4 | 60.088 | 7.646 | 3.823 | 12.167 |
| X5\_QM | N50 | 4 | 3.197 | 0.967 | 0.484 | 1.539 |

###### Statistical tests within each treatment condition.

Table 3 : Results of Anova test.

| Treatment | Effect | DFn | DFd | F | p | p<.05 | ges |
| --- | --- | --- | --- | --- | --- | --- | --- |
| N50 | Line | 4 | 15 | 76.663 | 8.44e-10 | \* | 0.953 |

Table 4 : Results of Tukey test.

| Treatment | group1 | group2 | p.adj | p.adj.signif |
| --- | --- | --- | --- | --- |
| N50 | X1\_WT1 | X2\_OXRSH1 | 1.25e-02 | \* |
| N50 | X1\_WT1 | X3\_WT2 | 5.59e-01 | ns |
| N50 | X1\_WT1 | X4\_rsh1.1 | 4.93e-05 | \*\*\*\* |
| N50 | X1\_WT1 | X5\_QM | 5.99e-07 | \*\*\*\* |
| N50 | X2\_OXRSH1 | X3\_WT2 | 6.67e-04 | \*\*\* |
| N50 | X2\_OXRSH1 | X4\_rsh1.1 | 1.96e-07 | \*\*\*\* |
| N50 | X2\_OXRSH1 | X5\_QM | 2.21e-04 | \*\*\* |
| N50 | X3\_WT2 | X4\_rsh1.1 | 7.41e-04 | \*\*\* |
| N50 | X3\_WT2 | X5\_QM | 8.70e-08 | \*\*\*\* |
| N50 | X4\_rsh1.1 | X5\_QM | 3.96e-10 | \*\*\*\* |

#### 4. R session information

```
InfoSession <- devtools::session_info()
print(InfoSession)
```

```
## - Session info ---------------------------------------------------------------
##  setting  value                       
##  version  R version 4.1.0 (2021-05-18)
##  os       Windows 10 x64              
##  system   x86_64, mingw32             
##  ui       RTerm                       
##  language (EN)                        
##  collate  English_United Kingdom.1252 
##  ctype    English_United Kingdom.1252 
##  tz       Europe/Paris                
##  date     2021-06-28                  
## 
## - Packages -------------------------------------------------------------------
##  package      * version date       lib source        
##  abind          1.4-5   2016-07-21 [1] CRAN (R 4.1.0)
##  assertthat     0.2.1   2019-03-21 [1] CRAN (R 4.1.0)
##  backports      1.2.1   2020-12-09 [1] CRAN (R 4.1.0)
##  beeswarm       0.4.0   2021-06-01 [1] CRAN (R 4.1.0)
##  boot         * 1.3-28  2021-05-03 [2] CRAN (R 4.1.0)
##  broom          0.7.7   2021-06-13 [1] CRAN (R 4.1.0)
##  cachem         1.0.5   2021-05-15 [1] CRAN (R 4.1.0)
##  callr          3.7.0   2021-04-20 [1] CRAN (R 4.1.0)
##  car            3.0-10  2020-09-29 [1] CRAN (R 4.1.0)
##  carData        3.0-4   2020-05-22 [1] CRAN (R 4.1.0)
##  cellranger     1.1.0   2016-07-27 [1] CRAN (R 4.1.0)
##  cli            2.5.0   2021-04-26 [1] CRAN (R 4.1.0)
##  colorspace     2.0-1   2021-05-04 [1] CRAN (R 4.1.0)
##  crayon         1.4.1   2021-02-08 [1] CRAN (R 4.1.0)
##  curl           4.3.1   2021-04-30 [1] CRAN (R 4.1.0)
##  data.table     1.14.0  2021-02-21 [1] CRAN (R 4.1.0)
##  DBI            1.1.1   2021-01-15 [1] CRAN (R 4.1.0)
##  desc           1.3.0   2021-03-05 [1] CRAN (R 4.1.0)
##  devtools     * 2.4.2   2021-06-07 [1] CRAN (R 4.1.0)
##  digest         0.6.27  2020-10-24 [1] CRAN (R 4.1.0)
##  dplyr        * 1.0.7   2021-06-18 [1] CRAN (R 4.1.0)
##  ellipsis       0.3.2   2021-04-29 [1] CRAN (R 4.1.0)
##  evaluate       0.14    2019-05-28 [1] CRAN (R 4.1.0)
##  fansi          0.5.0   2021-05-25 [1] CRAN (R 4.1.0)
##  farver         2.1.0   2021-02-28 [1] CRAN (R 4.1.0)
##  fastmap        1.1.0   2021-01-25 [1] CRAN (R 4.1.0)
##  forcats        0.5.1   2021-01-27 [1] CRAN (R 4.1.0)
##  foreign        0.8-81  2020-12-22 [2] CRAN (R 4.1.0)
##  fs             1.5.0   2020-07-31 [1] CRAN (R 4.1.0)
##  generics       0.1.0   2020-10-31 [1] CRAN (R 4.1.0)
##  ggbeeswarm   * 0.6.0   2017-08-07 [1] CRAN (R 4.1.0)
##  ggplot2      * 3.3.4   2021-06-16 [1] CRAN (R 4.1.0)
##  glue           1.4.2   2020-08-27 [1] CRAN (R 4.1.0)
##  gtable         0.3.0   2019-03-25 [1] CRAN (R 4.1.0)
##  haven          2.4.1   2021-04-23 [1] CRAN (R 4.1.0)
##  highr          0.9     2021-04-16 [1] CRAN (R 4.1.0)
##  hms            1.1.0   2021-05-17 [1] CRAN (R 4.1.0)
##  htmltools      0.5.1.1 2021-01-22 [1] CRAN (R 4.1.0)
##  knitr        * 1.33    2021-04-24 [1] CRAN (R 4.1.0)
##  labeling       0.4.2   2020-10-20 [1] CRAN (R 4.1.0)
##  lattice      * 0.20-44 2021-05-02 [2] CRAN (R 4.1.0)
##  lifecycle      1.0.0   2021-02-15 [1] CRAN (R 4.1.0)
##  magrittr     * 2.0.1   2020-11-17 [1] CRAN (R 4.1.0)
##  memoise        2.0.0   2021-01-26 [1] CRAN (R 4.1.0)
##  munsell        0.5.0   2018-06-12 [1] CRAN (R 4.1.0)
##  openxlsx       4.2.4   2021-06-16 [1] CRAN (R 4.1.0)
##  pillar         1.6.1   2021-05-16 [1] CRAN (R 4.1.0)
##  pkgbuild       1.2.0   2020-12-15 [1] CRAN (R 4.1.0)
##  pkgconfig      2.0.3   2019-09-22 [1] CRAN (R 4.1.0)
##  pkgload        1.2.1   2021-04-06 [1] CRAN (R 4.1.0)
##  plyr         * 1.8.6   2020-03-03 [1] CRAN (R 4.1.0)
##  prettyunits    1.1.1   2020-01-24 [1] CRAN (R 4.1.0)
##  processx       3.5.2   2021-04-30 [1] CRAN (R 4.1.0)
##  ps             1.6.0   2021-02-28 [1] CRAN (R 4.1.0)
##  purrr        * 0.3.4   2020-04-17 [1] CRAN (R 4.1.0)
##  R6             2.5.0   2020-10-28 [1] CRAN (R 4.1.0)
##  RColorBrewer * 1.1-2   2014-12-07 [1] CRAN (R 4.1.0)
##  Rcpp           1.0.6   2021-01-15 [1] CRAN (R 4.1.0)
##  readxl       * 1.3.1   2019-03-13 [1] CRAN (R 4.1.0)
##  remotes        2.4.0   2021-06-02 [1] CRAN (R 4.1.0)
##  rio            0.5.27  2021-06-21 [1] CRAN (R 4.1.0)
##  rlang          0.4.11  2021-04-30 [1] CRAN (R 4.1.0)
##  rmarkdown      2.9     2021-06-15 [1] CRAN (R 4.1.0)
##  Rmisc        * 1.5     2013-10-22 [1] CRAN (R 4.1.0)
##  rprojroot      2.0.2   2020-11-15 [1] CRAN (R 4.1.0)
##  rstatix      * 0.7.0   2021-02-13 [1] CRAN (R 4.1.0)
##  scales         1.1.1   2020-05-11 [1] CRAN (R 4.1.0)
##  sessioninfo    1.1.1   2018-11-05 [1] CRAN (R 4.1.0)
##  stringi        1.6.1   2021-05-10 [1] CRAN (R 4.1.0)
##  stringr        1.4.0   2019-02-10 [1] CRAN (R 4.1.0)
##  svglite      * 2.0.0   2021-02-20 [1] CRAN (R 4.1.0)
##  systemfonts    1.0.2   2021-05-11 [1] CRAN (R 4.1.0)
##  testthat       3.0.3   2021-06-16 [1] CRAN (R 4.1.0)
##  tibble         3.1.2   2021-05-16 [1] CRAN (R 4.1.0)
##  tidyr        * 1.1.3   2021-03-03 [1] CRAN (R 4.1.0)
##  tidyselect     1.1.1   2021-04-30 [1] CRAN (R 4.1.0)
##  usethis      * 2.0.1   2021-02-10 [1] CRAN (R 4.1.0)
##  utf8           1.2.1   2021-03-12 [1] CRAN (R 4.1.0)
##  vctrs          0.3.8   2021-04-29 [1] CRAN (R 4.1.0)
##  vipor          0.4.5   2017-03-22 [1] CRAN (R 4.1.0)
##  withr          2.4.2   2021-04-18 [1] CRAN (R 4.1.0)
##  xfun           0.24    2021-06-15 [1] CRAN (R 4.1.0)
##  yaml           2.2.1   2020-02-01 [1] CRAN (R 4.1.0)
##  zip            2.2.0   2021-05-31 [1] CRAN (R 4.1.0)
## 
## [1] C:/Users/Ben/Documents/R/win-library/4.1
## [2] C:/Program Files/R/R-4.1.0/library
```

#### 5. Citations

1. Bache, Stefan Milton and Wickham, Hadley (2020). magrittr: A Forward-Pipe Operator for R. R package version 2.0.1. https://CRAN.R-project.org/package=magrittr
2. Canty, Angelo and Ripley, Brian (2021). boot: Bootstrap R (S-Plus) Functions. R package version 1.3-28.
3. Clarke, Erik, and Scott Sherrill-Mix. 2017. Ggbeeswarm: Categorical Scatter (Violin Point) Plots. https://CRAN.R-project.org/package=ggbeeswarm.
4. Henry, Lionel and Wickham, Hadley (2020). purrr: Functional Programming Tools. R package version 0.3.4. https://CRAN.R-project.org/package=purrr
5. Hope, Ryan M. 2013. Rmisc: Rmisc: Ryan Miscellaneous. https://CRAN.R-project.org/package=Rmisc.
6. Kassambara, Alboukadel. 2021. Rstatix: Pipe-Friendly Framework for Basic Statistical Tests. https://CRAN.R-project.org/package=rstatix.
7. Neuwirth, Erich. 2014. RColorBrewer: ColorBrewer Palettes. https://CRAN.R-project.org/package=RColorBrewer.
8. R Core Team. 2020. R: A Language and Environment for Statistical Computing. Vienna, Austria: R Foundation for Statistical Computing. https://www.R-project.org/.
9. Wickham, Hadley. 2016. Ggplot2: Elegant Graphics for Data Analysis. Springer-Verlag New York. https://ggplot2.tidyverse.org.
10. Wickham, Hadley, and Jennifer Bryan. 2019. Readxl: Read Excel Files. https://CRAN.R-project.org/package=readxl.
11. Wickham, Hadley, Romain François, Lionel Henry, and Kirill Müller. 2021. Dplyr: A Grammar of Data Manipulation. https://CRAN.R-project.org/package=dplyr.
12. Wickham, Hadley (2021). tidyr: Tidy Messy Data. R package version 1.1.3. https://CRAN.R-project.org/package=tidyr
13. Wickham, Hadley, Hester, Jim and Chang, Winston (2021). devtools: Tools to Make Developing R Packages Easier. R package version 2.4.2. https://CRAN.R-project.org/package=devtools
