## Supplementary material for "A ppGpp-mediated brake on photosynthesis is required for acclimation to nitrogen limitation in Arabidopsis": R scripts: script_ci_FvFm.html

Plot and statistical analysis


### Plot and statistical analysis

#### ----

#### 1. Introduction

The analysis reported here is part of the manuscript Romand et al. 2021. The script performs statistical analysis on data grouped by `Treatment` and `Plant Line` and plots a graph. Different approaches are used depending on the normality of the data as determined by the Shapiro test. If data follow a normal distribution, then the script performs Anova and post-hoc Tukey tests and draws a plot with confidence interval centered on mean. If data do not follow a normal distribution then a Kruskal-Wallis and a pot-hoc Dunn test are performed. The same plot is drawn using the medians and boostrap calculated 95% confidence intervals. The plots are also saved as pdf and vector (SVG) files.

#### 2. Data import

The script will analyse a data file placed in the working directory with the following layout.

A 3 column data set in `xlsx`. Column 1, `Line`; column 2, `Treatment` and column 3 the measured quantity, in this case Quantum Yield (FvFm). The 2 first columns are factors that are used to group the data.

In this example we have 2 levels “N” and “NoN” for Treatment and 3 levels “WT2”, “QM”, “rsh1-1” for Line.

This script can be simply modified to analyse different parameters or different treatments. It can also be modified to analyse data grouped by additional factors such as Time.

#### 3. Data analysis

##### 3.1. Code

###### Set variables for personalisation

```
#1. Data
#------------------------------------------------------------------------------

# Put your input file in variable DATA 
DATA <- "QY.xlsx"


#2. Parameters for plot
#------------------------------------------------------------------------------
# Points color
COLOUR <- "#00B050"

# Select a palette for CI color
PALETTE <- "PuRd"
```

Working directory: C:/Users/Ben Field/AMUBOX/Shared/ppGpp\_nitrogen\_starvation/model markdown/markdown CI QY

###### Setup R packages

```
if (!require(gridExtra)) { install.packages("gridExtra", repos = "http://cran.us.r-project.org") }
if (!require(ggbeeswarm)) { install.packages("ggbeeswarm", repos = "http://cran.us.r-project.org") }
if (!require(Rmisc)) { install.packages("Rmisc", repos = "http://cran.us.r-project.org") }
if (!require(ggplot2)) { install.packages("ggplot2", repos = "http://cran.us.r-project.org") }
if (!require(RColorBrewer)) { install.packages("RColorBrewer", repos = "http://cran.us.r-project.org") }
if (!require(dplyr)) { install.packages("dplyr", repos = "http://cran.us.r-project.org") }
if (!require(rstatix)) { install.packages("rstatix", repos = "https://cloud.r-project.org") }
if (!require(purrr)) { install.packages("purrr", repos = "http://cran.us.r-project.org") }
if (!require(tidyr)) { install.packages("tidyr", repos = "http://cran.us.r-project.org") }
if (!require(magrittr)) { install.packages("magrittr", repos = "http://cran.us.r-project.org") }
if (!require(boot)) { install.packages("boot", repos = "http://cran.us.r-project.org") }
if (!require(devtools)) { install.packages("devtools", repos = "http://cran.us.r-project.org") }
if (!require(knitr)) { install.packages("knitr", repos = "http://cran.us.r-project.org") }
if (!require(svglite)) { install.packages("svglite", repos = "http://cran.us.r-project.org") }

library(gridExtra)
library(ggplot2)
library(ggbeeswarm)
library(Rmisc)
library(readxl)
library(RColorBrewer)
library(rstatix)
library(gridExtra)
library(boot)
library(magrittr)
library(dplyr)
library(purrr)
library(tidyr)
library(rstatix)
library(knitr)
library(svglite)
```

###### Functions used in the script

```
#===============================================================================
# Draw the plot if data follow a normal distribution
#===============================================================================

plot_normal <- function(df, my_colours, my_summary) {
    p<-ggplot(data = df, aes(x=Line, y=FvFm)) +
        geom_quasirandom(dodge.width=0.8,alpha = 0.6, colour = COLOUR)+
        geom_pointrange(data = my_summary, aes(ymin=FvFm-ci, ymax=FvFm+ci, color=Line), position=position_dodge(width=0.8))+
        scale_colour_manual(values=my_colours)+
        scale_y_continuous(breaks=seq(0,0.8,0.2), limits=c(0,0.95),expand = c(0, 0))+
        facet_wrap(~Treatment, strip.position = "bottom", scales = "free_x") +
        theme_classic() + 
        theme(panel.spacing = unit(0, "lines"), 
              strip.background = element_blank(),
              strip.placement = "outside")+
        theme(axis.text.x = element_text(angle = 90, vjust = 0.5, hjust=1))
    print(p)
}

#===============================================================================
# Draw the plot if data do not follow a normal distribution
#===============================================================================

plot_not_normal <- function(df, my_colours, conf_int) {
    p<-ggplot(data=df, aes(x=Line, y=FvFm)) +
        geom_quasirandom(dodge.width=0.8,alpha = 0.6, colour = COLOUR)+
        geom_linerange(data = booted_summary, aes(ymin=lower_ci_perc, ymax=upper_ci_perc, color=Line), position = position_dodge(width=0.8))+
        geom_point(data=booted_summary, aes(y=FvFm, color=Line), size = 2, 
                   position=position_dodge(width=0.8))+
        scale_colour_manual(values=my_colours)+
        scale_y_continuous(breaks=seq(0,0.8,0.2), limits=c(0,0.95),expand = c(0, 0))+
        facet_wrap(~Treatment, strip.position = "bottom", scales = "free_x") +
        theme_classic() + 
        theme(panel.spacing = unit(0, "lines"), 
              strip.background = element_blank(),
              strip.placement = "outside")+
        theme(axis.text.x = element_text(angle = 90, vjust = 0.5, hjust=1))
        print(p)
       }

#===============================================================================
# Check normality of the data. Exit the loop if one set of data doesn't follow a 
# normal distribution.
# Return TRUE if all data follow a normal distribution
#===============================================================================

check_normality <- function(shapiro_df) {
    # Data are supposed to follow normal distribution
    flag_normal <- TRUE
    
    for (i in 1 : nrow(shapiro_df)) {
        if(shapiro_df[i, 4] > 0.05) {
            
        } else {
            
            flag_normal <- FALSE
            break
        }
    }
    return(flag_normal)
}

#===============================================================================
#Function to calculate the median (for bootstrapping)
#===============================================================================

samplemedian <- function(d, i) {
    median(d[i])
}

#===============================================================================
#Function for Dunn test
#===============================================================================
test_dunn <- function() {
    pval <- as.data.frame(df %>% group_by(Treatment) %>% dunn_test(FvFm ~ Line, p.adjust.method = "BY"))
    print(df %>% group_by(Treatment) %>% dunn_test(FvFm ~ Line, p.adjust.method = "BY"))
    return(pval)
}
```

###### Main script

```
# Import Data
df <- read_excel(DATA, col_types = c("text", "text", "numeric"))

# Define color
my_colours = brewer.pal(n = 9, PALETTE)[9:3]

# Determine data normality using the Shapiro test
shapiro_df <- df %>%group_by(Treatment, Line)%>%
    summarise(statistic = shapiro.test(FvFm)$statistic, 
              p.value = shapiro.test(FvFm)$p.value)

flag_normal <- check_normality(shapiro_df)


# Data treatement according to normality status
if(flag_normal == TRUE) {
    #print("Data follow a normal distribution")
    
    # Summary
    my_summary <- summarySE(df, measurevar="FvFm", groupvars=c("Line", "Treatment"))
    
    # Plot
    plot_normal(df, my_colours, my_summary)
    
    # Stats
    anova_results <- df %>% group_by(Treatment) %>%  anova_test(FvFm ~ Line)
    
    tukey_results <- df %>% group_by(Treatment) %>%  tukey_hsd(FvFm ~ Line)
    
    # Save plot
    ggsave("my_ggplot1.pdf", width=4, height=5)
    ggsave("my_ggplot1.svg", width=4, height=5)
    
} else {
    #print("Data do not follow a normal distribution")
    
    # Calculate Median and 95%CI by bootstrap (Largely based on code by Peter Kamerman: 
    #https://www.painblogr.org/2017-10-18-purrring-through-bootstraps.html)
    
    #1. Set the seed
    set.seed(12345)
    
    #2. Create nested datafile to allow boostrapping on each subgroup.
    my_data_nested <- df %>% group_by(Line,Treatment) %>% nest()
    
    #3. Add boostrapped median values to my_nested_data
    my_data_nested %<>% dplyr::mutate(booted = purrr::map(.x = data, ~ boot::boot(data = .x$FvFm,statistic = samplemedian, R = 10000, stype = "i")))
    
    #4. Plot each <S3: boot> object to check boostrapping worked!
    ## Note: Saved to an object (plots) to stop the summary being printed
    plots<-purrr::map(.x = my_data_nested$booted, ~ plot(.x))
    print(plots)
    
    #5. Add boostrapped 95%CI around median values to my_nested_data, 
    #95%CI determined based on "perc" & "bca"
    my_data_nested %<>% mutate(booted_ci = purrr::map(.x = booted, 
                                                      ~ boot::boot.ci(.x,conf = 0.95, 
                                                                      type = c("perc", "bca"))))
    
    #6. Table of each <S3: bootci> object to check
    ci_table <-purrr::map(.x = my_data_nested$booted_ci, ~ print(.x))
    
    #7. Get the stats (median and 95% CI) from the nested dataframe into a regular dataframe 
    #(my_data_booted_summary) with 95%CI determined by "bca" or "perc" method
    booted_summary <- my_data_nested %>% mutate(FvFm= purrr::map(.x = booted_ci, ~ .x$t0),
                                                lower_ci_bca = purrr::map(.x = booted_ci,~ .x$bca[[4]]),    
                                                upper_ci_bca = purrr::map(.x = booted_ci,~ .x$bca[[5]]),
                                                lower_ci_perc = purrr::map(.x = booted_ci,~ .x$perc[[4]]),
                                                upper_ci_perc = purrr::map(.x = booted_ci,~ .x$perc[[5]])) %>%  
        dplyr::select(-data, -booted, -booted_ci) %>% tidyr::unnest(cols = c(FvFm, lower_ci_bca, upper_ci_bca, lower_ci_perc, upper_ci_perc))
    
    # Plot
    plot_not_normal(df, my_colours, booted_summary)
    
    # Stats
    
    kruskal_pval <- (df %>% group_by(Treatment)%>%kruskal_test(FvFm ~ Line)) %>% select(Treatment, p)
    pval_dunn <- test_dunn()
    
       # Save plot
    ggsave("my_ggplot1.pdf", width=4, height=5)
    ggsave("my_ggplot1.svg", width=4, height=5)
}
```

##### 3.2. Results

###### a. Normality test and summary statistics

Table 1 : Shapiro test results.

| Treatment | Line | statistic | p.value |
| --- | --- | --- | --- |
| N | 01\_WT2 | 0.9493549 | 3.410084e-03 |
| N | 02\_rsh1-1 | 0.9183088 | 1.462245e-04 |
| N | 03\_QM | 0.9555055 | 1.011324e-02 |
| No\_N | 01\_WT2 | 0.8687690 | 7.912134e-08 |
| No\_N | 02\_rsh1-1 | 0.9081497 | 3.850902e-06 |
| No\_N | 03\_QM | 0.8853211 | 3.069355e-07 |

**Data do not follow a normal distribution. Now calculating medians and confidence intervals by a boostrap procedure for each line-treatment combination.**

**Fig 1. : Plots of bootstrapping process.**

###### b. Final data plot

**Fig 2. : Confidence interval plot**

###### c. Statistical analysis

###### Summary statistics

Table 2 : Statistical summary.

| Line | Treatment | FvFm | lower\_ci\_bca | upper\_ci\_bca | lower\_ci\_perc | upper\_ci\_perc |
| --- | --- | --- | --- | --- | --- | --- |
| 01\_WT2 | N | 0.8404579 | 0.8375365 | 0.8419380 | 0.8376947 | 0.8420788 |
| 02\_rsh1-1 | N | 0.8382673 | 0.8357758 | 0.8425856 | 0.8357827 | 0.8425870 |
| 03\_QM | N | 0.8430361 | 0.8415357 | 0.8448404 | 0.8418465 | 0.8449285 |
| 01\_WT2 | No\_N | 0.6680693 | 0.6097395 | 0.7125317 | 0.6133099 | 0.7140256 |
| 02\_rsh1-1 | No\_N | 0.2865251 | 0.1844457 | 0.3553825 | 0.1844457 | 0.3577764 |
| 03\_QM | No\_N | 0.7711883 | 0.7633340 | 0.7768576 | 0.7633883 | 0.7773351 |

###### Statistical tests within each treatment condition.

Table 3 : Results of Kruskal-Wallis test.

| Treatment | p |
| --- | --- |
| N | 1.35e-02 |
| No\_N | 7.01e-44 |

Table 4 : Results of Dunn test.

| Treatment | group1 | group2 | p | p.adj | p.adj.signif |
| --- | --- | --- | --- | --- | --- |
| N | 01\_WT2 | 02\_rsh1-1 | 8.634118e-01 | 1.000000e+00 | ns |
| N | 01\_WT2 | 03\_QM | 1.324332e-02 | 3.641913e-02 | \* |
| N | 02\_rsh1-1 | 03\_QM | 9.127639e-03 | 3.641913e-02 | \* |
| No\_N | 01\_WT2 | 02\_rsh1-1 | 1.636479e-13 | 4.500318e-13 | \*\*\*\* |
| No\_N | 01\_WT2 | 03\_QM | 2.698707e-11 | 4.947630e-11 | \*\*\*\* |
| No\_N | 02\_rsh1-1 | 03\_QM | 4.246754e-45 | 2.335715e-44 | \*\*\*\* |

#### 4. R session information

```
InfoSession <- devtools::session_info()
print(InfoSession)
```

```
## - Session info ---------------------------------------------------------------
##  setting  value                       
##  version  R version 4.1.0 (2021-05-18)
##  os       Windows 10 x64              
##  system   x86_64, mingw32             
##  ui       RTerm                       
##  language (EN)                        
##  collate  English_United States.1252  
##  ctype    English_United States.1252  
##  tz       Europe/Paris                
##  date     2021-07-05                  
## 
## - Packages -------------------------------------------------------------------
##  package      * version date       lib source        
##  abind          1.4-5   2016-07-21 [1] CRAN (R 4.1.0)
##  assertthat     0.2.1   2019-03-21 [1] CRAN (R 4.1.0)
##  backports      1.2.1   2020-12-09 [1] CRAN (R 4.1.0)
##  beeswarm       0.4.0   2021-06-01 [1] CRAN (R 4.1.0)
##  boot         * 1.3-28  2021-05-03 [2] CRAN (R 4.1.0)
##  broom          0.7.8   2021-06-24 [1] CRAN (R 4.1.0)
##  cachem         1.0.5   2021-05-15 [1] CRAN (R 4.1.0)
##  callr          3.7.0   2021-04-20 [1] CRAN (R 4.1.0)
##  car            3.0-11  2021-06-27 [1] CRAN (R 4.1.0)
##  carData        3.0-4   2020-05-22 [1] CRAN (R 4.1.0)
##  cellranger     1.1.0   2016-07-27 [1] CRAN (R 4.1.0)
##  cli            2.5.0   2021-04-26 [1] CRAN (R 4.1.0)
##  colorspace     2.0-2   2021-06-24 [1] CRAN (R 4.1.0)
##  crayon         1.4.1   2021-02-08 [1] CRAN (R 4.1.0)
##  curl           4.3.1   2021-04-30 [1] CRAN (R 4.1.0)
##  data.table     1.14.0  2021-02-21 [1] CRAN (R 4.1.0)
##  DBI            1.1.1   2021-01-15 [1] CRAN (R 4.1.0)
##  desc           1.3.0   2021-03-05 [1] CRAN (R 4.1.0)
##  devtools     * 2.4.2   2021-06-07 [1] CRAN (R 4.1.0)
##  digest         0.6.27  2020-10-24 [1] CRAN (R 4.1.0)
##  dplyr        * 1.0.7   2021-06-18 [1] CRAN (R 4.1.0)
##  ellipsis       0.3.2   2021-04-29 [1] CRAN (R 4.1.0)
##  evaluate       0.14    2019-05-28 [1] CRAN (R 4.1.0)
##  fansi          0.5.0   2021-05-25 [1] CRAN (R 4.1.0)
##  farver         2.1.0   2021-02-28 [1] CRAN (R 4.1.0)
##  fastmap        1.1.0   2021-01-25 [1] CRAN (R 4.1.0)
##  forcats        0.5.1   2021-01-27 [1] CRAN (R 4.1.0)
##  foreign        0.8-81  2020-12-22 [2] CRAN (R 4.1.0)
##  fs             1.5.0   2020-07-31 [1] CRAN (R 4.1.0)
##  generics       0.1.0   2020-10-31 [1] CRAN (R 4.1.0)
##  ggbeeswarm   * 0.6.0   2017-08-07 [1] CRAN (R 4.1.0)
##  ggplot2      * 3.3.5   2021-06-25 [1] CRAN (R 4.1.0)
##  glue           1.4.2   2020-08-27 [1] CRAN (R 4.1.0)
##  gridExtra    * 2.3     2017-09-09 [1] CRAN (R 4.1.0)
##  gtable         0.3.0   2019-03-25 [1] CRAN (R 4.1.0)
##  haven          2.4.1   2021-04-23 [1] CRAN (R 4.1.0)
##  highr          0.9     2021-04-16 [1] CRAN (R 4.1.0)
##  hms            1.1.0   2021-05-17 [1] CRAN (R 4.1.0)
##  htmltools      0.5.1.1 2021-01-22 [1] CRAN (R 4.1.0)
##  knitr        * 1.33    2021-04-24 [1] CRAN (R 4.1.0)
##  lattice      * 0.20-44 2021-05-02 [2] CRAN (R 4.1.0)
##  lifecycle      1.0.0   2021-02-15 [1] CRAN (R 4.1.0)
##  magrittr     * 2.0.1   2020-11-17 [1] CRAN (R 4.1.0)
##  memoise        2.0.0   2021-01-26 [1] CRAN (R 4.1.0)
##  munsell        0.5.0   2018-06-12 [1] CRAN (R 4.1.0)
##  openxlsx       4.2.4   2021-06-16 [1] CRAN (R 4.1.0)
##  pillar         1.6.1   2021-05-16 [1] CRAN (R 4.1.0)
##  pkgbuild       1.2.0   2020-12-15 [1] CRAN (R 4.1.0)
##  pkgconfig      2.0.3   2019-09-22 [1] CRAN (R 4.1.0)
##  pkgload        1.2.1   2021-04-06 [1] CRAN (R 4.1.0)
##  plyr         * 1.8.6   2020-03-03 [1] CRAN (R 4.1.0)
##  prettyunits    1.1.1   2020-01-24 [1] CRAN (R 4.1.0)
##  processx       3.5.2   2021-04-30 [1] CRAN (R 4.1.0)
##  ps             1.6.0   2021-02-28 [1] CRAN (R 4.1.0)
##  purrr        * 0.3.4   2020-04-17 [1] CRAN (R 4.1.0)
##  R6             2.5.0   2020-10-28 [1] CRAN (R 4.1.0)
##  RColorBrewer * 1.1-2   2014-12-07 [1] CRAN (R 4.1.0)
##  Rcpp           1.0.6   2021-01-15 [1] CRAN (R 4.1.0)
##  readxl       * 1.3.1   2019-03-13 [1] CRAN (R 4.1.0)
##  remotes        2.4.0   2021-06-02 [1] CRAN (R 4.1.0)
##  rio            0.5.27  2021-06-21 [1] CRAN (R 4.1.0)
##  rlang          0.4.11  2021-04-30 [1] CRAN (R 4.1.0)
##  rmarkdown      2.9     2021-06-15 [1] CRAN (R 4.1.0)
##  Rmisc        * 1.5     2013-10-22 [1] CRAN (R 4.1.0)
##  rprojroot      2.0.2   2020-11-15 [1] CRAN (R 4.1.0)
##  rstatix      * 0.7.0   2021-02-13 [1] CRAN (R 4.1.0)
##  rstudioapi     0.13    2020-11-12 [1] CRAN (R 4.1.0)
##  scales         1.1.1   2020-05-11 [1] CRAN (R 4.1.0)
##  sessioninfo    1.1.1   2018-11-05 [1] CRAN (R 4.1.0)
##  stringi        1.6.1   2021-05-10 [1] CRAN (R 4.1.0)
##  stringr        1.4.0   2019-02-10 [1] CRAN (R 4.1.0)
##  svglite      * 2.0.0   2021-02-20 [1] CRAN (R 4.1.0)
##  systemfonts    1.0.2   2021-05-11 [1] CRAN (R 4.1.0)
##  testthat       3.0.3   2021-06-16 [1] CRAN (R 4.1.0)
##  tibble         3.1.2   2021-05-16 [1] CRAN (R 4.1.0)
##  tidyr        * 1.1.3   2021-03-03 [1] CRAN (R 4.1.0)
##  tidyselect     1.1.1   2021-04-30 [1] CRAN (R 4.1.0)
##  usethis      * 2.0.1   2021-02-10 [1] CRAN (R 4.1.0)
##  utf8           1.2.1   2021-03-12 [1] CRAN (R 4.1.0)
##  vctrs          0.3.8   2021-04-29 [1] CRAN (R 4.1.0)
##  vipor          0.4.5   2017-03-22 [1] CRAN (R 4.1.0)
##  withr          2.4.2   2021-04-18 [1] CRAN (R 4.1.0)
##  xfun           0.24    2021-06-15 [1] CRAN (R 4.1.0)
##  yaml           2.2.1   2020-02-01 [1] CRAN (R 4.1.0)
##  zip            2.2.0   2021-05-31 [1] CRAN (R 4.1.0)
## 
## [1] C:/Users/Ben Field/Documents/R/win-library/4.1
## [2] C:/Program Files/R/R-4.1.0/library
```

#### 5. Citations

1. Bache, Stefan Milton and Wickham, Hadley (2020). magrittr: A Forward-Pipe Operator for R. R package version 2.0.1. https://CRAN.R-project.org/package=magrittr
2. Canty, Angelo and Ripley, Brian (2021). boot: Bootstrap R (S-Plus) Functions. R package version 1.3-28.
3. Clarke, Erik, and Scott Sherrill-Mix. 2017. Ggbeeswarm: Categorical Scatter (Violin Point) Plots. https://CRAN.R-project.org/package=ggbeeswarm.
4. Henry, Lionel and Wickham, Hadley (2020). purrr: Functional Programming Tools. R package version 0.3.4. https://CRAN.R-project.org/package=purrr
5. Hope, Ryan M. 2013. Rmisc: Rmisc: Ryan Miscellaneous. https://CRAN.R-project.org/package=Rmisc.
6. Kassambara, Alboukadel. 2021. Rstatix: Pipe-Friendly Framework for Basic Statistical Tests. https://CRAN.R-project.org/package=rstatix.
7. Neuwirth, Erich. 2014. RColorBrewer: ColorBrewer Palettes. https://CRAN.R-project.org/package=RColorBrewer.
8. R Core Team. 2020. R: A Language and Environment for Statistical Computing. Vienna, Austria: R Foundation for Statistical Computing. https://www.R-project.org/.
9. Wickham, Hadley. 2016. Ggplot2: Elegant Graphics for Data Analysis. Springer-Verlag New York. https://ggplot2.tidyverse.org.
10. Wickham, Hadley, and Jennifer Bryan. 2019. Readxl: Read Excel Files. https://CRAN.R-project.org/package=readxl.
11. Wickham, Hadley, Romain François, Lionel Henry, and Kirill Müller. 2021. Dplyr: A Grammar of Data Manipulation. https://CRAN.R-project.org/package=dplyr.
12. Wickham, Hadley (2021). tidyr: Tidy Messy Data. R package version 1.1.3. https://CRAN.R-project.org/package=tidyr
13. Wickham, Hadley, Hester, Jim and Chang, Winston (2021). devtools: Tools to Make Developing R Packages Easier. R package version 2.4.2. https://CRAN.R-project.org/package=devtools
