## Supplementary material for "A ppGpp-mediated brake on photosynthesis is required for acclimation to nitrogen limitation in Arabidopsis": R scripts: R-markdown-GO.html

Report of Gene Ontology Analysis


### Report of Gene Ontology Analysis

#### ----

#### 1. Introduction

The analysis reported here is part of the project “G4PLAST”. The aim is to find gene ontology enrichment in lists of differentially expressed genes. Preparation of the data is performed by the `prepare_gene_ontology.pl` script (Terese et Lecampion : github).

The gene ontology analysis and plot described here are based on Bonnot *et al.* (A Simple Protocol for Informative Visualization of Enriched Gene Ontology Terms. T. Bonnot, MB. Gillard and DH. Nagel. Bio-101: e3429. DOI:10.21769/BioProtoc.3429).

#### 2. Data

The data used here are lists of differentially expressed genes. The “up” or “down” tables supplied by SARTools (Varet *et al.* 2016) are directly used by the the `prepare_gene_ontology.pl` script which automatically uses PANTHER and REVIGO for the identification and simplification of enriched GO terms using the first column of gene ids (according ot the manual approach described by Bonnot et al.). See Readme on github for more information.

Table 1: Partial view of the data

| Id | wt\_ms2\_1 | wt\_ms2\_2 | wt\_ms2\_3 | wt\_n50\_1 | wt\_n50\_2 | … |
| --- | --- | --- | --- | --- | --- | --- |
| AT1G34060.1 | 19 | 21 | 20 | 16786 | 11823 |  |
| AT1G66390.1 | 9 | 6 | 2 | 10674 | 9721 |  |
| AT2G14560.3 | 14 | 27 | 27 | 7907 | 8130 |  |

The `prepare_gene_ontology.pl` script produces \*.tsv files that are then used by this R markdown script to produce a publication quality plot. It is possible to use biological process (BP), cellular compartment (CC) or molecular function (MF) lists.

Working directory

Working directory: C:/Users/Ben Field/AMUBOX/Shared/ppGpp\_nitrogen\_starvation/Supp data sets/R scripts/R markdown GO analysis Fig\_4

#### 3. Graphic representation of the results

Install needed packages

```
if (!requireNamespace("ggplot2", quietly = TRUE))
    install.packages("ggplot2")
if (!requireNamespace("cowplot", quietly = TRUE))
    install.packages("cowplot")
if (!requireNamespace("devtools", quietly = TRUE))
    install.packages("devtools")

library(ggplot2)     
library(cowplot)
```

Load the data and select number of GO terms to include in plot (data are sorted on FDR)

```
GO_up <- read.table("OXRSH1vsWT.up_GO_output_BP.tsv",header=T,stringsAsFactors = T)     
GO_down <- read.table("OXRSH1vsWT.down_GO_output_BP.tsv",header=T,stringsAsFactors = T)

# reduce the number of GO ID used
GO_up <- GO_up[1:10,]   
GO_down <- GO_down[1:10,]
```

Create the plots

```
#For the GO_up
# List objects and their structure contained in the dataframe 'GO_all'
ls.str(GO_up)
```

```
## FDR :  num [1:10] 2.06e-62 2.60e-62 3.90e-62 8.99e-62 9.91e-57 ...
## Fold_enrichment :  num [1:10] 4.7 4.68 4.68 4.63 3.79 2.11 2.75 2.59 4.79 4.82
## Gene_number :  int [1:10] 191 190 190 190 213 434 274 298 147 146
## GO_id :  Factor w/ 296 levels "activation_of_immune_response_(GO:0002253)",..: 225 256 234 117 235 264 227 265 104 103
```

```
# Transform the column 'Gene_number' into a numeric variable
GO_up$Gene_number <- as.numeric(GO_up$Gene_number)

# Replace all the "_" by a space in the column containing the GO terms
GO_up$GO_id <- chartr("_", " ", GO_up$GO_id)

# Transform FDR values by -log10('FDR values')
GO_up$'|log10(FDR)|' <- -(log10(GO_up$FDR))

# Create a ggplot graph without printing it
up <- ggplot(GO_up, aes(x = GO_id, y = Fold_enrichment)) +
  geom_hline(yintercept = 1, color = "azure4", size=.5, alpha=0.7)+
  geom_point(data=GO_up, aes(x=GO_id, y=Fold_enrichment, size = Gene_number, colour = `|log10(FDR)|`), alpha=.7)+
  scale_size_continuous(range = c(4, 15))+  # Sets min and max point size
  scale_y_continuous(limits = c(1,13), breaks = c(1,4,8,12))+
  scale_x_discrete(limits= GO_up$GO_id)+
  scale_colour_gradient(low="blue", high="red", limits=c(0, 90))+
  coord_flip()+
  theme_classic()+
  ylab("Fold enrichment")+
  ggtitle("UP")+
  labs(color="-log(FDR)", size="Number\nof genes")+ #Replace by your variable names; \n allow a new line for text
  guides(size = "none",color = guide_colourbar(ticks = FALSE, barwidth = 0.5))

#For the GO_down
# List objects and their structure contained in the dataframe 'GO_all'
ls.str(GO_down)
```

```
## FDR :  num [1:10] 2.84e-39 2.73e-37 1.83e-31 1.95e-31 1.75e-30 ...
## Fold_enrichment :  num [1:10] 10.04 2.72 11.2 4.16 4.03 ...
## Gene_number :  int [1:10] 73 226 55 115 115 654 493 222 72 227
## GO_id :  Factor w/ 263 levels "alcohol_biosynthetic_process_(GO:0046165)",..: 156 203 155 223 230 40 116 144 83 9
```

```
# Transform the column 'Gene_number' into a numeric variable
GO_down$Gene_number <- as.numeric(GO_down$Gene_number)

# Replace all the "_" by a space in the column containing the GO terms
GO_down$GO_id <- chartr("_", " ", GO_down$GO_id)

# Transform FDR values by -log10('FDR values')
GO_down$'|log10(FDR)|' <- -(log10(GO_down$FDR))

# Create a ggplot graph without printing it
down <- ggplot(GO_down, aes(x = GO_id, y = Fold_enrichment)) +
  geom_hline(yintercept = 1, color = "azure4", size=.5, alpha=0.7)+
  geom_point(data=GO_down, aes(x=GO_id, y=Fold_enrichment, size = Gene_number, colour = `|log10(FDR)|`), alpha=.7)+
  scale_size_continuous(range = c(4, 15))+  # Sets min and max point size
  scale_y_continuous(limits = c(1,13),breaks = c(1,4,8,12))+
  scale_x_discrete(limits= GO_down$GO_id)+
  scale_color_gradient(low="blue", high="red", limits=c(0, 90))+
  coord_flip()+
  theme_classic()+
  ylab("Fold enrichment")+
  ggtitle("DOWN")+
  labs(color="-log(FDR)", size="Number\nof genes")+ #Replace by your variable names; \n allow a new line for text
  guides(size = "none",color = guide_colourbar(ticks = FALSE, barwidth = 0.5))
```

Combine the 2 graphs into one

```
cowplot::plot_grid(up, down, ncol = 1, align = "v")
```

```
ggsave("GO.pdf",width = 10, height = 6)
ggsave("GO.svg",width = 10, height = 6)
```

#### 5. R session information and parameters

```
devtools::session_info()
```

```
## - Session info ---------------------------------------------------------------
##  setting  value                       
##  version  R version 4.1.0 (2021-05-18)
##  os       Windows 10 x64              
##  system   x86_64, mingw32             
##  ui       RTerm                       
##  language (EN)                        
##  collate  English_United States.1252  
##  ctype    English_United States.1252  
##  tz       Europe/Paris                
##  date     2021-07-15                  
## 
## - Packages -------------------------------------------------------------------
##  package     * version date       lib source        
##  assertthat    0.2.1   2019-03-21 [1] CRAN (R 4.1.0)
##  cachem        1.0.5   2021-05-15 [1] CRAN (R 4.1.0)
##  callr         3.7.0   2021-04-20 [1] CRAN (R 4.1.0)
##  cli           2.5.0   2021-04-26 [1] CRAN (R 4.1.0)
##  colorspace    2.0-2   2021-06-24 [1] CRAN (R 4.1.0)
##  cowplot     * 1.1.1   2020-12-30 [1] CRAN (R 4.1.0)
##  crayon        1.4.1   2021-02-08 [1] CRAN (R 4.1.0)
##  DBI           1.1.1   2021-01-15 [1] CRAN (R 4.1.0)
##  desc          1.3.0   2021-03-05 [1] CRAN (R 4.1.0)
##  devtools      2.4.2   2021-06-07 [1] CRAN (R 4.1.0)
##  digest        0.6.27  2020-10-24 [1] CRAN (R 4.1.0)
##  dplyr         1.0.7   2021-06-18 [1] CRAN (R 4.1.0)
##  ellipsis      0.3.2   2021-04-29 [1] CRAN (R 4.1.0)
##  evaluate      0.14    2019-05-28 [1] CRAN (R 4.1.0)
##  fansi         0.5.0   2021-05-25 [1] CRAN (R 4.1.0)
##  farver        2.1.0   2021-02-28 [1] CRAN (R 4.1.0)
##  fastmap       1.1.0   2021-01-25 [1] CRAN (R 4.1.0)
##  fs            1.5.0   2020-07-31 [1] CRAN (R 4.1.0)
##  generics      0.1.0   2020-10-31 [1] CRAN (R 4.1.0)
##  ggplot2     * 3.3.5   2021-06-25 [1] CRAN (R 4.1.0)
##  glue          1.4.2   2020-08-27 [1] CRAN (R 4.1.0)
##  gtable        0.3.0   2019-03-25 [1] CRAN (R 4.1.0)
##  highr         0.9     2021-04-16 [1] CRAN (R 4.1.0)
##  htmltools     0.5.1.1 2021-01-22 [1] CRAN (R 4.1.0)
##  knitr         1.33    2021-04-24 [1] CRAN (R 4.1.0)
##  labeling      0.4.2   2020-10-20 [1] CRAN (R 4.1.0)
##  lifecycle     1.0.0   2021-02-15 [1] CRAN (R 4.1.0)
##  magrittr      2.0.1   2020-11-17 [1] CRAN (R 4.1.0)
##  memoise       2.0.0   2021-01-26 [1] CRAN (R 4.1.0)
##  munsell       0.5.0   2018-06-12 [1] CRAN (R 4.1.0)
##  pillar        1.6.1   2021-05-16 [1] CRAN (R 4.1.0)
##  pkgbuild      1.2.0   2020-12-15 [1] CRAN (R 4.1.0)
##  pkgconfig     2.0.3   2019-09-22 [1] CRAN (R 4.1.0)
##  pkgload       1.2.1   2021-04-06 [1] CRAN (R 4.1.0)
##  prettyunits   1.1.1   2020-01-24 [1] CRAN (R 4.1.0)
##  processx      3.5.2   2021-04-30 [1] CRAN (R 4.1.0)
##  ps            1.6.0   2021-02-28 [1] CRAN (R 4.1.0)
##  purrr         0.3.4   2020-04-17 [1] CRAN (R 4.1.0)
##  R6            2.5.0   2020-10-28 [1] CRAN (R 4.1.0)
##  remotes       2.4.0   2021-06-02 [1] CRAN (R 4.1.0)
##  rlang         0.4.11  2021-04-30 [1] CRAN (R 4.1.0)
##  rmarkdown     2.9     2021-06-15 [1] CRAN (R 4.1.0)
##  rprojroot     2.0.2   2020-11-15 [1] CRAN (R 4.1.0)
##  scales        1.1.1   2020-05-11 [1] CRAN (R 4.1.0)
##  sessioninfo   1.1.1   2018-11-05 [1] CRAN (R 4.1.0)
##  stringi       1.6.1   2021-05-10 [1] CRAN (R 4.1.0)
##  stringr       1.4.0   2019-02-10 [1] CRAN (R 4.1.0)
##  svglite       2.0.0   2021-02-20 [1] CRAN (R 4.1.0)
##  systemfonts   1.0.2   2021-05-11 [1] CRAN (R 4.1.0)
##  testthat      3.0.3   2021-06-16 [1] CRAN (R 4.1.0)
##  tibble        3.1.2   2021-05-16 [1] CRAN (R 4.1.0)
##  tidyselect    1.1.1   2021-04-30 [1] CRAN (R 4.1.0)
##  usethis       2.0.1   2021-02-10 [1] CRAN (R 4.1.0)
##  utf8          1.2.1   2021-03-12 [1] CRAN (R 4.1.0)
##  vctrs         0.3.8   2021-04-29 [1] CRAN (R 4.1.0)
##  withr         2.4.2   2021-04-18 [1] CRAN (R 4.1.0)
##  xfun          0.24    2021-06-15 [1] CRAN (R 4.1.0)
##  yaml          2.2.1   2020-02-01 [1] CRAN (R 4.1.0)
## 
## [1] C:/Users/Ben Field/Documents/R/win-library/4.1
## [2] C:/Program Files/R/R-4.1.0/library
```

#### 6. Bibliopraphy

1. PANTHER version 16: a revised family classification, tree-based classification tool, enhancer regions and extensive API. Huaiyu Mi, Dustin Ebert, Anushya Muruganujan, Caitlin Mills, Laurent-Philippe Albou, Tremayne Mushayamaha and Paul D Thomas . Nucl. Acids Res. (2020) doi: 10.1093/nar/gkaa1106s.
2. Supek F, Bošnjak M, Škunca N, Šmuc T. “REVIGO summarizes and visualizes long lists of Gene Ontology terms” .PLoS ONE 2011. doi:10.1371/journal.pone.0021800
3. A Simple Protocol for Informative Visualization of Enriched Gene Ontology Terms. T. Bonnot, MB. Gillard and DH. Nagel. Bio-101: e3429. DOI:10.21769/BioProtoc.3429
4. R Core Team. 2020. R: A Language and Environment for Statistical Computing. Vienna, Austria: R Foundation for Statistical Computing. https://www.R-project.org/.
5. Wickham, Hadley. 2016. Ggplot2: Elegant Graphics for Data Analysis. Springer-Verlag New York. https://ggplot2.tidyverse.org.
6. Wilke, Claus O. 2020. Cowplot: Streamlined Plot Theme and Plot Annotations for ’Ggplot2’. https://CRAN.R-project.org/package=cowplot.
7. Script prepare\_gene\_onthology.pl. Terese M. et Lecampion C. Github
