## Supplementary material for "A ppGpp-mediated brake on photosynthesis is required for acclimation to nitrogen limitation in Arabidopsis": R scripts: script_organellar_expression.html

Plot organellar transcript levels


### Plot organellar transcript levels

#### ----

#### 1. Introduction

The analysis reported here is part of the manuscript Romand et al. 2021. The script plots RNAseq data for organellar transcript abundance.

Working directory: C:/Users/Ben/Desktop/AMUBOX/Shared/ppGpp\_nitrogen\_starvation/Supp data sets/R scripts/R markdown organelle expression Fig\_5

##### Setup R packages

```
if (!require(magrittr)) { install.packages("magrittr", repos = "http://cran.us.r-project.org") }
if (!require(dplyr)) { install.packages("dplyr", repos = "http://cran.us.r-project.org") }
if (!require(knitr)) { install.packages("knitr", repos = "http://cran.us.r-project.org") }
if (!require(ggplot2)) { install.packages("ggplot2", repos = "http://cran.us.r-project.org") }

library(knitr)
library(ggplot2)
library(magrittr)
library(dplyr)
library(readxl)
```

#### 2. Data

Data is imported from an excel file containing a column of gene ids `id`, a column of log transformed fold change `logFC` values, a column of adjusted p values `padj`, and a column of different treatments or comparisons `treatment`.

```
 data1 <- read_excel("organelle_chloro.xlsx")

 # Add column convert pvalues into two groups- significant and not
 
 data1<-data1%>%mutate(groups = cut(padj, breaks = c(-Inf,0.05,Inf), labels=c("Sig","Not Sig")))
 
 # Set colours
 
 my_colours<- c("dark green","Purple", "Black")
```

#### 3. Plot Graph

Graphs are now plotted for different comparisons. Non significant points are shown, but with higher transparency.

```
 p<-ggplot(data=data1, aes(x=id, y=logFC, color=treatment, alpha=groups))+
   geom_point(aes(y=logFC))+
   scale_alpha_discrete(range = c(0.8, 0.2))+
   scale_color_manual(values = my_colours)+
   geom_hline(yintercept = 0)+
   scale_x_discrete(labels = NULL, breaks = NULL) + labs(x = "")+
   facet_wrap(~treatment, strip.position = "bottom", scales = "free_x", ncol=2) +
   theme_classic() + 
   theme(panel.spacing = unit(0, "lines"), 
         strip.background = element_blank(),
         strip.placement = "outside")+
   coord_cartesian(clip = 'off') +
   theme_classic()
 print(p)
```

```
 # outputs pdf and svg files with defined size
  # ggsave("my_ggplot5.pdf", width=7, height=6)
  # ggsave("my_ggplot5.svg", width=7, height=6)
```

Now the same analysis is performed on mitochondrial transcript levels.

```
data2 <- read_excel("organelle_mito.xlsx")

 data2<-data2%>%mutate(groups = cut(padj, breaks = c(-Inf,0.05,Inf), labels=c("Sig","Not Sig")))

 p<-ggplot(data=data2, aes(x=id, y=logFC, color=treatment, alpha=groups))+
    geom_point(aes(y=logFC))+
    scale_alpha_discrete(range = c(0.8, 0.2))+
    scale_color_manual(values = my_colours)+
    geom_hline(yintercept = 0)+
    scale_x_discrete(labels = NULL, breaks = NULL) + labs(x = "")+
    facet_wrap(~treatment, strip.position = "bottom", scales = "free_x", ncol=2) +
    theme_classic() + 
    theme(panel.spacing = unit(0, "lines"), 
          strip.background = element_blank(),
          strip.placement = "outside")+
    coord_cartesian(clip = 'off') +
    theme_classic()
 print(p)
```

```
 # outputs pdf and svg files with defined size
  # ggsave("mito_ggplot5.pdf", width=7, height=6)
  # ggsave("mito_ggplot5.svg", width=7, height=6)
```

#### 4. R session information

```
InfoSession <- devtools::session_info()
print(InfoSession)
```

```
## - Session info ---------------------------------------------------------------
##  setting  value                       
##  version  R version 4.1.0 (2021-05-18)
##  os       Windows 10 x64              
##  system   x86_64, mingw32             
##  ui       RTerm                       
##  language (EN)                        
##  collate  English_United Kingdom.1252 
##  ctype    English_United Kingdom.1252 
##  tz       Europe/Paris                
##  date     2021-07-15                  
## 
## - Packages -------------------------------------------------------------------
##  package     * version date       lib source        
##  assertthat    0.2.1   2019-03-21 [1] CRAN (R 4.1.0)
##  cachem        1.0.5   2021-05-15 [1] CRAN (R 4.1.0)
##  callr         3.7.0   2021-04-20 [1] CRAN (R 4.1.0)
##  cellranger    1.1.0   2016-07-27 [1] CRAN (R 4.1.0)
##  cli           2.5.0   2021-04-26 [1] CRAN (R 4.1.0)
##  colorspace    2.0-1   2021-05-04 [1] CRAN (R 4.1.0)
##  crayon        1.4.1   2021-02-08 [1] CRAN (R 4.1.0)
##  DBI           1.1.1   2021-01-15 [1] CRAN (R 4.1.0)
##  desc          1.3.0   2021-03-05 [1] CRAN (R 4.1.0)
##  devtools      2.4.2   2021-06-07 [1] CRAN (R 4.1.0)
##  digest        0.6.27  2020-10-24 [1] CRAN (R 4.1.0)
##  dplyr       * 1.0.7   2021-06-18 [1] CRAN (R 4.1.0)
##  ellipsis      0.3.2   2021-04-29 [1] CRAN (R 4.1.0)
##  evaluate      0.14    2019-05-28 [1] CRAN (R 4.1.0)
##  fansi         0.5.0   2021-05-25 [1] CRAN (R 4.1.0)
##  farver        2.1.0   2021-02-28 [1] CRAN (R 4.1.0)
##  fastmap       1.1.0   2021-01-25 [1] CRAN (R 4.1.0)
##  fs            1.5.0   2020-07-31 [1] CRAN (R 4.1.0)
##  generics      0.1.0   2020-10-31 [1] CRAN (R 4.1.0)
##  ggplot2     * 3.3.4   2021-06-16 [1] CRAN (R 4.1.0)
##  glue          1.4.2   2020-08-27 [1] CRAN (R 4.1.0)
##  gtable        0.3.0   2019-03-25 [1] CRAN (R 4.1.0)
##  highr         0.9     2021-04-16 [1] CRAN (R 4.1.0)
##  htmltools     0.5.1.1 2021-01-22 [1] CRAN (R 4.1.0)
##  knitr       * 1.33    2021-04-24 [1] CRAN (R 4.1.0)
##  labeling      0.4.2   2020-10-20 [1] CRAN (R 4.1.0)
##  lifecycle     1.0.0   2021-02-15 [1] CRAN (R 4.1.0)
##  magrittr    * 2.0.1   2020-11-17 [1] CRAN (R 4.1.0)
##  memoise       2.0.0   2021-01-26 [1] CRAN (R 4.1.0)
##  munsell       0.5.0   2018-06-12 [1] CRAN (R 4.1.0)
##  pillar        1.6.1   2021-05-16 [1] CRAN (R 4.1.0)
##  pkgbuild      1.2.0   2020-12-15 [1] CRAN (R 4.1.0)
##  pkgconfig     2.0.3   2019-09-22 [1] CRAN (R 4.1.0)
##  pkgload       1.2.1   2021-04-06 [1] CRAN (R 4.1.0)
##  prettyunits   1.1.1   2020-01-24 [1] CRAN (R 4.1.0)
##  processx      3.5.2   2021-04-30 [1] CRAN (R 4.1.0)
##  ps            1.6.0   2021-02-28 [1] CRAN (R 4.1.0)
##  purrr         0.3.4   2020-04-17 [1] CRAN (R 4.1.0)
##  R6            2.5.0   2020-10-28 [1] CRAN (R 4.1.0)
##  Rcpp          1.0.6   2021-01-15 [1] CRAN (R 4.1.0)
##  readxl      * 1.3.1   2019-03-13 [1] CRAN (R 4.1.0)
##  remotes       2.4.0   2021-06-02 [1] CRAN (R 4.1.0)
##  rlang         0.4.11  2021-04-30 [1] CRAN (R 4.1.0)
##  rmarkdown     2.9     2021-06-15 [1] CRAN (R 4.1.0)
##  rprojroot     2.0.2   2020-11-15 [1] CRAN (R 4.1.0)
##  scales        1.1.1   2020-05-11 [1] CRAN (R 4.1.0)
##  sessioninfo   1.1.1   2018-11-05 [1] CRAN (R 4.1.0)
##  stringi       1.6.1   2021-05-10 [1] CRAN (R 4.1.0)
##  stringr       1.4.0   2019-02-10 [1] CRAN (R 4.1.0)
##  testthat      3.0.3   2021-06-16 [1] CRAN (R 4.1.0)
##  tibble        3.1.2   2021-05-16 [1] CRAN (R 4.1.0)
##  tidyselect    1.1.1   2021-04-30 [1] CRAN (R 4.1.0)
##  usethis       2.0.1   2021-02-10 [1] CRAN (R 4.1.0)
##  utf8          1.2.1   2021-03-12 [1] CRAN (R 4.1.0)
##  vctrs         0.3.8   2021-04-29 [1] CRAN (R 4.1.0)
##  withr         2.4.2   2021-04-18 [1] CRAN (R 4.1.0)
##  xfun          0.24    2021-06-15 [1] CRAN (R 4.1.0)
##  yaml          2.2.1   2020-02-01 [1] CRAN (R 4.1.0)
## 
## [1] C:/Users/Ben/Documents/R/win-library/4.1
## [2] C:/Program Files/R/R-4.1.0/library
```

#### 5. Citations

1. Bache, Stefan Milton and Wickham, Hadley (2020). magrittr: A Forward-Pipe Operator for R. R package version 2.0.1. https://CRAN.R-project.org/package=magrittr
2. R Core Team. 2020. R: A Language and Environment for Statistical Computing. Vienna, Austria: R Foundation for Statistical Computing. https://www.R-project.org/.
3. Wickham, Hadley. 2016. Ggplot2: Elegant Graphics for Data Analysis. Springer-Verlag New York. https://ggplot2.tidyverse.org.
4. Wickham, Hadley, and Jennifer Bryan. 2019. Readxl: Read Excel Files. https://CRAN.R-project.org/package=readxl.
5. Wickham, Hadley, Romain François, Lionel Henry, and Kirill Müller. 2021. Dplyr: A Grammar of Data Manipulation. https://CRAN.R-project.org/package=dplyr.
