## Supplementary material for "A ppGpp-mediated brake on photosynthesis is required for acclimation to nitrogen limitation in Arabidopsis": R scripts: script_prop_test.html

Proportion test and CI calculation


### Proportion test and CI calculation

#### ----

#### 1. Introduction

The analysis reported here is part of the manuscript Romand et al. 2021. The script performs statistical analysis on categorical/nominal data: comparison of dead cotyledon counts between different plant lines grown under N limiting conditions.

The analysis requires the rstatix package and the rcompanion package.

###### Setup R packages

```
if (!require(rstatix)) { install.packages("rstatix", repos = "http://cran.us.r-project.org") }
if (!require(rcompanion)) { install.packages("rcompanion", repos = "http://cran.us.r-project.org") }
if (!require(knitr)) { install.packages("knitr", repos = "http://cran.us.r-project.org") }
if (!require(tidyverse)) { install.packages("tidyverse", repos = "http://cran.us.r-project.org") }
if (!require(devtools)) { install.packages("devtools", repos = "http://cran.us.r-project.org") }


library(rstatix)
library(rcompanion)
library(knitr)
library(tidyverse)
library(devtools)
```

#### 2. Data

Data is inputted manually into a table. In this case it is the summed counts from three biological replicates of the same experiment. Dead is for plants with dead cotyledons, and all is the total number of plants evaluated.

```
# Create data matrix for analysis

xtab <- matrix(c(13, 280, 51, 234, 17, 281, 8, 287, 55, 240 ), nrow = 5, ncol = 2, byrow = TRUE,
               dimnames = list(c("Col", "OXRSH1", "qrt", "rsh1-1","QM"),
                               c("dead", "all")))

kable(xtab)
```

|  | dead | all |
| --- | --- | --- |
| Col | 13 | 280 |
| OXRSH1 | 51 | 234 |
| qrt | 17 | 281 |
| rsh1-1 | 8 | 287 |
| QM | 55 | 240 |

#### 3. Data analysis

A proportion test is performed using the proportion test (prop\_test) from the rstatix package.

```
table<-prop_test(xtab, conf.level = 0.95, detailed=TRUE, correct=TRUE)
kable(table)
```

| n | n1 | n2 | n3 | n4 | n5 | estimate1 | estimate2 | estimate3 | estimate4 | estimate5 | statistic | p | df | method | alternative | p.signif |
| --- | --- | --- | --- | --- | --- | --- | --- | --- | --- | --- | --- | --- | --- | --- | --- | --- |
| 1466 | 293 | 285 | 298 | 295 | 295 | 0.0443686 | 0.1789474 | 0.057047 | 0.0271186 | 0.1864407 | 79.02052 | 0 | 4 | Prop test | two.sided | \*\*\*\* |

Now a post-hoc pairwaise Fisher test is performed to identify which differences are signficant using the rcompanion package. See `https://rcompanion.org/handbook/H_04.html` for more information.

```
PT = pairwiseNominalIndependence(xtab,
                                 compare = "row",
                                 fisher  = TRUE,
                                 gtest   = FALSE,
                                 chisq   = FALSE,
                                 method  = "fdr",  # False Discovery Rate, see ?p.adjust for options
                                 digits  = 3)

# Add a significance column

PT<-PT %>%
  mutate(Significance = case_when(
    p.adj.Fisher <=0.01 ~ "**",
    p.adj.Fisher <=0.05 ~ "*",
    p.adj.Fisher >=0.05 ~ "ns",
    ))

kable(PT)
```

| Comparison | p.Fisher | p.adj.Fisher | Significance |
| --- | --- | --- | --- |
| Col : OXRSH1 | 2.00e-07 | 6.00e-07 | \*\* |
| Col : qrt | 5.75e-01 | 6.39e-01 | ns |
| Col : rsh1-1 | 2.76e-01 | 3.45e-01 | ns |
| Col : QM | 1.00e-07 | 2.00e-07 | \*\* |
| OXRSH1 : qrt | 4.30e-06 | 7.20e-06 | \*\* |
| OXRSH1 : rsh1-1 | 0.00e+00 | 0.00e+00 | \*\* |
| OXRSH1 : QM | 8.31e-01 | 8.31e-01 | ns |
| qrt : rsh1-1 | 1.00e-01 | 1.43e-01 | ns |
| qrt : QM | 1.20e-06 | 2.30e-06 | \*\* |
| rsh1-1 : QM | 0.00e+00 | 0.00e+00 | \*\* |

Next confidence intervals are calculated using the binomial test.

```
data<-as.data.frame(xtab)

# Add columns to show line names and proportion dead 

data$Lines <- row.names(data)

data$dead_proportion = data$dead / data$all

data<-data[,c(3,1,2,4)]

# Perform binom test and extract confidence intervals

data<-data %>% 
    rowwise() %>% 
    mutate(CI= list(enframe(binom.test(dead, all, alternative =  "two.sided", conf.level = 0.95)$conf.int))) %>% 
    unnest(CI) %>% 
    spread(name, value) %>% 
    rename("LCB" = "1", "UCB" = "2")


kable(data, digits= 3)
```

| Lines | dead | all | dead\_proportion | LCB | UCB |
| --- | --- | --- | --- | --- | --- |
| Col | 13 | 280 | 0.046 | 0.025 | 0.078 |
| OXRSH1 | 51 | 234 | 0.218 | 0.167 | 0.276 |
| QM | 55 | 240 | 0.229 | 0.178 | 0.288 |
| qrt | 17 | 281 | 0.060 | 0.036 | 0.095 |
| rsh1-1 | 8 | 287 | 0.028 | 0.012 | 0.054 |

#### 4. R session

```
InfoSession <- devtools::session_info()
print(InfoSession)
```

```
## - Session info ---------------------------------------------------------------
##  setting  value                       
##  version  R version 4.1.0 (2021-05-18)
##  os       Windows 10 x64              
##  system   x86_64, mingw32             
##  ui       RTerm                       
##  language (EN)                        
##  collate  English_United States.1252  
##  ctype    English_United States.1252  
##  tz       Europe/Paris                
##  date     2021-06-29                  
## 
## - Packages -------------------------------------------------------------------
##  package      * version date       lib source        
##  abind          1.4-5   2016-07-21 [1] CRAN (R 4.1.0)
##  assertthat     0.2.1   2019-03-21 [1] CRAN (R 4.1.0)
##  backports      1.2.1   2020-12-09 [1] CRAN (R 4.1.0)
##  boot           1.3-28  2021-05-03 [2] CRAN (R 4.1.0)
##  broom          0.7.8   2021-06-24 [1] CRAN (R 4.1.0)
##  cachem         1.0.5   2021-05-15 [1] CRAN (R 4.1.0)
##  callr          3.7.0   2021-04-20 [1] CRAN (R 4.1.0)
##  car            3.0-11  2021-06-27 [1] CRAN (R 4.1.0)
##  carData        3.0-4   2020-05-22 [1] CRAN (R 4.1.0)
##  cellranger     1.1.0   2016-07-27 [1] CRAN (R 4.1.0)
##  class          7.3-19  2021-05-03 [2] CRAN (R 4.1.0)
##  cli            2.5.0   2021-04-26 [1] CRAN (R 4.1.0)
##  codetools      0.2-18  2020-11-04 [2] CRAN (R 4.1.0)
##  coin           1.4-1   2021-02-08 [1] CRAN (R 4.1.0)
##  colorspace     2.0-2   2021-06-24 [1] CRAN (R 4.1.0)
##  crayon         1.4.1   2021-02-08 [1] CRAN (R 4.1.0)
##  curl           4.3.1   2021-04-30 [1] CRAN (R 4.1.0)
##  data.table     1.14.0  2021-02-21 [1] CRAN (R 4.1.0)
##  DBI            1.1.1   2021-01-15 [1] CRAN (R 4.1.0)
##  dbplyr         2.1.1   2021-04-06 [1] CRAN (R 4.1.0)
##  desc           1.3.0   2021-03-05 [1] CRAN (R 4.1.0)
##  DescTools      0.99.42 2021-06-17 [1] CRAN (R 4.1.0)
##  devtools     * 2.4.2   2021-06-07 [1] CRAN (R 4.1.0)
##  digest         0.6.27  2020-10-24 [1] CRAN (R 4.1.0)
##  dplyr        * 1.0.7   2021-06-18 [1] CRAN (R 4.1.0)
##  e1071          1.7-7   2021-05-23 [1] CRAN (R 4.1.0)
##  ellipsis       0.3.2   2021-04-29 [1] CRAN (R 4.1.0)
##  evaluate       0.14    2019-05-28 [1] CRAN (R 4.1.0)
##  Exact          2.1     2020-10-02 [1] CRAN (R 4.1.0)
##  expm           0.999-6 2021-01-13 [1] CRAN (R 4.1.0)
##  fansi          0.5.0   2021-05-25 [1] CRAN (R 4.1.0)
##  fastmap        1.1.0   2021-01-25 [1] CRAN (R 4.1.0)
##  forcats      * 0.5.1   2021-01-27 [1] CRAN (R 4.1.0)
##  foreign        0.8-81  2020-12-22 [2] CRAN (R 4.1.0)
##  fs             1.5.0   2020-07-31 [1] CRAN (R 4.1.0)
##  generics       0.1.0   2020-10-31 [1] CRAN (R 4.1.0)
##  ggplot2      * 3.3.5   2021-06-25 [1] CRAN (R 4.1.0)
##  gld            2.6.2   2020-01-08 [1] CRAN (R 4.1.0)
##  glue           1.4.2   2020-08-27 [1] CRAN (R 4.1.0)
##  gtable         0.3.0   2019-03-25 [1] CRAN (R 4.1.0)
##  haven          2.4.1   2021-04-23 [1] CRAN (R 4.1.0)
##  highr          0.9     2021-04-16 [1] CRAN (R 4.1.0)
##  hms            1.1.0   2021-05-17 [1] CRAN (R 4.1.0)
##  htmltools      0.5.1.1 2021-01-22 [1] CRAN (R 4.1.0)
##  httr           1.4.2   2020-07-20 [1] CRAN (R 4.1.0)
##  jsonlite       1.7.2   2020-12-09 [1] CRAN (R 4.1.0)
##  knitr        * 1.33    2021-04-24 [1] CRAN (R 4.1.0)
##  lattice        0.20-44 2021-05-02 [2] CRAN (R 4.1.0)
##  libcoin        1.0-8   2021-02-08 [1] CRAN (R 4.1.0)
##  lifecycle      1.0.0   2021-02-15 [1] CRAN (R 4.1.0)
##  lmom           2.8     2019-03-12 [1] CRAN (R 4.1.0)
##  lmtest         0.9-38  2020-09-09 [1] CRAN (R 4.1.0)
##  lubridate      1.7.10  2021-02-26 [1] CRAN (R 4.1.0)
##  magrittr       2.0.1   2020-11-17 [1] CRAN (R 4.1.0)
##  MASS           7.3-54  2021-05-03 [2] CRAN (R 4.1.0)
##  Matrix         1.3-3   2021-05-04 [2] CRAN (R 4.1.0)
##  matrixStats    0.59.0  2021-06-01 [1] CRAN (R 4.1.0)
##  memoise        2.0.0   2021-01-26 [1] CRAN (R 4.1.0)
##  modelr         0.1.8   2020-05-19 [1] CRAN (R 4.1.0)
##  modeltools     0.2-23  2020-03-05 [1] CRAN (R 4.1.0)
##  multcomp       1.4-17  2021-04-29 [1] CRAN (R 4.1.0)
##  multcompView   0.1-8   2019-12-19 [1] CRAN (R 4.1.0)
##  munsell        0.5.0   2018-06-12 [1] CRAN (R 4.1.0)
##  mvtnorm        1.1-2   2021-06-07 [1] CRAN (R 4.1.0)
##  nortest        1.0-4   2015-07-30 [1] CRAN (R 4.1.0)
##  openxlsx       4.2.4   2021-06-16 [1] CRAN (R 4.1.0)
##  pillar         1.6.1   2021-05-16 [1] CRAN (R 4.1.0)
##  pkgbuild       1.2.0   2020-12-15 [1] CRAN (R 4.1.0)
##  pkgconfig      2.0.3   2019-09-22 [1] CRAN (R 4.1.0)
##  pkgload        1.2.1   2021-04-06 [1] CRAN (R 4.1.0)
##  plyr           1.8.6   2020-03-03 [1] CRAN (R 4.1.0)
##  prettyunits    1.1.1   2020-01-24 [1] CRAN (R 4.1.0)
##  processx       3.5.2   2021-04-30 [1] CRAN (R 4.1.0)
##  proxy          0.4-26  2021-06-07 [1] CRAN (R 4.1.0)
##  ps             1.6.0   2021-02-28 [1] CRAN (R 4.1.0)
##  purrr        * 0.3.4   2020-04-17 [1] CRAN (R 4.1.0)
##  R6             2.5.0   2020-10-28 [1] CRAN (R 4.1.0)
##  rcompanion   * 2.4.1   2021-05-18 [1] CRAN (R 4.1.0)
##  Rcpp           1.0.6   2021-01-15 [1] CRAN (R 4.1.0)
##  readr        * 1.4.0   2020-10-05 [1] CRAN (R 4.1.0)
##  readxl         1.3.1   2019-03-13 [1] CRAN (R 4.1.0)
##  remotes        2.4.0   2021-06-02 [1] CRAN (R 4.1.0)
##  reprex         2.0.0   2021-04-02 [1] CRAN (R 4.1.0)
##  rio            0.5.27  2021-06-21 [1] CRAN (R 4.1.0)
##  rlang          0.4.11  2021-04-30 [1] CRAN (R 4.1.0)
##  rmarkdown      2.9     2021-06-15 [1] CRAN (R 4.1.0)
##  rootSolve      1.8.2.1 2020-04-27 [1] CRAN (R 4.1.0)
##  rprojroot      2.0.2   2020-11-15 [1] CRAN (R 4.1.0)
##  rstatix      * 0.7.0   2021-02-13 [1] CRAN (R 4.1.0)
##  rstudioapi     0.13    2020-11-12 [1] CRAN (R 4.1.0)
##  rvest          1.0.0   2021-03-09 [1] CRAN (R 4.1.0)
##  sandwich       3.0-1   2021-05-18 [1] CRAN (R 4.1.0)
##  scales         1.1.1   2020-05-11 [1] CRAN (R 4.1.0)
##  sessioninfo    1.1.1   2018-11-05 [1] CRAN (R 4.1.0)
##  stringi        1.6.1   2021-05-10 [1] CRAN (R 4.1.0)
##  stringr      * 1.4.0   2019-02-10 [1] CRAN (R 4.1.0)
##  survival       3.2-11  2021-04-26 [2] CRAN (R 4.1.0)
##  testthat       3.0.3   2021-06-16 [1] CRAN (R 4.1.0)
##  TH.data        1.0-10  2019-01-21 [1] CRAN (R 4.1.0)
##  tibble       * 3.1.2   2021-05-16 [1] CRAN (R 4.1.0)
##  tidyr        * 1.1.3   2021-03-03 [1] CRAN (R 4.1.0)
##  tidyselect     1.1.1   2021-04-30 [1] CRAN (R 4.1.0)
##  tidyverse    * 1.3.1   2021-04-15 [1] CRAN (R 4.1.0)
##  usethis      * 2.0.1   2021-02-10 [1] CRAN (R 4.1.0)
##  utf8           1.2.1   2021-03-12 [1] CRAN (R 4.1.0)
##  vctrs          0.3.8   2021-04-29 [1] CRAN (R 4.1.0)
##  withr          2.4.2   2021-04-18 [1] CRAN (R 4.1.0)
##  xfun           0.24    2021-06-15 [1] CRAN (R 4.1.0)
##  xml2           1.3.2   2020-04-23 [1] CRAN (R 4.1.0)
##  yaml           2.2.1   2020-02-01 [1] CRAN (R 4.1.0)
##  zip            2.2.0   2021-05-31 [1] CRAN (R 4.1.0)
##  zoo            1.8-9   2021-03-09 [1] CRAN (R 4.1.0)
## 
## [1] C:/Users/Ben Field/Documents/R/win-library/4.1
## [2] C:/Program Files/R/R-4.1.0/library
```

#### 5. Citations

1. Kassambara, Alboukadel. 2021. Rstatix: Pipe-Friendly Framework for Basic Statistical Tests. https://CRAN.R-project.org/package=rstatix.
2. Mangiafico, Salvatore. 2021. Rcompanion: Functions to Support Extension Education Program Evaluation. https://CRAN.R-project.org/package=rcompanion.
3. R Core Team. 2020. R: A Language and Environment for Statistical Computing. Vienna, Austria: R Foundation for Statistical Computing. https://www.R-project.org/.
4. Wickham et al., (2019). Welcome to the tidyverse. Journal of Open Source Software, 4(43), 1686, https://doi.org/10.21105/joss.01686
5. Wickham, Hadley, Hester, Jim and Chang, Winston (2021). devtools: Tools to Make Developing R Packages Easier. R package version 2.4.2. https://CRAN.R-project.org/package=devtools
6. Yihui Xie (2021). knitr: A General-Purpose Package for Dynamic Report Generation in R. R package version 1.33.
