## Supplementary figures and images for "A ppGpp-mediated brake on photosynthesis is required for acclimation to nitrogen limitation in Arabidopsis"

### ETRN50.pdf

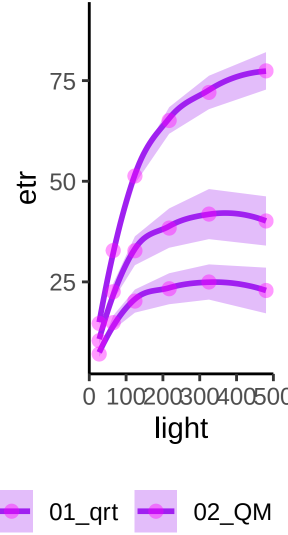

### full77K.pdf

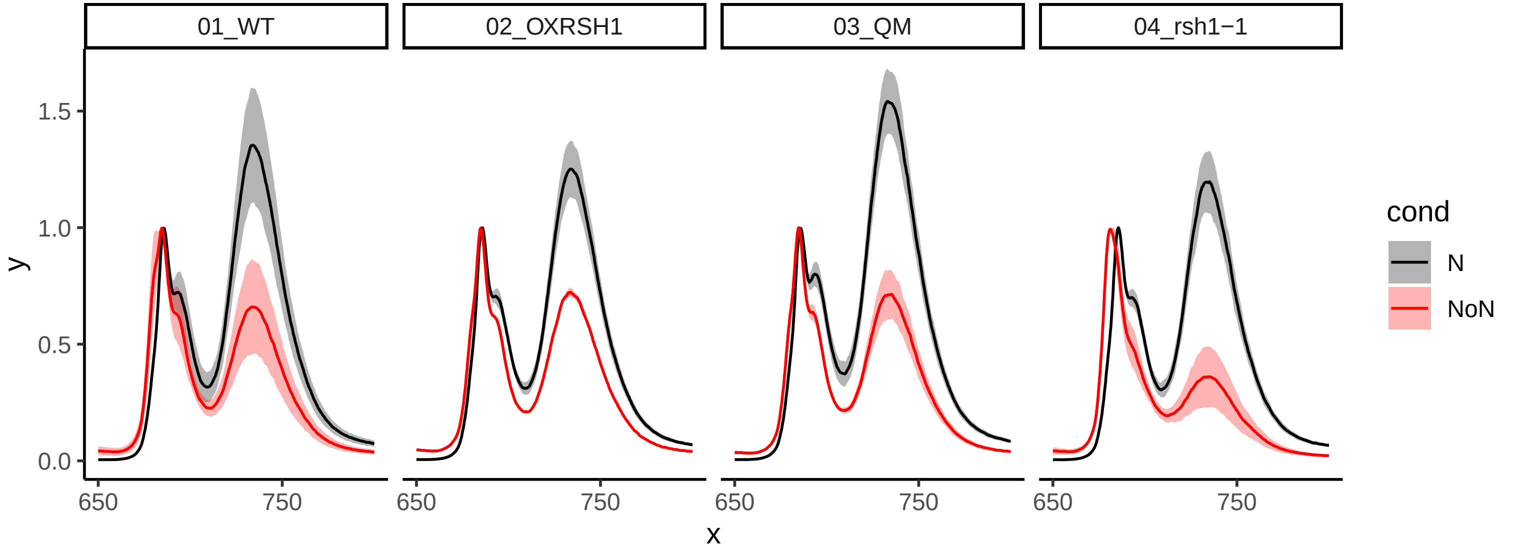

### GO.pdf

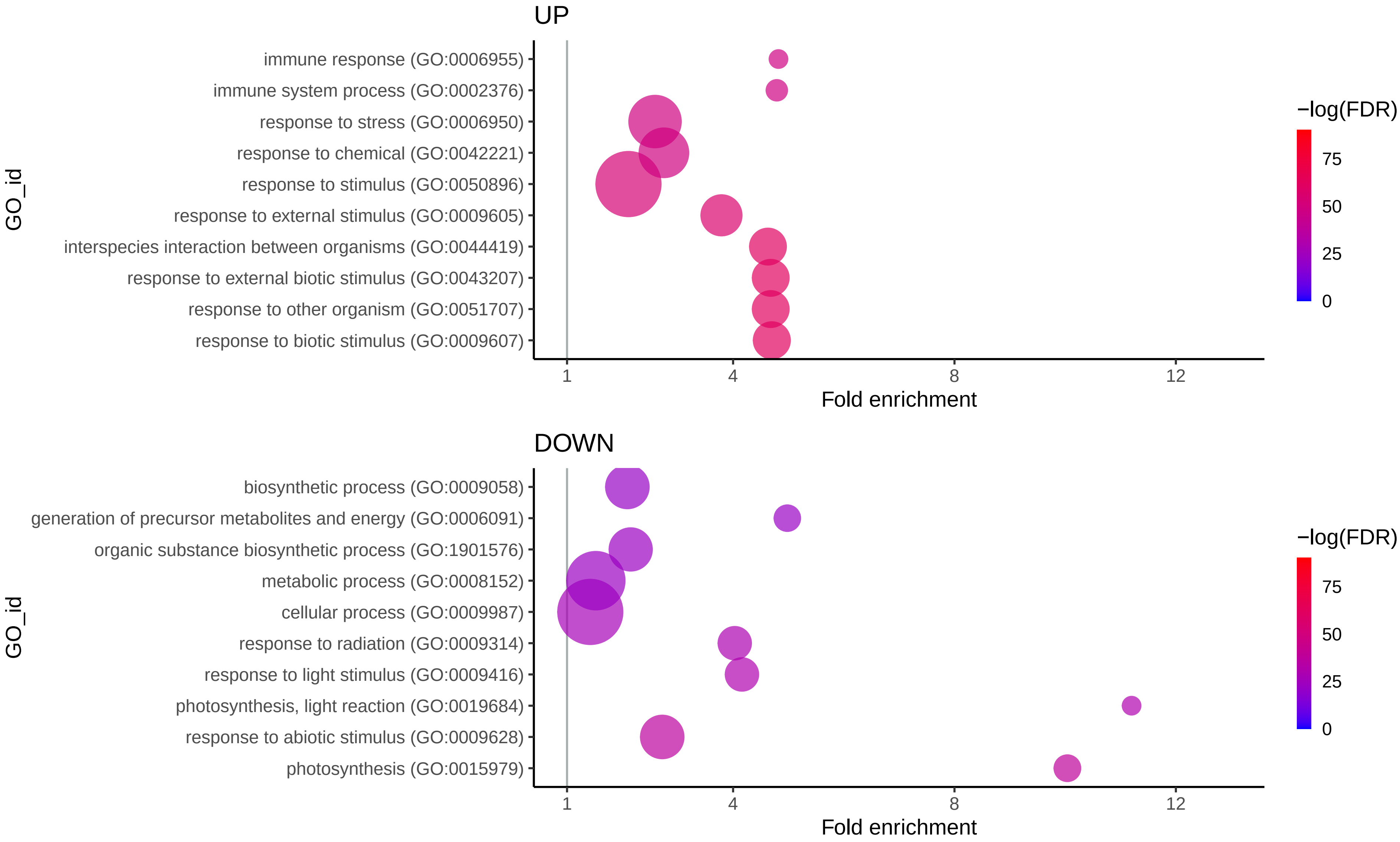

### hGO.pdf

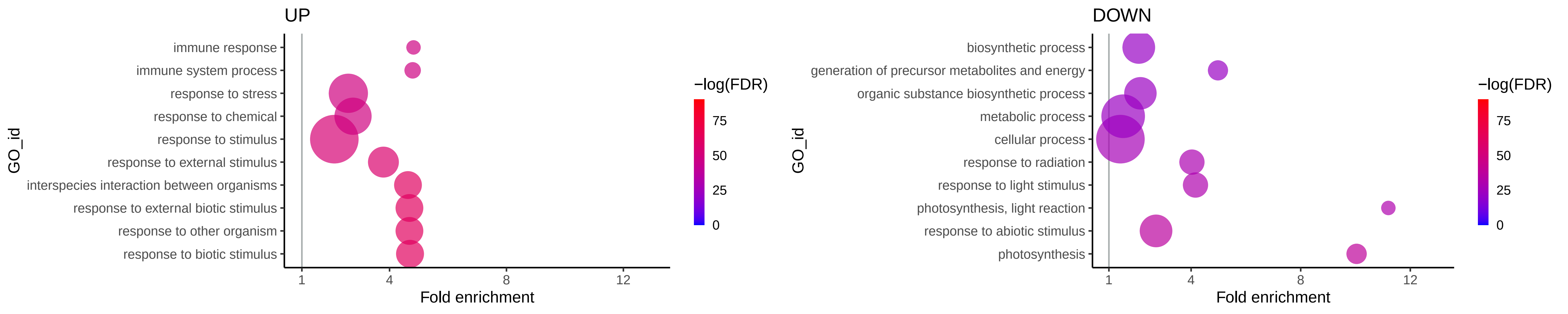

### my_ggplot1.pdf

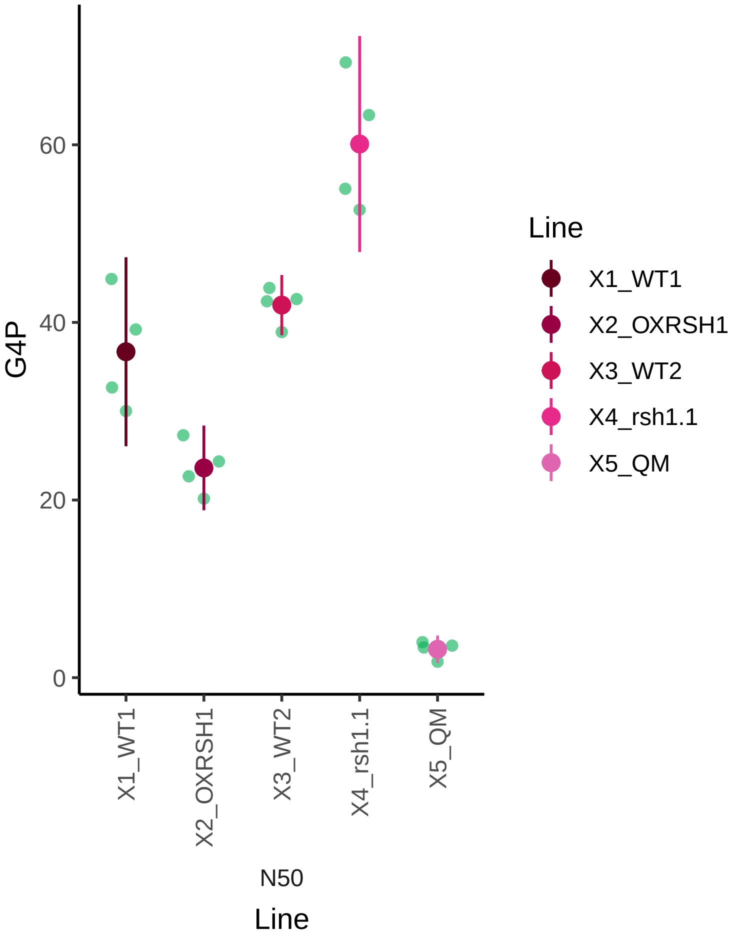

### my_ggplot1.pdf

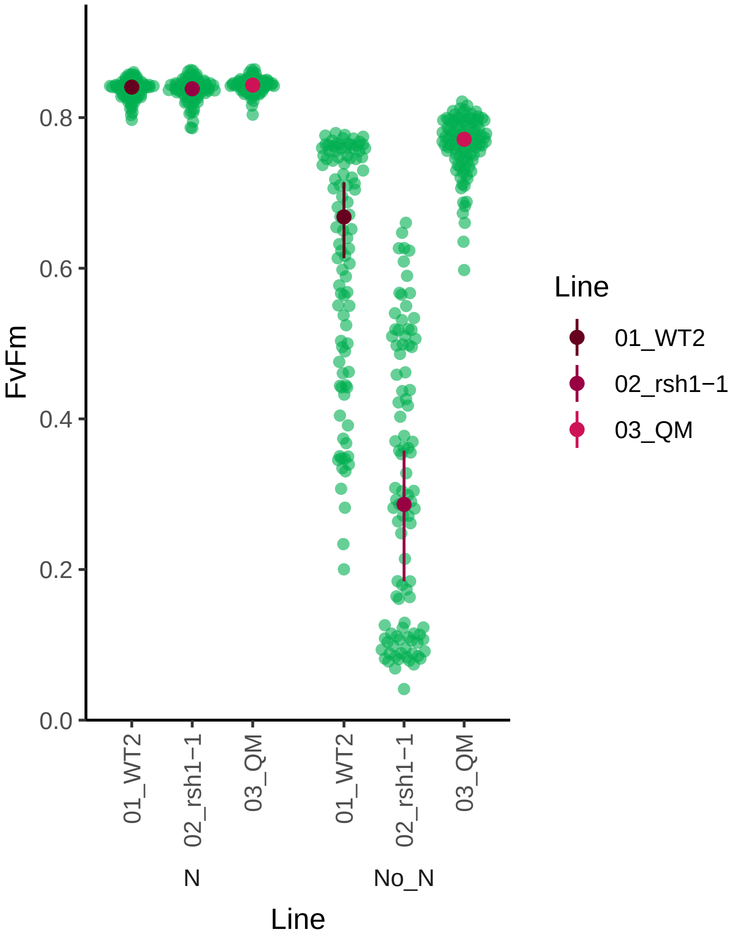

### zoom77K.pdf

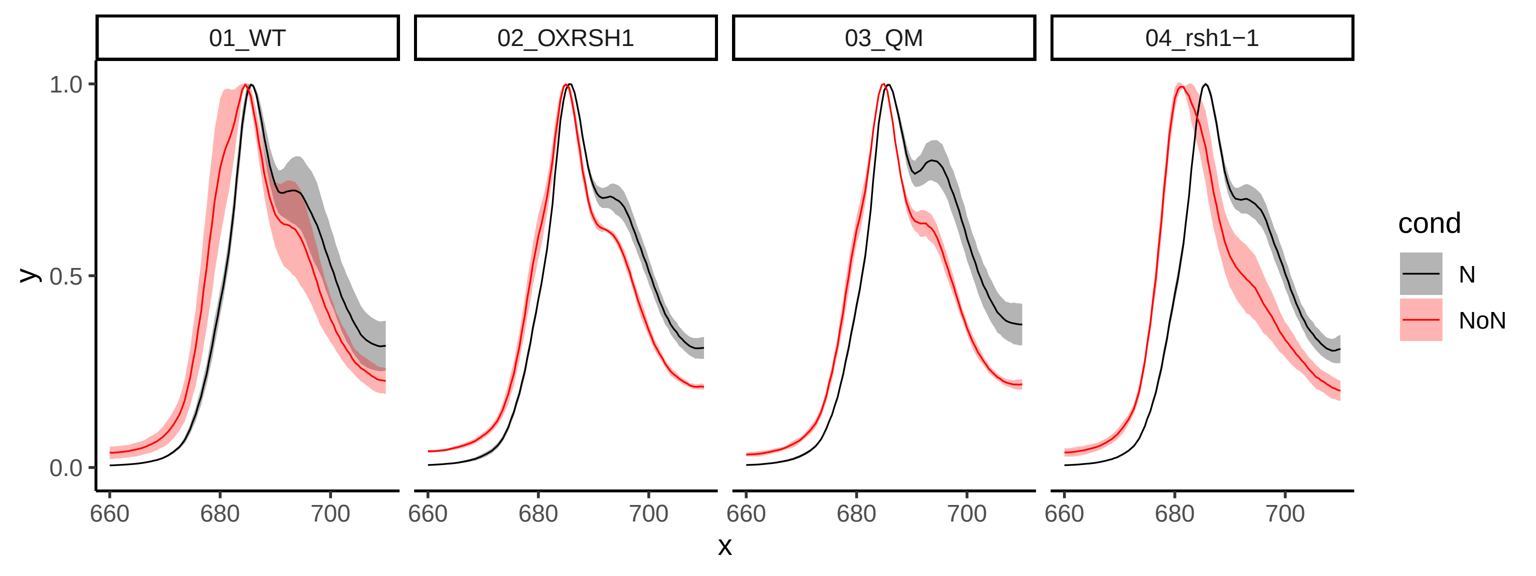
