## Supplementary material for "A ppGpp-mediated brake on photosynthesis is required for acclimation to nitrogen limitation in Arabidopsis": RNAseq analysis reports: 2020_07_22_16 0411 LECAMPION Cecile G4PLAST1_pipeline.html

### Projet G4PLAST1

RNA seq single end.  
2 lignées, WT et OXRSH1  
2 conditions, MS2 et N50

#### Vérification de la qualité des séquences

###### 1- Utilisation de FASTQC (--version 0.11.5)

> Andrews S. (2010). FastQC: a quality control tool for high throughput sequence data. Available online at: http://www.bioinformatics.babraham.ac.uk/projects/fastqc

```
cecile@cecile-vm-linux:~
$ cd '/home/cecile/tmp/FastQC'
```

###### 2- Ouverture d'une interface graphique pour FASTQC

```
cecile@cecile-vm-linux:~/tmp/FastQC
$ ./fastqc
```

#### Alignement contre le transcriptome avec bowtie2

Avec Bowtie2 (--version 2.2.9)

> Langmead B, Salzberg S. Fast gapped-read alignment with Bowtie 2. Nature Methods. 2012, 9:357-359.

On aligne contre Araport en utilisant : `Araport11_cdna_20160703_representative_gene_model.fa`

###### 1- Faire l'index

Dans le dossier `/path_to/Ref_files`

```
$ mkdir Araport11_cdna_20160703_representative_gene_model_bowtie2_index

$ bowtie2-build Araport11_cdna_20160703_representative_gene_model.fa Araport11_cdna_20160703_representative_gene_model_bowtie2_index/At
```

###### 2- Faire l'alignement

Dans le dossier `/path_to/G4PLAST1`

Créer un répertoire pour les alignements

```
$ mkdir bowtie2_align
```

Faire l'alignement

```
$ for F in *.fastq; do echo "=> $F"; bowtie2 --nofw  -p 2 --local -x /home/cecile/partage/Data/ref_files/Araport11_cdna_20160703_representative_gene_model_bowtie2_index/At -U $F | samtools view -bT  /home/cecile/partage/Data/ref_files/Araport11_cdna_20160703_representative_gene_model.fa > ~/partage/F/G4PLAST1/bowtie2_align/${F%.*}.bam; done 2>&1 | tee ~/partage/F/G4PLAST1/bowtie2_align/bowtie.log
```

###### 3- Trier par coordonnées

```
$ mkdir bowtie2_align_sorted
```

Se placer dans le dossier `/path_to/bowtie2_align`

```
$ for F in *.bam; do echo "=> $F"; samtools sort $F > ~/partage/F/G4PLAST1/bowtie2_align_sorted/${F%.*}_sorted.bam; done
```

#### Comptage des reads

Créer un répertoire

```
$ mkdir count
```

Le comptage ne peut pas être réalisé avec HTseq-count dans le cas d'un alignement contre le transcriptome. J'utilise une commande dérivé de celle décrite dans :

> RNA-Seq : revelation of the messengers. Pieterse MJ, Verk MC Van, Hickman R, Wees SCM Van (2013). Trends Plant Sci 18:175–179.

```
$ samtools view -h file.bam | perl -wane 'if (/^\@/) { if (/^\@SQ/) { $trans = $F[1]; $trans =~ s/^SN://; $h{$trans} = 0 } } else { $trans = $F[2]; $h{$trans}++ } END { for $trans (sort keys %h) { print "$trans\t$h{$trans}\n" } }' > ~/partage/F/G4PLAST1/count/file.count
```

#### Etude statistique et génération du fichier d'expression des gènes avec SARTools

> Varet, Hugo and Brillet-Guéguen, Loraine and Coppée, Jean-Yves and Dillies, Marie-Agnès (2016). SARTools: A DESeq2- and EdgeR-Based R Pipeline for Comprehensive Differential Analysis of RNA-Seq Data. In PLoS ONE, 11 (6), pp.

###### 1- Création du fichier `target.txt`

Il est necessaire de créer un fichier contenant des colonnes séparées par des tabulations. Ce fichier permet de décrire les conditions dans lesquelle la manipulation a été faite (nombre de réplicat, date...)

Pour la comparaison OXRSH1\_MS2 vs WT\_MS2

```
label       files                       group   batch
wt_1        WT_MS2_1_sorted.count       WT      day1
wt_2        WT_MS2_2_sorted.count       WT      day2
wt_3        WT_MS2_3_sorted.count       WT      day3
oxrsh1_1    OXRSH1_MS2_1_sorted.count   OXRSH1  day1
oxrsh1_2    OXRSH1_MS2_2_sorted.count   OXRSH1  day2
oxrsh1_3    OXRSH1_MS2_3_sorted.count   OXRSH1  day3
```

On crée le même fichier pour chaque comparaison

###### 2- Utilisation du script `template_script_Edge.r`

Il faut ouvrir une console Rstudio et remplir les paramètres propres à chaque condition :

```
  ################################################################################
  ### R script to compare several conditions with the SARTools and edgeR packages
  ### Hugo Varet
  ### November 28th, 2019
  ### designed to be executed with SARTools 1.7.3
  ################################################################################

  ################################################################################
  ###                parameters: to be modified by the user                    ###
  ################################################################################
rm(list=ls())                                        # remove all the objects from the R session

workDir <- "/home/cecile/partage/F/G4PLAST1/sartools_OXRSH1_MS2_vs_WT_MS2"      # working directory for the R session

projectName <- "G4PLAST1"                         # name of the project
author <- "Cécile"                                # author of the statistical analysis/report

targetFile <- "target_file.txt"                           # path to the design/target file
rawDir <- "/home/cecile/partage/F/G4PLAST1/count"                                      # path to the directory     containing raw counts files
featuresToRemove <- c("alignment_not_unique",        # names of the features to be removed
                  "ambiguous", "no_feature",     # (specific HTSeq-count information and rRNA for example)
                  "not_aligned", "too_low_aQual")# NULL if no feature to remove

varInt <- "group"                                    # factor of interest
condRef <- "WT"                                      # reference biological condition
batch <- "batch"                                        # blocking factor: NULL (default) or "batch" for example

alpha <- 0.05                                        # threshold of statistical significance
pAdjustMethod <- "BH"                                # p-value adjustment method: "BH" (default) or "BY"

cpmCutoff <- 1                                       # counts-per-million cut-off to filter low counts
gene.selection <- "pairwise"                         # selection of the features in MDSPlot
normalizationMethod <- "TMM"                         # normalization method: "TMM" (default), "RLE" (DESeq) or "upperquartile"

colors <- c("#f3c300", "#875692", "#f38400",         # vector of colors of each biological condition on the plots
        "#a1caf1", "#be0032", "#c2b280",
        "#848482", "#008856", "#e68fac",
        "#0067a5", "#f99379", "#604e97")

forceCairoGraph <- FALSE
```

Puis on exécute le script

On obtient, pour chaque condition, un rapport statistique et 3 fichiers `.txt` contenant les valeurs chiffrées de l'expression différentielle des gènes : une liste complète, une liste des gènes réprimés et une liste des gènes activés.
